# The maintenance of genetic polymorphism underlying sexually antagonistic traits

**DOI:** 10.1101/2023.10.10.561678

**Authors:** Ewan Flintham, Vincent Savolainen, Sarah Otto, Max Reuter, Charles Mullon

## Abstract

Selection often favours different trait values in males and females, leading to genetic conflicts between the sexes when traits have a shared genetic basis. Such sexual antagonism has been proposed to maintain genetic polymorphism. However, this notion is based on insights from population genetic models of single loci with fixed fitness effects. It is thus unclear how readily polymorphism emerges from sex-specific selection acting on continuous traits, where fitness effects arise from the genotype-phenotype map and the fitness landscape. Here we model the evolution of a continuous trait that has a shared genetic basis but different optima in males and females, considering a wide variety of genetic architectures and fitness landscapes. For autosomal loci, the long-term maintenance of polymorphism requires strong conflict between males and females that generates uncharacteristic sex-specific fitness patterns. Instead, more plausible sex-specific fitness landscapes typically generate stabilising selection leading to an evolutionarily stable state that consists of a single homozygous genotype. Except for sites tightly linked to the sex determining region, our results indicate that genetic variation due to sexual antagonism should arise only rarely and often be transient, making these signatures challenging to detect in genomic data.

## 1 Introduction

In sexual populations, genetic conflict can arise between males and females when traits that are genetically correlated across the sexes show different optima [1–3]. Here, alleles with similar phenotypic effects in males and females can be favoured in one sex but disfavoured in the other [1, 4–9]. Such “sexual antagonism” [9–11] is thought to have wide ranging ecological and evolutionary implications, from driving a fitness load across the sexes [1, 12] to shaping patterns of genetic variation within, and even between, species [1, 10, 13] ([14–16] for empirical examples).

The potential influence of sexually antagonistic selection on genetic variation has been highlighted by multiple population genetic models [17–36]. One salient point from this theory is that sexual antagonism can maintain a balanced genetic polymorphism for two alleles at a locus either when selection is strong and symmetrical between the sexes, or when allelic dominance is sex-specific such that alleles are more dominant when beneficial and more recessive when deleterious (“sex-specific dominance reversal” [18, 23, 30, 37]). Models of trait evolution [10, 37] have found that such dominance relationships frequently emerge during the course of sex-specific adaptation in the weak-mutation strong-selection limit (i.e. such that a maximum of one di-allelic locus segregates at any given time) if fitness landscapes decline slowly away from phenotypic optima (e.g. Gaussian fitness landscapes). This has led to the suggestion that sexually antagonistic balancing selection is a common outcome of sex-specific selection [10, 13, 30, 37–41], including selection acting on traits influenced by multiple segregating loci or by loci at which alleles with a range of phenotypic effects may arise over time [2, 42, 43] (i.e., multi-allelic or continuous traits [44, 45]).

In contrast to alleles at a single di-allelic locus, however, the fitness consequences of alleles underlying polygenic or continuous traits are not straightforward, being determined by the genotype-phenotype map and the male and female fitness landscapes [45]. This calls into question whether insights into the maintenance of genetic variation from single-locus theory readily extend to traits with a more complex basis. For example, multi-locus population genetic modelling suggests that variation is reduced across loci when males and females have different genotype-phenotype maps but selection is stabilising for the same trait value in both sexes [46]. Quantitative genetic models investigating sexspecific adaptation in complex traits, meanwhile, provide little information with regards to the maintenance of genetic variation as variation is assumed constant through time (i.e., that traits are determined by many loci of small effects such that cross-sex genetic correlations are constant [12, 47, 48]). Bridging population and quantitative genetic models, recent simulation work has shown how sexual dimorphism in a polygenic trait can evolve outside of the weak-mutation strong-selection limit, but does not consider the potential for unresolved sexual antagonism to maintain variation [11].

Here, we study the maintenance of genetic variation for a continuous trait subject to male- and female-specific fitness landscapes with a shared genetic basis across the sexes, such that genetic conflict emerges. Using a combination of mathematical analyses and multi-locus simulations, we show that sexually antagonistic selection should rarely drive polymorphism when traits are polygenic or where loci undergo continual mutation. Rather, our results suggest that sexually antagonistic selection will typically diminish genetic variation, unless traits are determined by very few loci experiencing strong genetic constraints (e.g., where very few alleles can arise through mutation). Our findings further imply that detecting sexually antagonistic variation from genomic data of natural populations will likely remain challenging because such polymorphism should be transient.

## 2 Model

### 2.1 Life-cycle, traits, and sex-specific fitness landscape

We consider a diploid population of large and constant size that is composed of equal numbers of adult males and females. At each generation, adult males and females produce large numbers of gametes that fuse randomly to produce zygotes. These zygotes become male and female juveniles with equal probability (so the sex ratio at birth is unbiased). Adults die and juveniles compete randomly within each sex to become the male and female adults of the next generation.

Each individual expresses a continuous trait, *z* ∈ **ℝ**, that influences their fitness. We consider a scenario whereby this trait affects the number of gametes produced (i.e., fecundity), but trait effects on fitness could equivalently be through survival and/or competition (e.g., for breeding spots) within sexes (as in [10, 11, 18, 19, 49, 50]). Male and female fecundity may depend on trait *z* differently, and we denote the fecundity of a male with trait *z* as *w*_m_(*z*) and fecundity of a female with trait *z* as *w*_f_(*z*). The functions *w*_m_(*z*) and *w*_f_(*z*) thus characterise the male and female fitness landscape, respectively. We focus on a range of trait values over which there is conflict between male and female trait expression, i.e., a range of trait values over which male and female fecundity change in opposing directions with *z* (e.g., *w ′_m_*(*z*) *<* 0 and *w ′_f_*(*z*) *>* 0 where ′ denotes derivative). In Box I, we present flexible forms of *w*_m_(*z*) and *w*_f_(*z*) based on power functions that are respectively centered around a male (−*θ*) and a female (*θ*) optimum. We will use these functions for our analyses where necessary, in particular for individual-based simulations (we also consider Gaussian functions in the Appendix, where we obtain equivalent results to those presented in the main text).

#### Box 1. Fecundity power functions

To illustrate how different fitness landscapes influence outcomes of sexual antagonism, we introduce the power functions

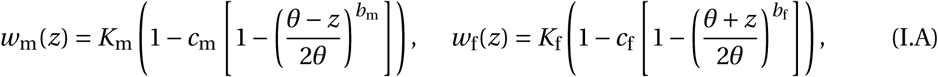

where *K*_m_ and *K*_f_ are the maximum number of gametes produced by a male and a female (both assumed to be large). According to eq. (I.A), *w*_m_(*z*) decreases from *K*_m_ to (1 − *c*_m_)*K*_m_ as *z* increases from *θ*, whereas *w*_f_(*z*) decreases from *K*_f_ to *K*_f_(1 *− c*_f_) as *z* decreases from *θ* (Fig. 1). Here, we assume that *z* is bounded to *θ* ≤ z ≤ µ so that our analyses are focused on the trait space between the male and female optima where selection is sexually antagonistic. The coefficients 0 *≤ c*_m_ *≤* 1 and 0 *≤ c*_f_ *≤* 1 tune the steepness of the fitness decline in each sex and so the strength of the trade-off over male and female fecundity (see [18, 23] for a similar approach for a single locus). Finally, *b*_m_ and *b*_f_ control the shape of the sex-specific landscapes between *θ* and *θ*, such that landscapes are accelerating in males and females when *b*_m_ *>* 1 and *b*_f_ *>* 1 and diminishing when *b*_m_ *<* 1 and *b*_f_ *<* 1. For example, *b*_f_ *<* 1 and *b*_f_ *>* 1 would correspond to negative and positive second-order effects in a quadratic regression of female fitness on the trait *z* [62]. To aid presentation, we focus in the main text on the case where landscapes have the same shape in both sexes (*b*_m_ *= b*_f_ *= b*); this assumption is relaxed in Appendix C.3.

### 2.2 Genetic architecture, genotype-phenotype map, and evolution

We investigate the evolution of trait *z* under two models of its genetic basis: Kimura’s continuumof-alleles [51] model and a polygenic model. We assume initially that all loci are autosomal (nonrecombining and recombining segments of sex-chromosomes are considered later in section 3.3). These two models, which we detail below, provide complementary insights into the evolution of antagonistic traits by relaxing different assumptions about the genetic architecture of *z* that are typical of population genetic models of balancing selection [17–36]. Specifically, by allowing a wide range of possible alleles to arise through mutation, our first approach relaxes the assumption that only two alleles with fixed fitness effects may segregate at a sexually antagonistic locus. This enables us to characterise the nature of selection that arises from the male- and female-specific fitness landscapes. Meanwhile, our second model relaxes the assumption that traits are encoded by one or two loci by letting *z* be polygenic. Doing so allows us to investigate how polygenic variation responds to sex-specific selection.

#### 2.2.1 Continuum-of-alleles model

The continuum-of-alleles model [51] considers *z* to be encoded by a single additive locus and follows its evolution through recurrent mutations that each create a new allele whose value deviates from the original allele by a random amount (Appendix A for details). When mutations are rare, phenotypic evolution under this model can be characterised from an invasion analysis, the basis of which is the geometric growth rate *W* (*x_•_*, *x*) of a new allelic mutant with phenotypic effect *x_•_* arising in a population otherwise monomorphic for allelic effect *x* (e.g., chapter 12 in [52] for textbook). We assume throughout that allelic effects on phenotype are the same in males and females, so that homozygous resident males and females express *z =* 2*x* while heterozygous mutant males and females express *z_•_ = x_•_ + x*. In this case, invasion fitness equals

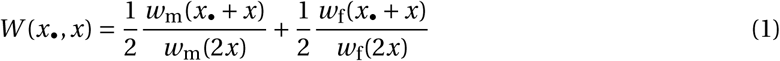

(Appendix A eqs. A-1-A-5 for derivation).

The assumption that allelic effects are the same across the sexes generates the strongest possible genetic trade-off between male and female fitness and so provides the most favourable conditions for sexually antagonistic polymorphism. Without this constraint, sexual dimorphism would readily evolve, resolving genetic conflict through the fixation of alleles with sex-specific effects [11, 12, 53].

When mutations have small phenotypic effects (i.e., *x_•_* and *x* are similar), evolution proceeds in two phases under the continuum-of-alleles model [52, 54–56]. Initially, the population evolves under directional selection, whereby positively selected mutations rapidly sweep to fixation so that the population transitions from being largely monomorphic for one trait value to being monomorphic for another. The population may thus converge to a singular trait value *z^*^ =* 2*x^*^*, which is such that *s*(*x^*^*) *=* 0 and *s^′^*(*x*^*^) < 0, where *s*(*x*) = *∂W* (*x*_•_, *x*)/∂x_•_|_x=x_ is the selection gradient acting on allelic value [57–60]. Once the population is fixed for *x*^*^, it experiences either: (i) stabilising selection when *h*(*x*^*^) = ∂^2^*W* (*x*_•_, *x*^*^)/∂*x*^2^_•_|*x*_•_ =*x*^*^ ≤ 0, such that the distribution of allelic effects remains unimodally distributed around *x^*^*; or (ii) disruptive selection, exhibiting negative frequency dependence, when *h*(*x^*^*) *>* 0 and therefore becomes polymorphic in a process referred to as ‘evolutionary branching’ [55, 59, 61]. In this latter case, selection maintains two alleles that initially encode weakly differentiated phenotypes around *x^*^*, which then become increasingly divergent as the two alleles accumulate further mutations [60]. We will refer to such negative frequency-dependent disruptive selection as ‘diversifying selection’ [61] for short.

#### 2.2.2 Polygenic diallelic model

In the polygenic model, we assume *z* is encoded by *L* loci such that the phenotype expressed by an individual of either sex is given by

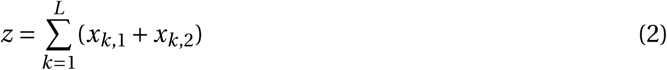

where *x_k_*_,1_ and *x_k_*_,2_ are the allelic values carried at the paternal and maternal copy at locus *k*, respectively (as in the continuum-of-alleles model, we assume here that allelic effects are fully sex-concordant). Genetic effects on phenotype are thus additive within and between loci, but they may translate into epistatic and dominance effects on fitness, depending on the fecundity functions (*w*_f_(*z*) and *w*_m_(*z*)). We assume that each locus *k* ∈ {1,…, *L*} has two possible alleles, A*_k_* and a*_k_* . Allele A*_k_* increases the size of the trait *z* and allele a*_k_* decreases it. Specifically, the effect of the allelic copy *x_k_*_,*l*_ inherited from parent *l* ∈ 2 {1, 2} at locus *k*is

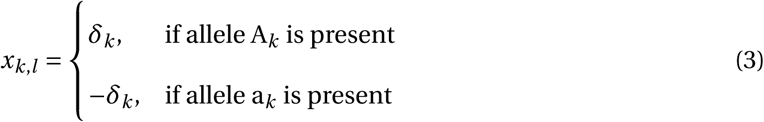

where *δ_k_ >* 0 is the absolute phenotypic contribution of a single allele at the locus. We initially assume that each locus contributes equally, with *δ_k_ = δ* for all *k*, so that the maximum and minimum phenotypes are 2*δL* and *−*2*δL*, respectively, which we assume correspond to the male and female optima, i.e., *µ =* 2*δL* (Appendix B for analysis).

With *L =* 1, this model reduces to classic single-locus population genetic models (e.g., [18, 23, 26], see Appendix B.1), and with many loci of small effects (*L* → ∞ and *δ* → 0) to quantitative genetic models of sexual antagonism (e.g., [12] in the absence of environmental effects on phenotype and with a cross-sex genetic correlation equal to one). Mutations from A*_k_* to a*_k_* and from a*_k_* to A*_k_* occur at a per-locus rate of *μ*. We assume that the *L* loci are evenly distributed across a single large chromosome, and in the main text that recombination is free between adjacent loci (i.e., occurs at a rate *r =* 0.5). We relax some of these genetic assumptions in Appendix C, where we allow for loci to have variable effect sizes (i.e. variation in *δ_k_* across *k* to model the presence of large and small effect loci, Appendix C.1) and for genetic linkage (i.e. *r <* 0.5, Appendix C.2). We find that none of these departures from our baseline model have a significant influence on the results presented below (as detailed in the Discussion).

## 3 Results

### 3.1 Strong sexual antagonism is required for diversifying selection

We first investigate evolutionary dynamics under the continuum-of-alleles model (section 2.2.1) with arbitrary sex-specific fitness landscapes (i.e., arbitrary *w*_m_(*z*) and *w*_f_(*z*)). We characterise the criteria for male- and female-landscapes to produce diversifying selection and so generate polymorphism in Appendices A.2-A.3. We find that two conditions must be satisfied. First, the population must have evolved such that on average individuals express an intermediate trait value *z^*^ =* 2*x^*^* for which

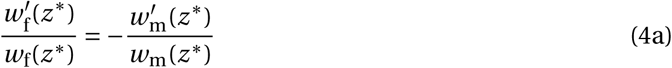

where *w ^0^* (*z^*^*)/*w_u_*(*z^*^*) is the slope of fitness landscape in sex *u* [62], i.e., such that a change in trait value has opposing and balanced additive fitness effects across the sexes. For example, under symmetric power functions (eq. I.A in Box I with *b = b*_m_ *= b*_f_), *z^*^* is halfway between the male and female optima when fitness declines with the same intensity away from each sex’s optimum (*c*_m_ *= c*_f_); otherwise, *z^*^* tends to be closer to the optimum of one sex (Fig. 2A). In the continuum-of-alleles model and in the absence of strong constraints on mutational input, convergence towards such a trait value *z^*^* is the expected outcome of evolution [57–60].

**Figure 1:**
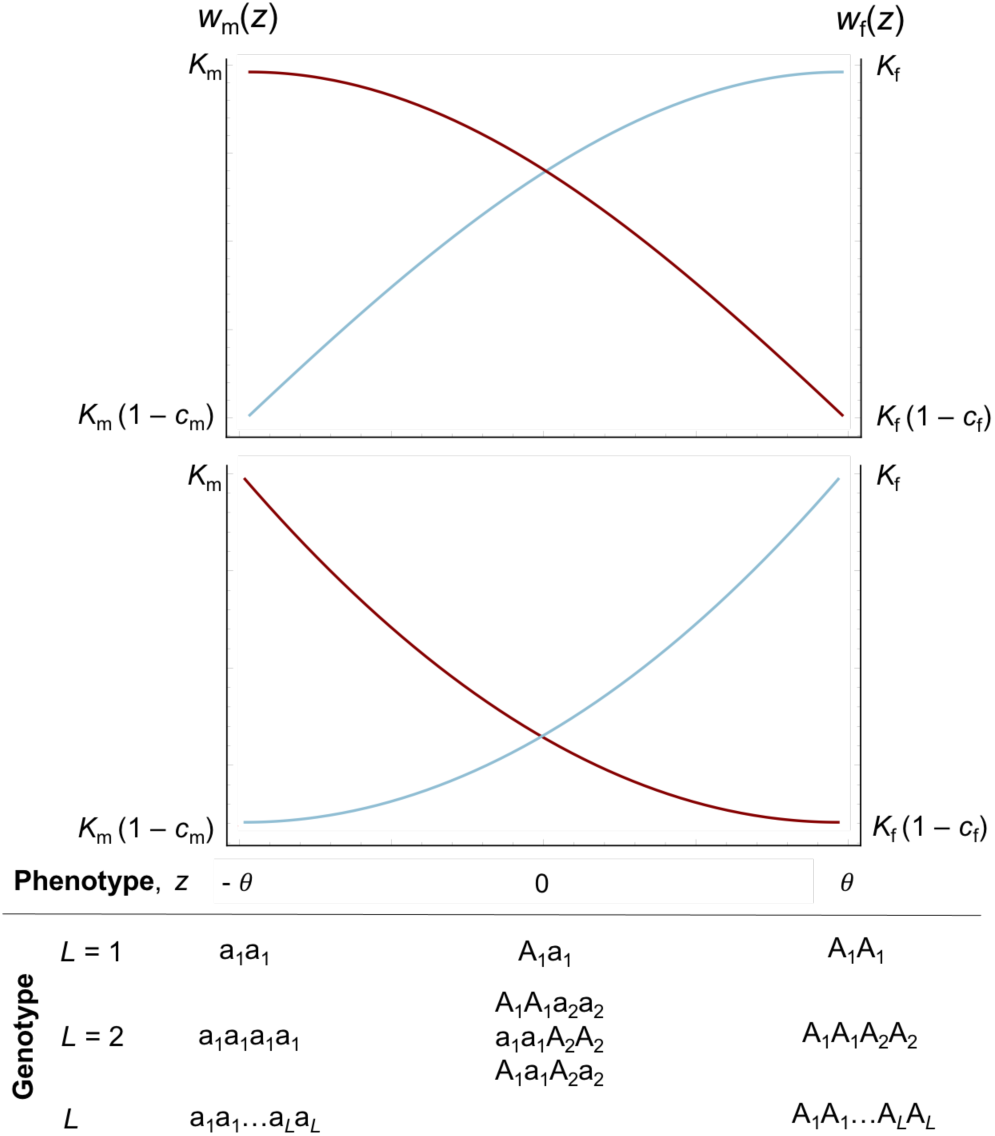
Sex-specific fecundity landscapes. Male (red curves) and female (blue curves) fecundity as a function of phenotype, *z*, eq. (I.A). A male’s fecundity is greatest when he expresses the phenotype *z = θ*, i.e., he is homozygous for allele “a” at every locus, while a female’s fecundity is maximised when she expresses *z = µ*, i.e she is homozgous for allele “A” at all loci (eqs. 2-3). Top and bottom panels show diminishing (*b*_m_ *= b*_f_ *<* 1) and accelerating (*b*_m_ *= b*_f_ *>* 1) returns to adaptation in each sex, respectively.

**Figure 2:**
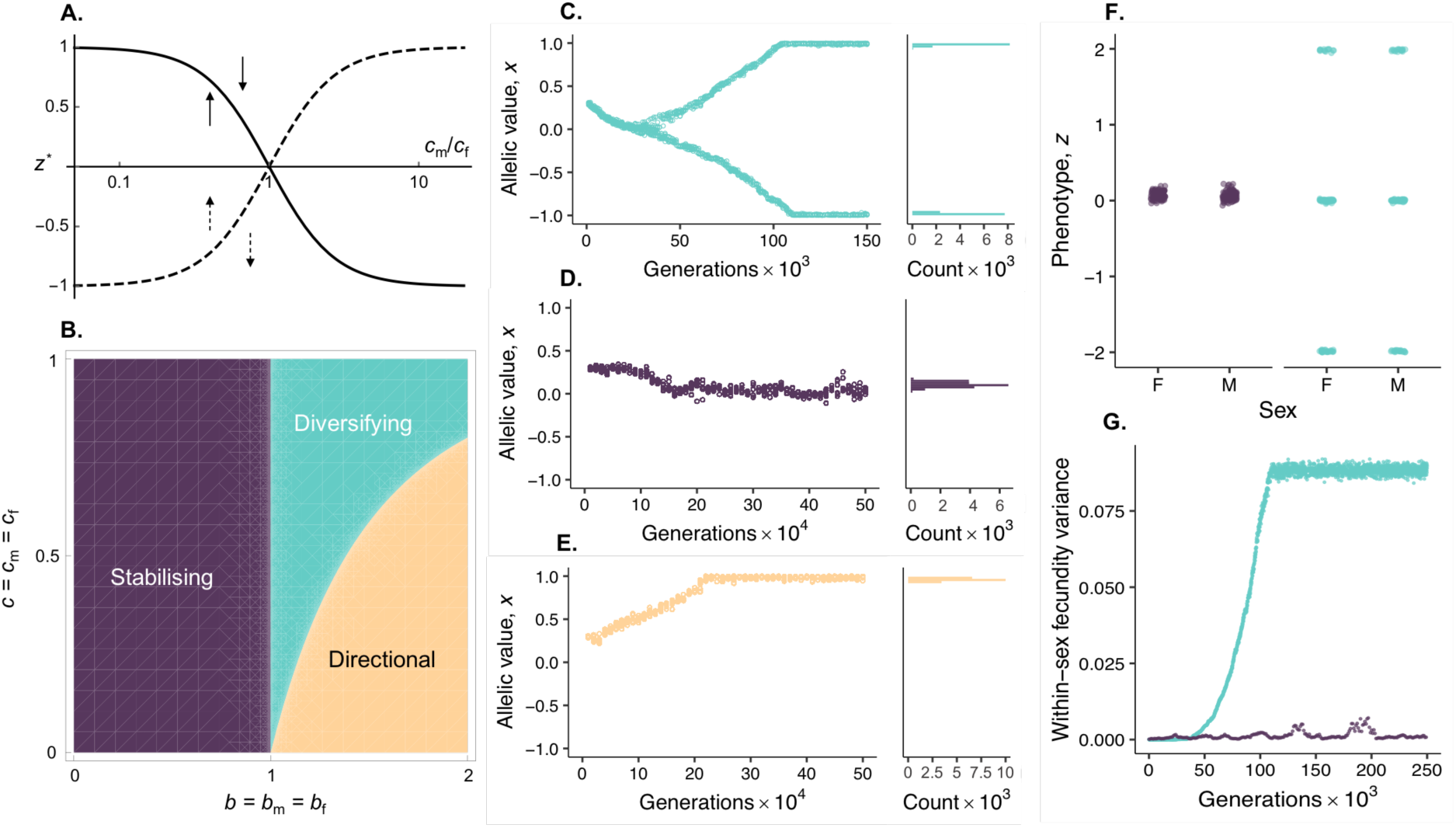
Genetic evolution in a continuum-of-alleles model of sexual antagonism. Panels show mathematical and simulation results for the continuum-of-alleles model when male and female fecundity follow power functions (eq. I.A, Box I). Panel **A** shows the singular trait value *z^*^ = µ ·* 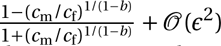 (where *ε* > 0 is a parameter of the order *c* and *c*_f_) as a function of the fecundity costs in males versus females *c*_m_/*c*_f_. Solid curve shows *z^*^* when it is an attractor of evolutionary dynamics (leading to “diversifying” or “stabilising” selection, see e.g. Fig. 2C and D), and dashed curve shows *z^*^* when it is a repellor so that selection favours evolution towards either the female or the male optimum depending on initial condition (“directional selection” see e.g. Fig. 2E). Panel **B** shows the parameter space leading to each mode of selection (Appendix A for full analysis). Panels **C**-**E** show evolution of allelic values *x* in a simulation of our continuum-of-alleles model when sexual antagonism leads to diversifying (*s =* 0.9, *b =* 1.5), stabilising (*c =* 0.05, *b =* 0.5), and directional (*c =* 0.05, *b =* 1.5) selection, respectively. In left-hand frame of each panel, dots represent a random sample of 10 alleles from the population at intervals of 10^3^ (in C) or 10^4^ (in D-E) generations. Righthand frames show the distribution of all alleles in the final generation. Simulations were initialised with all individuals homozygous for an allelic value of *x =* 0.3, with *µ =* 2, Appendix A.4.3 for full simulation procedure. Panel **F** shows the phenotypes of a random sample of 50 male and 50 female individuals from a simulation after 2.5 × 10^5^ generations under stabilising (purple dots) and diversifying selection (green dots). Phenotypic variance is much greater under diversifying selection where three types coexist: one expressing the male and another the female optimum, and a third intermediate. Panel **G** shows the average fecundity variance (Var[*w*_m_(*z*)]/2 *+* Var[*w*_f_(*z*)]/2) through time under diversifying (green dots) and stabilising (purple dots) selection. The variance in fitness increases when polymorphism arises under diversifying selection. Unless otherwise stated, in all panels the strength of sex-specific selection is equal across males and females, *c = c*_m_ *= c*_f_, and fecundity landscapes are the same shape, *b = b*_m_ *= b*_f_.

The second condition for diversifying selection is more restrictive: that *w*_m_(*z*) and *w*_f_(*z*) have special forms close to *z^*^*such that

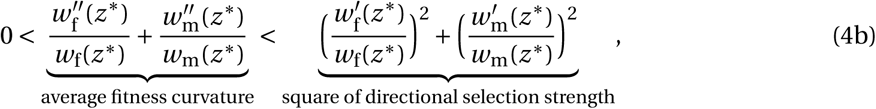

where *w ^00^*(*z^*^*)/*w_u_*(*z^*^*) is the curvature of fitness in sex *u* [62]. When *w ^00^*(*z^*^*)/*w_u_*(*z^*^*) *>* 0, fitness in sex *u* is accelerating (i.e., convex), whereas it is diminishing (i.e., concave) when *w ^00^*(*z^*^*)/*w_u_*(*z^*^*) *<* 0. The middle term of condition (4b) is therefore the curvature in the sex-averaged fitness landscape and indicates whether a population expressing *z^*^* is at a local minimum (curvature is positive) or a local maximum (curvature is negative) of this landscape. The right-hand term, meanwhile, is the strength of directional selection in each sex raised to the power of two. Condition (4b) thus tells us that, for selection to be diversifying, fitness must be on average accelerating around *z^*^* (middle term) but weakly relative to the strength of directional selection (right-hand term). This reflects the notion that diversifying selection here arises from the tension between two effects: on one hand, disruptive selection across both sexes against intermediate phenotypes (middle term); and on the other, strong competition within each sex that leads to negative frequency-dependent selection on maleor femaleadapted phenotypes (right-hand term). Generating the right balance between these forces is difficult, notably because it requires large variation in fitness for both sexes (i.e., large *w ^0^* (*z^*^*)/*w_u_*(*z^*^*)). In fact, condition (4b) reveals that whatever the fitness landscape, polymorphism will never arise under the continuum-of-alleles model when directional selection is weak, i.e., where terms of order [*w ^0^* (*z^*^*)/*w_u_*(*z^*^*)]^2^ or higher are negligible, as is often assumed, e.g., [26, 27, 35, 36, 53, 63]).

In line with these general observations, when sex-specific fitness follows power functions (eq. I.A, Box I), diversifying selection arises and generates a polymorphism only when fitness is accelerating (*b >* 1) and trades off strongly between males and females (*c*_m_ and *c*_f_ large and similar, green regions in Fig. 2B and Supplementary Fig. 1 top row panels, Fig. 2C for an example). Otherwise, selection across the sexes is either stabilising around *z^*^*, which occurs when fitness landscapes are diminishing (*b <* 1, Fig. 2B, purple region, Fig. 2D for an example), or directional towards the male or female optimum, which occurs when landscapes are accelerating (*b >* 1) but selection is not sufficiently strong to maintain polymorphism (Fig. 2B yellow region; here *z^*^* is a repeller of evolutionary dynamics and *z* evolves towards *θ* or *θ* depending on the value of *z* relative to *z^*^*, Fig. 2E for an example). Accordingly, phenotypic and fitness variance within sexes is strongly elevated when sexual antagonism drives diversifying selection, but is otherwise low (Fig. 2F-G).

### 3.2 Sexual antagonism diminishes polygenic variation unless selection is diversifying

Having characterised the types of selection that can arise from sex-specific fitness landscapes, we next investigate the response when *z* has a specified genetic architecture, in particular when it is underlain by multiple genes. To do this, we ran individual-based simulations of the polygenic model under a wide range of parameters that modulate genetic conflict (eq. 2 and eq. 3 with eq. (I.A) from Box 1 using SLiM 4 [64], see Appendix B.3 for details and also Appendix C.4 for an equivalent analysis with Gaussian fecundity functions). Results of these simulations show that the only conditions under which average heterozygosity across loci is greater than the neutral expectation are those that lead to diversifying selection (i.e., when condition (4b) holds, Fig. 3A). Here, heterozygosity is elevated because diversifying selection favours male- and female-adapted genotypes composed of different alleles at many loci, thus maintaining genetic polymorphism across sites. However, because genetic variation is shuffled by Mendelian segregation and recombination at each generation, diversifying selection leads to a wide distribution of phenotypes among individuals that is increasingly smooth and continuous as the number (*L*) of underlying trait loci increases (Fig. 3B). Predictably, the highest levels of heterozygosity and phenotypic variation are observed under conditions where diversifying selection is the most intense, i.e., where (i) fitness landscapes are strongly accelerating (*b ¿* 1), as this entails that intermediate genotypes show especially low fitness; combined with (ii) exceptionally strong sexual antagonism (*c*_m_ *ª* 1 and *c*_f_ *ª* 1) to oppose the fixation of the male or female-adapted genotypes (Fig. 3A). In addition, selection is most effective in driving elevated heterozygosity when the number *L* of loci that contribute to the trait is small relative to population size. This is because otherwise evolution at each individual locus becomes dominated by genetic drift (Supplementary Fig. 2).

**Figure 3:**
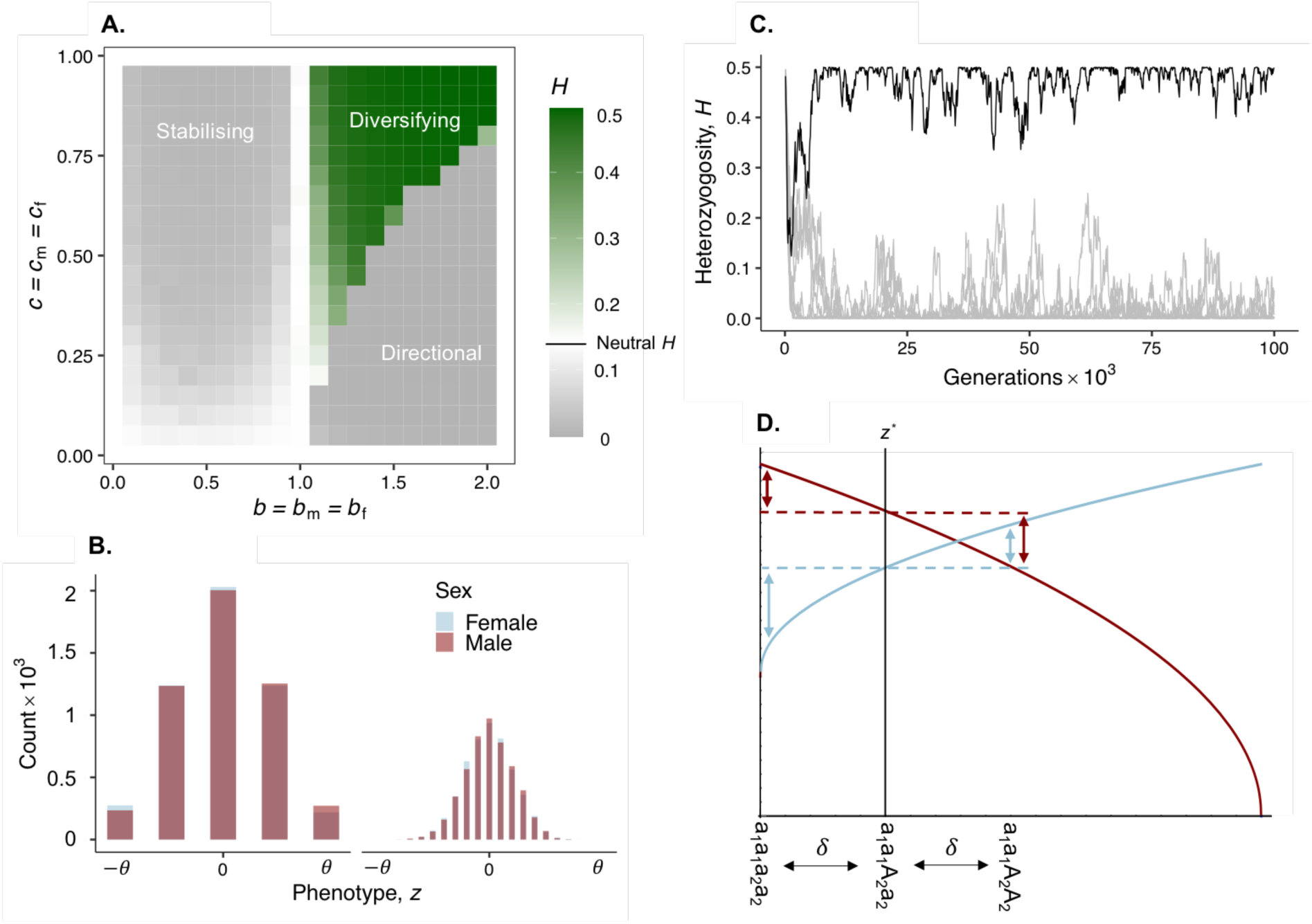
Genetic and phenotypic variation in a polygenic model of sexual antagonism. Panels show simulation results for our polygenic di-allelic model when male and female fecundity follow power functions (eq. I.A, Box I). Panel **A** shows average heterozygosity across loci from polygenic simulations. White, green and grey squares represent average heterozygosity values that are respectively equal, higher, and lower, than that of a neutral locus at mutation-drift balance (Appendix B.3 for full simulation procedures – unless otherwise stated, all panels show results from simulations with *L =* 10 di-allelic loci, *δ =* 1 and *µ =* 10^−5^). Panel **B** shows the distribution of male and female phenotypes of a polygenic trait (*L =* 2 on left and *L =* 10 on right) under diversifying selection (*c*_m_ *= c*_f_ *=* 0.9, *b =* 1.5) after 10^5^ generations of a simulation (note that male and female distributions are mostly indistinguishable). Panel **C** shows heterozygosity through time at each individual locus of a polygenic trait (*L =* 10) in a simulation where fitness declines more strongly in males than females (*c*_m_ *=* 0.85 × 0.3, *c*_f_*=* 0.15 × 0.3, *b =* 0.5), black curve shows heterozygosity at the “remainder” locus. Panel **D** is a graphic showing male (blue curve) and female (red curve) fecundity under stabilising selection (*b <* 1) when *c*_m_ *> c*_f_ (*c*_m_ *=* 0.85 × 0.3, *c*_f_ *=* 0.15 × 0.3, *b =* 0.5, as in panel C). Dotted horizontal lines show sex-specific fecundity for individuals expressing the optimal phenotype *z^*^* (which here is closer to the male optimum, i.e., *z^*^ <* 0). Vertical arrows show the fecundity of double homozygous genotypes in a two-locus model (*L =* 2) compared to intermediate genotypes that are homozygous for the male beneficial allele at locus 2 (a_2_) and heterozygous at locus 1. These indicate dominance reversal at locus 2 (with downward arrows shorter than upward arrows in each sex).

Diversifying selection that drives high heterozygosity levels also generates a strong sex load (Supplementary Fig. 3). For example a two-fold increase in heterozygosity relative to neutrality – equivalent to that produced by a doubling of the mutation rate – is associated with a minimum average fitness reduction of *ª* 20% across the sexes (a significantly stronger load is produced when fecundity follows Gaussian functions, Supplementary Fig. 3).

By contrast, if sexual antagonism does not lead to diversifying selection, average heterozygosity is diminished relative to neutrality. Under directional selection, selection continuously pushes for the optimum of one sex, favouring the fixation of either all A or all a alleles across loci. Meanwhile stabilising selection reduces heterozygosity because it favours genotypes that encode the equilibrium phenotype *z^*^* in homozygous rather than heterozygous state. This is because homozygotes are more likely to produce offspring also expressing *z^*^*, i.e., their genotypes are less likely to be broken down by random Mendelian segregation (an effect well-known to occur under sex-concordant stabilising selection [46, 65, 66]). Genetic variation observed in these cases is limited to the ephemeral segregation of alleles owing to recurrent mutation and genetic drift (Supplementary Fig. 4A, e.g., when *L* is large or selection weak).

Although stabilising selection reduces the average heterozygosity across loci relative to neutrality, in some instances we observe the stable maintenance of a balanced genetic polymorphism at a maximum of one locus (out of the *L* that code for the trait, Fig. 3C). This occurs whenever no homozygous genotype codes for a trait value close enough to *z^*^* ([66] for a description of this under sex-concordant selection). Polymorphism in this case is maintained at one “remainder” locus that shows sex-specific dominance reversal: both alleles have a larger effect in the sex they benefit than in the sex they harm (owing to the diminishing shape of the fitness landscapes, see Fig. 3D for a graphical representation and Appendix B.2.2 for analysis of *L =* 2 case). Such a remainder locus reflects a genetic constraint preventing the population from evolving to *z^*^*. This is the underlying assumption behind population genetics models that see the maintenance of polymorphism at a single sexual antagonistic locus under weak selection [18, 23, 37]. In our polygenic model however, genetic drift and recurrent mutation drive turnover in the identity of the remainder locus, so that unless selection is particularly strong, any one site shows only transient polymorphism here (Supplementary Fig. 4B, also Appendix C.1 for this effect when loci vary in their effect size).

Some of the above findings show apparent similarities with those of [46], who analysed a multilocus population genetics model where selection favours the same optimum in males and females but where alleles can have sex-dependent effects on trait expression, leading them to be potentially sexually antagonistic. It is shown that such sexual antagonism maintains a maximum of one balanced polymorphic locus under weak Gaussian selection and loose linkage (which allows for a “Quasi Linkage Equilibrium approach” [46]). This is the same outcome as seen in our model under weak selection, and the reason for this convergence is that in both cases selection across the sexes is stabilising when variation in fitness is small (see App C.6, eq. C-26 for the conditions for stabilising or diversifying selection in [46]’s model). However, [46] find no cases of polygenic polymorphism when selection is strong (unlike our results under diversifying selection). Specifically, [46] numerically iterated recursions for 10 additive loci with randomly and independently sampled sex-dependent allelic effects and found that no more than two loci ever remained polymorphic in the long run. Presumably, polygenic variation is not maintained in [46] because diversifying selection in their model necessitates large systematic differences in male and female phenotype (see App. C.6, eq. C-26), and parameter combinations leading to such systematic differences are extremely unlikely under the reasonable assumption that allelic effects are independently distributed across loci (as assumed in [46]). More broadly, the discrepancy between [46]’s results and ours demonstrates the distinct biology of sexual antagonism that arises from sex-specific landscapes vs. sex-specific allelic effects, leading to differences in the scope for balancing selection across loci.

All our analyses thus far assume that the phenotypic distribution in the population is initially unimodal, i.e., that individuals rarely differ strongly in their value for *z*. We relax this in Appendix C.5 to consider the consequences of sexually antagonistic selection for two already-differentiated genetic morphs, such as diverged alleles that have arisen through large effect mutation or gene flow. We show in this Appendix that there exist sex-specific fitness landscapes *w*_m_(*z*) and *w*_f_(*z*) that are such that they do not lead to diversifying selection (i.e., condition 4b does not hold), but do allow for a pre-established polymorphism at one locus to be robust to small effects mutations. In other words, there are situations where sexual antagonism will stably maintain a coalition of two alleles with highly divergent phenotypic effects although these alleles cannot emerge from gradual evolution under the continuum-of-alleles model. Such situations however have limited scope to contribute to genetic variation, and this is for three reasons. First, they require sex-specific fitness landscapes that show very particular properties (Appendix C.5.1, eq. C-20). Second, these sex-specific fitness landscapes favour polymorphisms that are sensitive to fluctuations in allele frequency (e.g., due to genetic drift) or further large effect mutations (Appendix C.5.1, eq. C-23). Finally and most importantly, these polymorphisms break down under recombination, such that sexually antagonistic selection here cannot maintain elevated polygenic variation when a trait is encoded by multiple smaller effect loci (Appendix C.5.2). This reinforces the notion that balanced sexually antagonistic polymorphism requires either diversifying selection or genetic constraints on allelic values, with polygenic variation being extremely difficult to maintain in the latter case.

### 3.3 Sex chromosomes and PAR: cold and hotspots for diversifying selection

Our results suggest that the scope for sex-specific selection to maintain polymorphism at autosomal loci is narrow. We therefore extend our analyses to consider sex-linked loci, as they may be more amenable to sexually antagonistic polymorphism. In particular, models of single di-allelic sex-linked loci with arbitrary dominance and selection coefficients have suggested that conditions for balancing selection may be especially lenient on sex chromosomes [19, 21, 67], usually based on arguments of allelic dominance (e.g., that the absence of male dominance for X-linked promotes the invasion of male-beneficial alleles [19], although see [23, 34]). This has led to much discussion over whether sex-chromosomes or autosomes should be expected to harbour higher levels of sexually antagonistic variation [19, 21, 27, 34, 36, 67–69].

We first investigate the non-recombining portion of the X-chromosome (or Z-chromosome in ZW species) where such loci do not have Y-linked/W-linked homologues contributing to the same trait (hemizygous regions). We derive the relevant conditions for diversifying selection in Appendix D.1. The equations describing these conditions are slightly involved (eqs. D-5 and D-20 in Appendix D) and so we refrain from giving them here, but they reveal that the criteria for diversifying selection are typically less permissive at X-linked than autosomal loci (i.e., compared to eqs. 4a-4b). Thus, in contrast to some classic single-locus results (e.g., [19, 21]), we find the X-chromosome is in fact a coldspot for polymorphism underlying sexually antagonistic traits. This is a direct consequence of the sexbiased inheritance of X-chromosomes (eq. D-20 in Appendix D for formal argument): while females pass on half their X-linked genes to their sons and daughters, males pass on X-linked genes only to daughters. This leads to a bias whereby male-beneficial alleles are disproportionately inherited by females, as these genes are over-represented in high fecundity fathers. Simultaneously, because males only receive X-linked genes from their mother, an equivalent bias arises whereby female-beneficial alleles tend to be more frequently expressed in males than females, because these alleles are overrepresented in high fecundity mothers. Sexually antagonistic alleles therefore become associated with the context in which they confer low fitness (i.e., male-beneficial alleles with female-carriage and female-beneficial alleles with male-carriage), making it more difficult to maintain polymorphism in the population [56].

We also consider loci that sit in the “pseudo-autosomal region” (PAR) [67–69] of the sex chromosome, linked to a sex-determining region (SDR) with recombination rate *r ∏* 0 (Appendix D.2 for derivation and eqs. D-22-D-23 for conditions for diversifying selection). In short, we find that diversifying selection owing to sexual antagonism occurs more readily on the PAR than on autosomes or on nonhomologous (hemizygous) sex chromosome regions, especially where recombination with the SDR is rare, supporting the notion that these regions are conducive to balancing selection [67, 69]. Sexually antagonistic polymorphism in the PAR may even arise under weak selection when linkage with the SDR is tight compared to selection (specifically where *r* is of the order of the strength of directional selection raised to the power of two, eqs. D-22 and D-23 in Appendix D for details). This greater scope for polymorphism on the PAR is because linkage with a SDR allows male-beneficial alleles to become associated with Y-carriage (and so with male expression), and female-beneficial alleles to become associated with X-carriage (and so, on average, with female expression). This generates a positive correlation between alleles and the genomic context in which they confer high fitness (in contrast to the non-recombining X-chromosome), relaxing the conditions for both alleles to co-exist adaptively in the population (eq. D-22 in Appendix D [56]). Nevertheless, while the PAR may be a hotspot for diversifying selection, it is difficult to see how this would translate into the maintenance of polygenic variation, due to the requirement that large numbers of loci affecting the same trait are all in tight linkage with the SDR.

### 3.4 Sexual antagonism produces weak genomic signatures

Finally, as a last step we ask whether sexually antagonism at quantitative trait loci that happen to segregate, whether due to mutation-selection-drift balance or to the rare cases of balancing selection, produces signatures that may be found in sequence data. To answer this, we analysed the simulated data from our polygenic model with two population genomic approaches that are commonly leveraged to identify sexually antagonistic loci, focusing on autosomal loci for simplicity (Appendix E for full procedures and calculations). First, we tested whether sexually antagonistic selection drives allelic differentiation between the sexes. This is typically quantified as between-sex *F*_ST_ [13, 70–74], which in the context of sexually antagonistic fecundity selection is measured amongst the successful parents in a given generation [74]. Second, we tested whether sexually antagonistic selection leads to detectable associations between allelic state and sex-specific fitness, as in GWAS for sex-specific fitness and sexual antagonism [15, 73, 75] (our model represents a best-case scenario for this test, as we assume *z* is under pure genetic control with full heritability). To quantify sex-differences in fitness associations, we computed *|β*_m,*k*_ *−β*_f,*k*_ | for every locus *k*where *β*_m,*k*_ and *β*_f,*k*_ are the linear regression slopes of reproductive success (i.e., number of recruited offspring) in males and females, respectively, against within-individual frequency of allele A at locus *k*. See Appendix E for details on these analyses.

Fig. 4A shows that between-sex allelic differentiation is generally weak for quantitative trait loci under sexually antagonistic selection and that the distribution of *F*_ST_ across loci is in fact largely similar to that under neutrality. Moreover, this differentiation is also similar to that where either male or female fecundity (*w*_m_ or *w*_f_) is randomly assigned for each individual regardless of their sex. This is expected, as generating appreciable between-sex *F*_ST_ values typically requires strong sex-specific selection acting on a given locus [72, 76], and sexually antagonistic selection acting on a polygenic trait translates into weak effects on each individual locus. One exception is where sexual antagonism causes strong diversifying selection (Fig. 4A, green boxplots). This leads to elevated allelic differentiation among the sexes, although this effect declines significantly with the number of loci encoding a trait (e.g., with *c*_m_ *= c*_f_ *=* 0.6 an *F*_ST_ of the order of 10^−2^ is generated at a single locus, see Fig. 5 in [72], which also captures our model with *L =* 1, but only an *F*_ST_ of the order 10^−4^ when *L =* 10, Fig. 4A). This indicates that measures of genetic differentiation such as sex-specific *F*_ST_ will produce a signal for sexually antagonistic quantitative trait loci only under the relatively extreme biological scenarios that lead to diversifying selection. However, because polygenicity spreads selection thinly across loci, this signal is nonetheless weak and detecting it will require the availability of particularly large datasets or methods that combine *F*_ST_ patterns across the genome (e.g., by relating to the degree of biased gene expression [71]).

**Figure 4:**
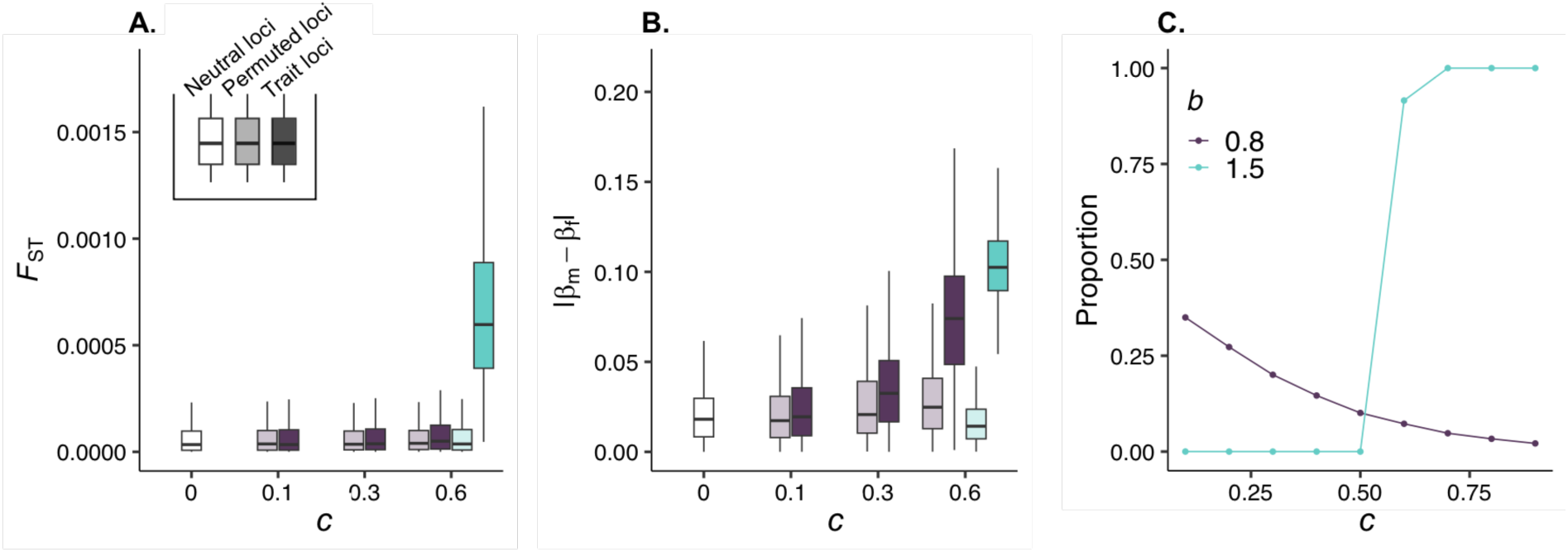
Patterns of allelic variation and fitness associations in polygenic simulations. Panels show results from pooled data for simulations of our di-allelic polygenic model when male and female fecundity follow power functions (eq. I.A, Box I, see Appendix E for full simulation procedures and calculations, parameters used: *L =* 10, *δ =* 1, *µ =* 10^−5^). Panel **A** shows distributions of allelic differentiation (between-sex *F*_ST_). Panel **B** shows distributions of absolute sex-differences in the linear regression coefficients of fitness against within-individual allele frequency (*|β*_m,*k*_ *−β*_f,*k*_ |). In both panels, white boxplots show values for neutral alleles, coloured boxplots show values for sexually antagonistic trait loci (red for *b =* 0.8, green for *b =* 1.5, only parameter values leading to either stabilising or diversifying selection were used, with diversifying selection only arising when fecundity costs were strong enough, *c =* 0.6, to satisfy eq. 4b, see eq. A-19), and translucent coloured boxplots show values for permuted loci as controls (i.e., trait loci where the sex of each individual is randomly assigned). Boxes show the 25^th^, 50^th^ and 75^th^ quantiles, vertical black lines show the interquartile range. Panel **C** shows the proportion of alleles with minor-allele-frequency greater than 0.05 for loci encoding a trait under sexually antagonistic selection.

When feasible, performing genetic associations with fitness may offer a more productive approach. Indeed, we observe that compared to controls, sexual antagonism drives an increase in *|β*_m,*k*_ *−β*_f,*k*_ | when selection is either stabilising or diversifying, so long as selection is sufficiently strong (i.e. *c*_m_ and *c*_f_ not too small, Fig. 4B). Sex-specific fitness associations are possible when selection is stabilising for *z^*^* because fitness landscapes in each sex can be steep close to *z^*^* (e.g., see Fig. 1 Top where *z^*^ =* 0). As a consequence, genetic variation segregating either at a remainder locus or ephemerally owing to mutation and drift can covary noticeably with male and female fitness in opposite ways. However, detecting sexually antagonistic loci when stabilising selection is strong may prove challenging. This is because, outside of a remainder locus, selection is effective in eliminating variation in this case, such that most loci either fix or show low minor allele frequency (Fig. 4C), leaving them unamenable to statistical calculations unless large volumes of data are available. Together, our results thus support the notion that genomic approaches for identifying sexually antagonistic alleles show strong limitations [72–74, 77]. In particular, they highlight that, owing to a lack of balancing selection, detecting antagonistic loci underlying polygenic traits is likely to be even more difficult than suggested by prior single-locus theory (73 for review).

## 4 Discussion

### 4.1 A narrow scope for sexual antagonism to maintain genetic variation

Our analyses indicate that sex-specific selection on shared continuous traits can drive above-neutral levels of genetic variation, but that this occurs only under restrictive conditions that should not commonly be met in natural populations. This is because diversifying selection is needed to maintain elevated heterozygosity when traits depend on many alleles and such selection typically entails strong conflict over trait expression between males and females. That is, selection in each sex must act in opposing directions, be both strong and of similar intensity in both sexes, and must occur on an accelerating fitness landscape: a restrictive combination of conditions. For example, using datasets of collated sex-specific selection gradients from wild populations (634 trait estimates [78]) and humans (34 trait estimates [79]), we found no traits for which estimated selection gradients were consistent with diversifying selection (i.e., we found no traits for which selection gradients satisfied eq. 4b, Appendix F for details).

Moreover, where sexual antagonism is strong enough to generate diversifying selection, it should rarely last over substantial evolutionary periods. This is because random Mendelian segregation leads phenotypic variation in a polymorphic population to remain distributed around an intermediate value (Fig. 3B) that confers especially low fitness in both sexes. Consequently, even a modest boost to heterozygosity in a polygenic sexually antagonistic trait is associated with a strong depression in mean fitness (Supplementary Fig. 3). Owing to this large load, selection to resolve conflict [1, 12] (through, e.g., the evolution of sex-specific gene expression [80]) is very strong. Therefore, such polymorphism is unlikely to persist over extended evolutionary periods unless there are extreme constraints on genetic architecture that preclude sexual dimorphism [11, 12].

If conflict over trait expression is not strong enough to elevate heterozygosity in a continuous trait, sexual antagonism acts to erode genetic variation. When this happens, balanced sexually antagonistic polymorphism can be maintained only in two specific cases. First, when sexual antagonism causes stabilising selection for an optimal phenotype that is unattainable by any homozygote, polymorphism can develop at a maximum of one remainder locus owing to patterns of allelic dominance (sex-specific dominance reversal for an autosomal locus, Fig. 3C-D, and recessivity in females for an X-linked locus, Appendix D; see [10, 18, 19, 23] for single locus models). However, variation is unlikely to be long-lasting at any given site as mutation and drift drive regular turnover in the identity of the remainder locus (Supplementary Fig. 4, Appendix C.1), and because polymorphism is not robust to the appearance of new alleles that encode the optimal phenotype when homozygous. Second, polymorphism can occur at a locus when two highly diverged alleles are present before the onset of sexual antagonism, or arise through large effect mutations or through gene flow (Appendix C.5 for details). However these allele coalitions are especially sensitive to fluctuations in allele frequency (e.g., due to genetic drift) or to further large effect mutations, so that a polymorphism of this sort is also unlikely to show long term persistence.

In fact, the only circumstance where we find permissive conditions for elevated sexually antagonistic polymorphism when multiple alleles affect a trait is when loci are in tight linkage with the sex-determining region (for di-allelic models of this, see [67, 69]). Diversifying selection can occur here because genetic linkage allows sexual antagonism to drive an association between the sex in which an allele is beneficial and the sex in which an allele resides more commonly, so that polymorphism becomes easier to maintain in the population [56]. In this way, our results echo conditions for the maintenance of polymorphism owing to local adaptation to two different environments connected by dispersal (e.g. [81–85]). In these models, polymorphism occurs only under limited dispersal (analogous to the recombination rate being low here) as this allows alleles to become over-represented in the environment in which they are adaptive; otherwise gene flow breaks this association and thus inhibits polymorphism. (In fact, a two patch local adaption model with soft selection [86] and random dispersal is equivalent to a model of autosomal sexual antagonism [87], and with limited dispersal to a model of sex-specific selection on the PAR, see App. C.6 and G for details and further discussion on this connection.) Nevertheless, we do not expect diversifying selection in the PAR to drive polygenic polymorphism, due to the requirement that multiple loci influencing the same trait be present in this region and in tight linkage with an SDR.

Altogether, our results indicate that not only are the conditions for polygenic variation restrictive (requiring strong sexual antagonism), but also that the long-term maintenance of sexually antagonistic polymorphism at any locus requires strong genetic constraints: (1) Constraints on gene expression that maintain strong intersexual genetic correlations i.e., such that diversifying selection will not quickly drive the evolution of sexual dimorphism; (2) Constraints on the possible allelic variants that may arise at an antagonistic locus (in order for polymorphism at a remainder locus to arise under stabilising selection); (3) Constraints on the rate of recombination between an antagonistic locus and the sex-determining region (allowing for tight linkage between these loci and therefore for diversifying selection without imposing a strong sex load). This conclusion is in spite of a number of assumptions in our model that maximise opportunity for sexual antagonism to produce balancing selection; namely a perfect intersexual genetic correlation and the absence of environmental effects on trait expression. Without either of these assumptions, the scope for strong and enduring sexual antagonism is significantly reduced [11, 12], indicating our results are in fact conservative. Moreover, our conclusions were also robust to relaxing two constraints from our polygenic model (Appendix C): that alleles at all loci have equal phenotypic effect sizes and that recombination between loci is free. We found that neither of these assumptions qualitatively affected the parameter conditions for polymorphism (we discuss implications of the ecological and genetic assumptions of our model in depth in Appendix G).

Our conclusions may appear at odds with previous results stemming from other models based on fitness landscapes [10, 88]. In particular, Conallon and Clark [10] consider a continuous trait evolving on a two-sex fitness landscape. This trait may be polygenic but it is assumed that mutations are rare enough that only one site segregates at a time (so that the trait can only ever show polymorphism at a single locus). They calculate the probability that a mutation that arises in a monomorphic background and that may have sex-specific effects on the trait experiences balancing selection. They find that this probability can be high owing to sex-specific dominance reversal for fitness that emerges from concave/saturating landscapes close to the optima (see also [37]). These insights have led multiple authors to suggest that sexual antagonism can drive genetic variation in polygenic traits due to balancing selection (e.g., [10, 13, 30, 37, 38, 41]). However, our analyses show that balancing selection here is a direct consequence of the genetic constraints imposed by the assumption that only a single locus ever segregates. Sexual antagonism acting on a polygenic trait in fact typically opposes genetic variation across loci, leading to a balanced polymorphism at a maximum of one locus irrespective of the number of loci encoding a trait. In other words, while concave/saturating fitness landscapes may be conducive to single-locus polymorphisms, they do not drive elevated genetic variation in a polygenic trait. Rather, selection must be diversifying, which entails very different landscapes to those highlighted in previous models: requiring accelerating/convex and steep fitness curves. However, because of the strength of selection and the sex load they induce, we expect these landscapes to be rare.

### 4.2 Empirical implications

Our analyses provide several insights for empirical work on sexually antagonistic polymorphism. First, our results indicate that traits under sexually antagonistic selection will typically exhibit relatively little additive genetic variation. Where sexually antagonistic traits are found to show high levels of standing variation at autosomal loci, we expect such traits to be under strong selection with fitness landscapes being on average accelerating across the sexes, two necessary conditions for generating diversifying selection. Empirically, this could be tested directly by fitting quadratic regressions of reproductive success on trait values in each sex [62]. We note, however, that non-linear selection gradients on sexually antagonistic traits are rarely significant (e.g., [42, 89, 90]), likely due to the requirement of large datasets to detect second-order fitness effects. In principle, the effect of sexual antagonism on levels of genetic variation could also be assessed experimentally, by measuring the impact of manipulating the strength and nature of antagonistic selection on levels of additive genetic variation for fitness. Studies using family-based selection for sexual dimorphism (e.g., [41, 91]) provide templates for such an approach. One such study [41] found that standing genetic variation did decrease under antagonistic selection, albeit less so than under other modes of selection.

At the genetic level, our results suggest that antagonistic selection on continuous traits leaves few signatures in patterns of polymorphism within populations—except in the presence of very strong diversifying selection (Fig. 4). Therefore, in most cases, genetic signatures of sexual antagonism will be subtle and correspond mostly to allelic variation that is maintained due to mutation. Whether and to what extent these signals are detectable will depend on circumstances. Most importantly, detectability will depend on a number of experimental parameters that have predictable effects on statistical power and that we have not explored here, such as sample size and the degree of environmental noise in the trait. Beyond these, our results indicate that the approach used to detect genetic signatures will have a major impact on power. In particular, we find that both methods based on between-sex genetic differentiation (e.g., sex-specific *F*_ST_ [13, 70–74]) and fitness associations (e.g., GWAS for sex-specific fitness [15, 73, 75, 79, 92]) are constrained by the fact that sexual antagonism typically reduces genetic variation and so leaves few loci at which any detectable signal of sexual conflict would be present. Indeed, these signals may easily be drowned in genomic background noise unless a candidate set of loci is known. Even where a locus is detected, it would necessarily provide a poor picture of the loci that are subject to antagonistic selection, which will include many lacking variation. Furthermore, when sexually antagonistic variation does segregate (either due to mutation or balancing selection), sexspecific *F*_ST_ methods likely have especially low power to detect individual trait loci that contribute to phenotypes with a wide genetic basis. This is because such a signal is based on frequency change over a single generation [72] and selection on individual loci underlying a polygenic trait is weak unless sexual antagonism leads to extremely strong diversifying selection at the phenotypic-level (although this may be less problematic for approaches looking at genome-wide signs of sex-specific selection and so gain power across loci [71]). Meanwhile, approaches based on fitness associations (e.g., GWAS for sex-specific fitness [15, 73, 75, 79, 92]) are more powerful—relatively speaking, but nonetheless, the small per-locus effects of sexual antagonism also compromises their usefulness where selection is not strong.

A corollary to our finding that sexual antagonistic selection acting on a given trait typically maintains only a single polymorphic locus is that: where multiple sexually antagonistic loci showing elevated genetic variation are identified, these most likely underlie independent traits. This is consistent with antagonistic loci detectable by GWAS approaches showing no functional enrichment [15]. Additionally, the predicted turnover in the identity of trait loci harbouring polymorphism at any given time under stabilising selection fits with the observation that polygenic variation underlying traits under antagonistic selection shows little signal of long-term balancing selection [74, 93]. Our results therefore suggest that the rare loci where a strong signal of antagonism and balancing selection is detectable are expected to be those showing large phenotypic (and hence fitness) effects. It is these loci that are most likely to generate the genetic constraints that are necessary to maintain long-term polymorphism under stabilising selection across the sexes. Indeed this is true for allelic variants at the *VGLL3* gene in Atlantic salmon that generate a sexually antagonistic polymorphism in maturation time and account for 39% of trait variation [94]. Similarly, large and experimentally replicable effects were demonstrated at the *DsFAR2-B* locus that underlies a sexually antagonistic polymorphism in cuticular hydrocarbon profiles in the tropical fly *Drosophila serrata* and that shows signatures of balancing selection [95].

### 4.3 Conclusions

Previous theory on sexual antagonism has relied overwhelmingly on models of Mendelian traits that are determined by a limited number of alleles with fixed effects [17–36]. This reliance may have led to the expectation that sexually antagonistic genetic polymorphisms are a regular feature of sexual species [10, 19, 23, 30, 37, 38, 40, 41] and that such polymorphisms should be detectable from genomic data ([13, 73, 96]). Our work has shown that this expectation rarely extends to continuous traits. By considering some of the complexity behind such traits, our model lays bare that previous conditions for polymorphism are direct consequences of constraining the number and the nature of alleles that can affect a trait. These conditions do not generalise to cases where a wider variety of alleles can arise either within or across loci. In fact, our analyses indicate that sexual antagonism is not a straightforward driver of the long-term maintenance of polymorphism at loci underlying continuous traits and rarely produces sex-specific patterns of variation outside of sex-determining regions. We therefore suggest that most sexually antagonistic polymorphisms associated with quantitative trait loci are transient and should be observed no more frequently than at neutral sites, except when males and females face strongly divergent fitness requirements or impregnable genetic constraints are operating.

## Acknowledgements

We thank Seth Watt for extensive discussions about evolution with sex-differences in selection and for exploring and sharing early simulations with different fitness functions not included here. We also thank Tim Connallon, Filip Ruzicka and Carl Mackintosh for useful discussion.

## Funding

We are grateful to the Swiss National Science Foundation (PCEFP3181243 to CM), the Biotechnology and Biological Sciences Research Council (BB/W007703/1 and BB/T019921/1 to MR), the Leverhulme Trust (RPG-2021-414 to MR), the Natural Sciences and Engineering Research Council of Canada (NSERC RGPIN2022-03726 to SPO), and the UK Natural Environment Research Council (NERC grant to VS and QMEE CDT Studentship to EF) for funding.

## Supplementary Figures

**Supplementary Figure 1:**
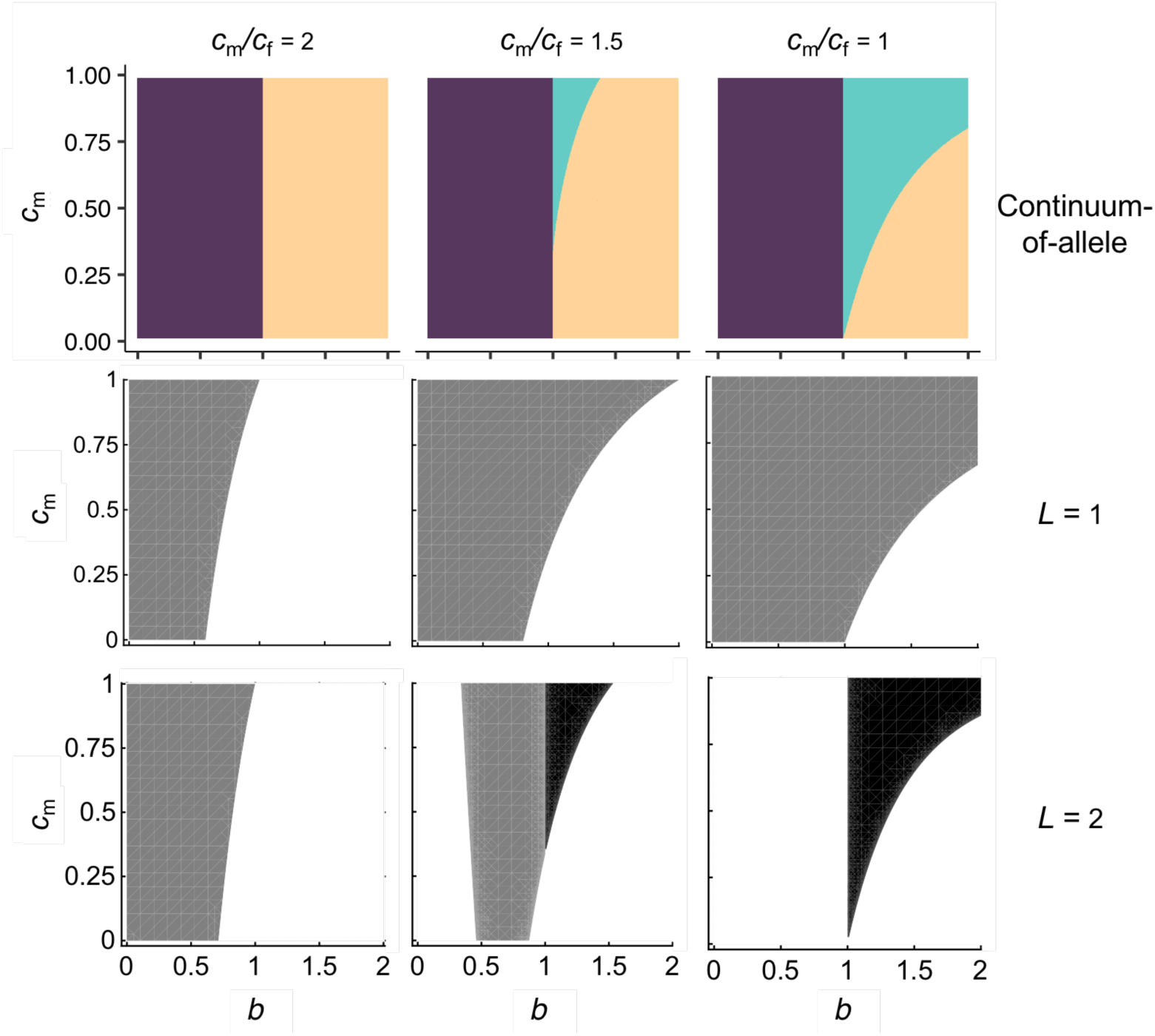
The mode of selection and maintenance of genetic variation in the continuum-of-alleles and di-allelic models. Top row panels show parameter space corresponding to stabilising (purple regions), directional (yellow regions), and diversifying (green regions) selection in the continuum-of-alleles model when male and female fecundity follow power functions (eq. I.A, Box I). These plots were generated by numerically solving eq. (A-7) to find singular points and evaluating their stability (Appendix A.4.2 for details). Middle row panels show parameter conditions for polymorphism (grey regions) at a single locus di-allelic (*L =* 1, *δ =* 1, see eqs. 3-2). Polymorphism condition corresponds to when inequality (B-1) is satisfied. Bottom row panels show polymorphism space for a trait encoded by two di-allelic loci (*L =* 2, *δ =* 1). Grey regions show conditions for polymorphism at one locus (i.e., the other is fixed for one allele), and black region shows conditions for polymorphism at two loci. Grey regions correspond to the conditions where all four haploid genotypes are invadable by another genotype but there is a stable equilibrium where one locus is fixed for one allele, while black regions correspond to parameter space that we found such an equilibrium did not exist (found by numerically iterating eq. (B-4)-(B-5), Appendix B.2.2 for details). In all panels, parameter space is in terms of the shape of fitness landscapes (*b*) and costs in males (*c*_m_) on the x and y axes respectively. Panel columns show plots for different strengths of asymmetry in male and female selection

**Supplementary Figure 2:**
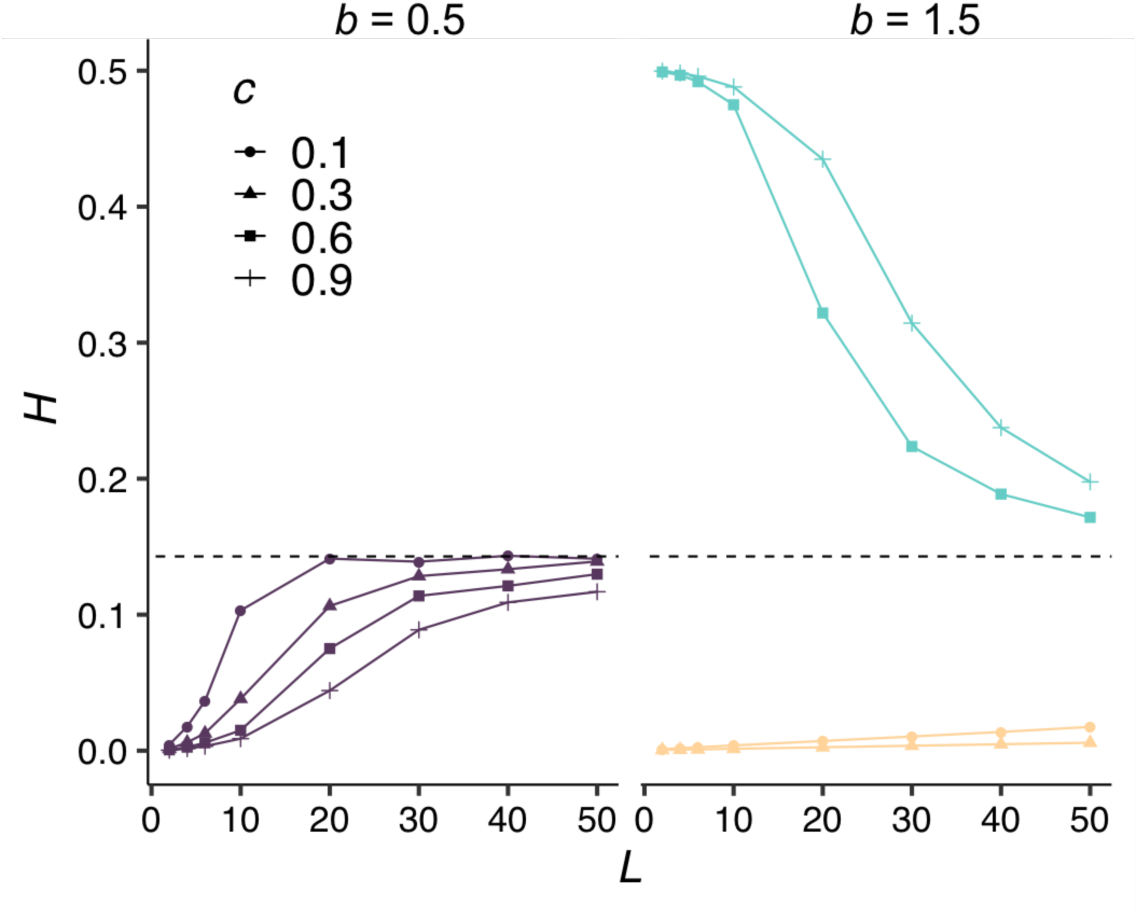
Effect of number of loci on heterozygosity in polygenic simulations. Curves show average heterozygosity across *L* ∈ [2, 50] di-allelic loci encoding *z* from simulations of our polygenic di-alleleic model (Appendix B.3 for simulation details, other parameters used: *δ =* 1, *µ =* 10^−5^). Different curve/point combinations represent different strength of sexually antagonistic selection (*c = c*_m_ *= c*_f_ *2* [0.1, 0.9]), left-hand panel shows results for diminishing fitness landscapes (*b =* 0.5 *<* 1) and right-hand panel for accelerating fitness landscapes (*b =* 1.5 *>* 1). Horizontal dashed line shows expected heterozygosity for a neutral locus at mutation-drift balance.

**Supplementary Figure 3:**
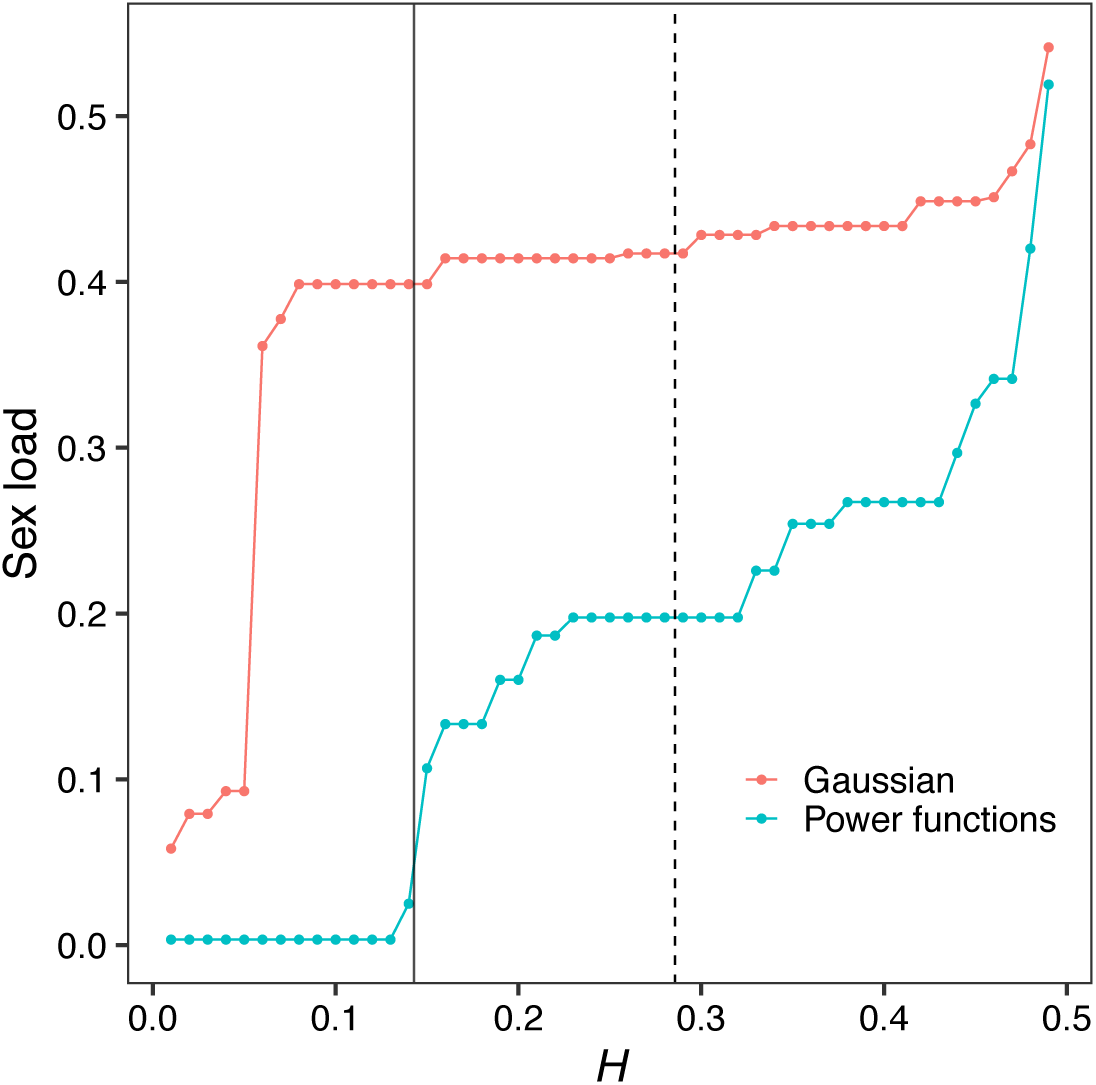
Minimum sex load produced from sexual antagonism leading to different levels of heterozygosity in individual based simulations. Dots show the minimum sex load, 1 *−* [*w*_f_(*z̄*_f_) *+ w*_m_(*z̄*_m_)]/2, where *z̄*_f_ and *z̄*_m_ are the mean male and female trait values in the population, produced by parameter combinations that lead to a given level of heterozygosity (*H*) in individual-based simulations of our di-allelic polygenic model with 10 loci (*L =* 10). Tourquise and red dots refer to simulations using Power (Fig. 3) and Gaussian fecundity functions (Appendix Fig. 7), respectively, and solid vertical line shows expected heterozygosity at neutrality while dotted vertical line shows twice the neutral expectation.

**Supplementary Figure 4:**
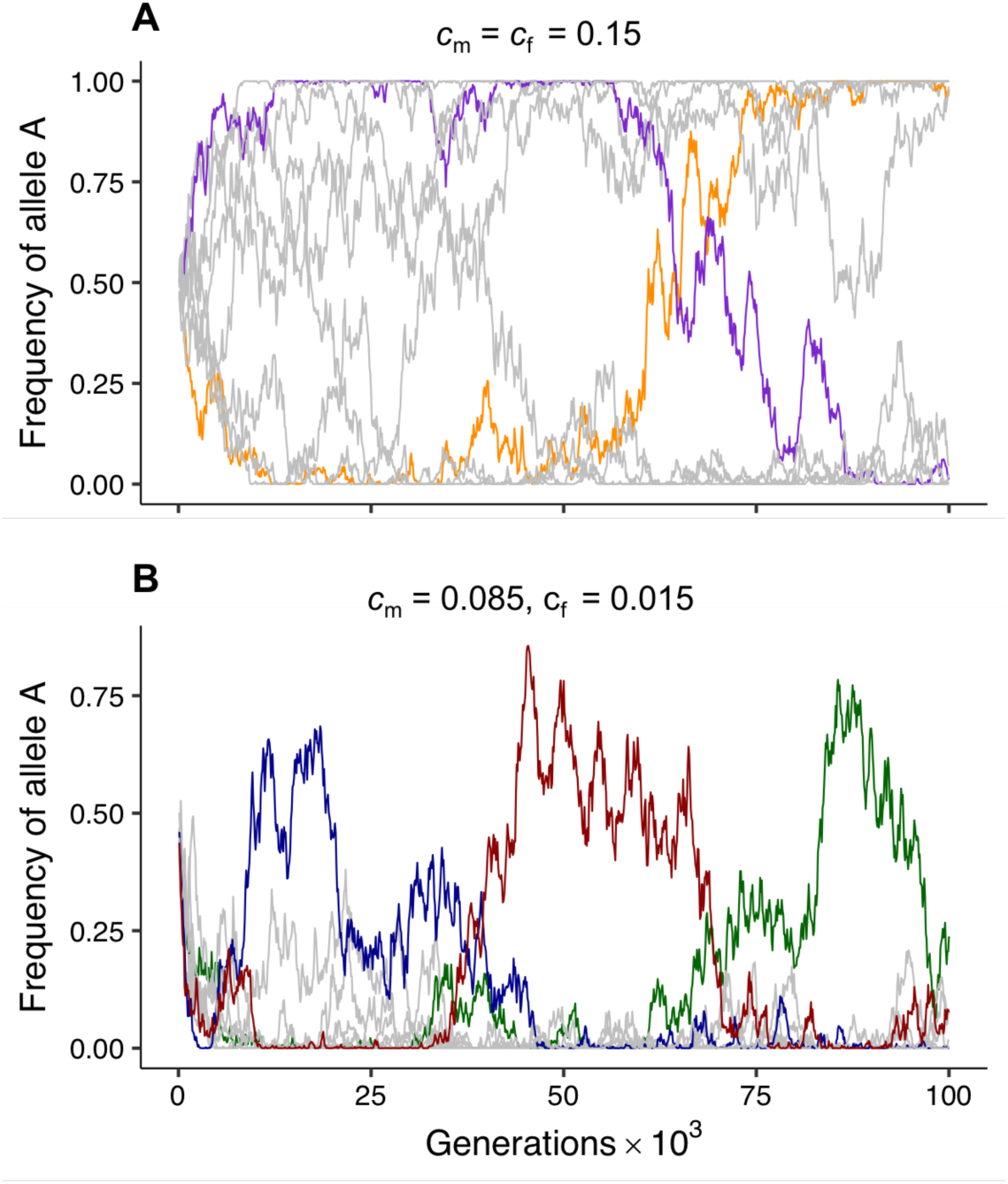
Allele frequencies through time under stabilising selection. Lines show frequency of the female-beneficial allele A at each locus through time in simulations of our di-allelic polygenic model when male and female fecundity follow power functions (eq. I.A, Box I, see Appendix E for full simulation procedures and calculations, parameters used: *L =* 10, *δ =* 1, *µ =* 10^−5^). Panel **A** shows a case with a symmetric fecundity trade-off across the sexes, whereby selection favours the fixation of equal numbers of A and a alleles across loci. Orange and purple lines show an example of two loci that are initially fixed for one allele but exhibit a transient genetic polymorphism by switching to become fixed for the opposing one (due to the combined effects of selection mutation and drift). Panel **B** shows a case of male-biased selection that leads selection to favour polymorphism at a single remainder locus. Dark blue, red, and green, lines showing three different loci that rise to high frequency due to turnover in the identity of the remainder locus owing to mutation and drift.

## Notes

### Competing Interest Statement

The authors have declared no competing interest.

### Summary of Updates

We have added text to clarify the connection between our results and previous theory.

## References

[1] Bonduriansky R, Chenoweth SF. 2009 Intralocus sexual conflict. Trends in ecology & evolution 24, 280–288.

[2] Cox RM, Calsbeek R. 2009 Sexually antagonistic selection, sexual dimorphism, and the resolution of intralocus sexual conflict. The American Naturalist 173, 176–187.

[3] Poissant J, Wilson AJ, Coltman DW. 2010 Sex-specific genetic variance and the evolution of sexual dimorphism: a systematic review of cross-sex genetic correlations. Evolution 64, 97–107.

[4] Parker G et al.. 1979 Sexual selection and sexual conflict. Sexual selection and reproductive competition in insects 123, 166.

[5] Rice W, Chippindale A. 2001 Intersexual ontogenetic conflict. Journal of Evolutionary Biology 14, 685–693.

[6] Chapman T, Arnqvist G, Bangham J, Rowe L. 2003 Sexual conflict. Trends in Ecology & Evolution 18, 41–47.

[7] Arnqvist G, Rowe L. 2005 Sexual Conflict vol. 31. Princeton University Press.

[8] Van Doorn GS. 2009 Intralocus sexual conflict. Annals of the New York Academy of Sciences 1168, 52–71.

[9] Connallon T, Clark AG. 2014a Evolutionary inevitability of sexual antagonism. Proceedings of the Royal Society B: Biological Sciences 281, 20132123.

[10] Connallon T, Clark AG. 2014b Balancing selection in species with separate sexes: insights from Fisher’s geometric model. Genetics 197, 991–1006.

[11] Muralidhar P, Coop G. 2023 Polygenic outcomes of sexually antagonistic selection. bioRxiv pp. 2023–03.

[12] Lande R. 1980 Sexual dimorphism, sexual selection, and adaptation in polygenic characters. Evolution 34, 292–305.

[13] Mank JE. 2017 Population genetics of sexual conflict in the genomic era. Nature Reviews Genetics 18, 721–730.

[14] Foerster K, Coulson T, Sheldon BC, Pemberton JM, Clutton-Brock TH, Kruuk LE. 2007 Sexually antagonistic genetic variation for fitness in red deer. Nature 447, 1107.

[15] Ruzicka F, Hill MS, Pennell TM, Flis I, Ingleby FC, Mott R, Fowler K, Morrow EH, Reuter M. 2019 Genome-wide sexually antagonistic variants reveal long-standing constraints on sexual dimorphism in fruit flies. PLoS biology 17, e3000244.

[16] Eyer PA, Blumenfeld AJ, Vargo EL. 2019 Sexually antagonistic selection promotes genetic divergence between males and females in an ant. Proceedings of the National Academy of Sciences 116, 24157–24163.

[17] Owen A. 1953 A genetical system admitting of two distinct stable equilibria under natural selection. Heredity 7, 97.

[18] Kidwell J, Clegg M, Stewart F, Prout T. 1977 Regions of stable equilibria for models of differential selection in the two sexes under random mating. Genetics 85, 171–183.

[19] Rice WR. 1984 Sex chromosomes and the evolution of sexual dimorphism. Evolution 38, 735– 742.

[20] Albert AY, Otto SP. 2005 Sexual selection can resolve sex-linked sexual antagonism. Science 310, 119–121.

[21] Gavrilets S, Rice WR. 2006 Genetic models of homosexuality: generating testable predictions. Proceedings of the Royal Society B: Biological Sciences 273, 3031–3038.

[22] Patten MM, Haig D. 2009 Maintenance or loss of genetic variation under sexual and parental antagonism at a sex-linked locus. Evolution 63, 2888–2895.

[23] Fry JD. 2010 The genomic location of sexually antagonistic variation: some cautionary comments. Evolution: International Journal of Organic Evolution 64, 1510–1516.

[24] Patten MM, Haig D, Ubeda F. 2010 Fitness variation due to sexual antagonism and linkage disequilibrium. Evolution 64, 3638–3642.

[25] Ubeda F, Haig D, Patten MM. 2011 Stable linkage disequilibrium owing to sexual antagonism. Proceedings of the Royal Society B: Biological Sciences 278, 855–862.

[26] Connallon T, Clark AG. 2012 A general population genetic framework for antagonistic selection that accounts for demography and recurrent mutation. Genetics 190, 1477–1489.

[27] Mullon C, Pomiankowski A, Reuter M. 2012 The effects of selection and genetic drift on the genomic distribution of sexually antagonistic alleles. Evolution: International Journal of Organic Evolution 66, 3743–3753.

[28] Jaquiéry J, Rispe C, Roze D, Legeai F, Le Trionnaire G, Stoeckel S, Mieuzet L, Da Silva C, Poulain J, Prunier-Leterme N, Ségurens B, Tagu D, Simon JC. 2013 Masculinization of the X Chromosome in the Pea Aphid. PLoS Genetics 9, e1003690–15.

[29] Jordan CY, Connallon T. 2014 Sexually antagonistic polymorphism in simultaneous hermaphrodites. Evolution 68, 3555–3569.

[30] Arnqvist G, Vellnow N, Rowe L. 2014 The effect of epistasis on sexually antagonistic genetic variation. Proceedings of the Royal Society B: Biological Sciences 281, 20140489.

[31] Spencer HG, Priest NK. 2016 The evolution of sex-specific dominance in response to sexually antagonistic selection. The American Naturalist 187, 658–666.

[32] Hill MS, Reuter M, Stewart AJ. 2019 Sexual antagonism drives the displacement of polymorphism across gene regulatory cascades. Proceedings of the Royal Society B 286, 20190660.

[33] Otto SP. 2019 Evolutionary potential for genomic islands of sexual divergence on recombining sex chromosomes. New Phytologist 224, 1241–1251.

[34] Ruzicka F, Connallon T. 2020 Is the X chromosome a hot spot for sexually antagonistic polymorphisms? Biases in current empirical tests of classical theory. Proceedings of the Royal Society B 287, 20201869.

[35] Hitchcock TJ, Gardner A. 2020 A gene’s-eye view of sexual antagonism. Proceedings of the Royal Society B 287, 20201633.

[36] Flintham EO, Savolainen V, Mullon C. 2021 Dispersal alters the nature and scope of sexually antagonistic variation. The American Naturalist 197, 543–559.

[37] Connallon T, Chenoweth SF. 2019 Dominance reversals and the maintenance of genetic variation for fitness. PLoS biology 17, e3000118.

[38] Grieshop K, Arnqvist G. 2018 Sex-specific dominance reversal of genetic variation for fitness. PLoS Biology 16, e2006810.

[39] Reid JM. 2022 Intrinsic emergence and modulation of sex-specific dominance reversals in threshold traits. Evolution 76, 1924–1941.

[40] Grieshop K, Ho EK, Kasimatis KR. 2024 Dominance reversals: the resolution of genetic conflict and maintenance of genetic variation. Proceedings of the Royal Society B 291, 20232816.

[41] Kaufmann P, Howie JM, Immonen E. 2023 Sexually antagonistic selection maintains genetic variance when sexual dimorphism evolves. Proceedings of the Royal Society B 290, 20222484.

[42] Stearns SC, Govindaraju DR, Ewbank D, Byars SG. 2012 Constraints on the coevolution of contemporary human males and females. Proceedings of the Royal Society B: Biological Sciences 279, 4836–4844.

[43] Khramtsova EA, Davis LK, Stranger BE. 2019 The role of sex in the genomics of human complex traits. Nature Reviews Genetics 20, 173–190.

[44] Charlesworth B, Charlesworth D. 2010 Elements of Evolutionary Genetics. Roberts and Company Publishers.

[45] Walsh B, Lynch M. 2018 Evolution and selection of quantitative traits. Oxford University Press.

[46] Turelli M, Barton N. 2004 Polygenic variation maintained by balancing selection: pleiotropy, sex-dependent allelic effects and G*£* E interactions. Genetics 166, 1053–1079.

[47] Connallon T, Matthews G. 2019 Cross-sex genetic correlations for fitness and fitness components: connecting theoretical predictions to empirical patterns. Evolution letters 3, 254–262.

[48] Matthews G, Hangartner S, Chapple DG, Connallon T. 2019 Quantifying maladaptation during the evolution of sexual dimorphism. Proceedings of the Royal Society B 286, 20191372.

[49] Haldane J. 1924 A mathematical theory of natural and artificial selection. Trans Camb Phil Soc p. 19.

[50] Bodmer WF. 1965 Differential fertility in population genetics models. Genetics 51, 411.

[51] Kimura M. 1965 A stochastic model concerning the maintenance of genetic variability in quantitative characters.. Proceedings of the National Academy of Sciences 54, 731–736.

[52] Otto SP, Day T. 2011 A Biologist’s Guide to Mathematical Modeling in Ecology and Evolution. Princeton University Press.

[53] Connallon T, Clark AG. 2010 Sex linkage, sex-specific selection, and the role of recombination in the evolution of sexually dimorphic gene expression. Evolution: International Journal of Organic Evolution 64, 3417–3442.

[54] Metz J, Geritz S, Meszena G, Jacobs F, Van Heerwaarden J. 1996 Adaptive dynamics: a geometrical study of the consequences of nearly faithful reproduction. In Stochastic and spatial structures of dynamical systems pp. 183–231. North-Holland.

[55] Dercole F, Rinaldi S. 2008 Analysis of evolutionary processes. Princeton University Press.

[56] Avila P, Mullon C. 2023 Evolutionary game theory and the adaptive dynamics approach: adaptation where individuals interact. Philosophical Transactions of the Royal Society B 378, 20210502.

[57] Eshel I. 1983 Evolutionary and continuous stability. Journal of theoretical Biology 103, 99–111.

[58] Christiansen FB. 1991 On conditions for evolutionary stability for a continuously varying character. The American Naturalist 138, 37–50.

[59] Geritz SA, Kisdi E, Metz JA et al.. 1998 Evolutionarily singular strategies and the adaptive growth and branching of the evolutionary tree. Evolutionary ecology 12, 35–57.

[60] Geritz SA, Kisdi É. 2000 Adaptive dynamics in diploid, sexual populations and the evolution of reproductive isolation. Proceedings of the Royal Society of London. Series B: Biological Sciences 267, 1671–1678.

[61] Rueffler C, Van Dooren TJ, Leimar O, Abrams PA. 2006 Disruptive selection and then what?. Trends in Ecology & Evolution 21, 238–245.

[62] Lande R, Arnold SJ. 1983 The measurement of selection on correlated characters. Evolution pp. 1210–1226.

[63] Jordan CY, Connallon T. 2014 Sexually antagonistic polymorphism in simultaneous hermaphrodites. Evolution 68, 3555–3569.

[64] Haller BC, Messer PW. 2023 SLiM 4: multispecies eco-evolutionary modeling. The American Naturalist 201, E127–E139.

[65] Bulmer M. 1971 The effect of selection on genetic variability. The American Naturalist 105, 201–211.

[66] Bürger R, Gimelfarb A. 1999 Genetic variation maintained in multilocus models of additive quantitative traits under stabilizing selection. Genetics 152, 807–820.

[67] Jordan CY, Charlesworth D. 2012 The potential for sexually antagonistic polymorphism in different genome regions. Evolution: International Journal of Organic Evolution 66, 505–516.

[68] Otto SP, Pannell JR, Peichel CL, Ashman TL, Charlesworth D, Chippindale AK, Delph LF, Guerrero RF, Scarpino SV, McAllister BF. 2011 About PAR: the distinct evolutionary dynamics of the pseudoautosomal region. Trends in Genetics 27, 358–367.

[69] Charlesworth B, Jordan CY, Charlesworth D. 2014 The evolutionary dynamics of sexually antagonistic mutations in pseudoautosomal regions of sex chromosomes. Evolution 68, 1339–1350.

[70] Lucotte EA, Laurent R, Heyer E, Segurel L, Toupance B. 2016 Detection of allelic frequency differences between the sexes in humans: a signature of sexually antagonistic selection. Genome Biology and Evolution 8, 1489–1500.

[71] Cheng C, Kirkpatrick M. 2016 Sex-specific selection and sex-biased gene expression in humans and flies. PLoS Genetics 12.

[72] Kasimatis KR, Ralph PL, Phillips PC. 2019 Limits to genomic divergence under sexually antagonistic selection. G3: Genes, Genomes, Genetics 9, 3813–3824.

[73] Ruzicka F, Dutoit L, Czuppon P, Jordan CY, Li XY, Olito C, Runemark A, Svensson EI, Yazdi HP, Connallon T. 2020 The search for sexually antagonistic genes: Practical insights from studies of local adaptation and statistical genomics. Evolution letters 4, 398–415.

[74] Ruzicka F, Holman L, Connallon T. 2022 Polygenic signals of sex differences in selection in humans from the UK Biobank. PLoS Biology 20, e3001768.

[75] Berger D, Grieshop K, Lind MI, Goenaga J, Maklakov AA, Arnqvist G. 2014 Intralocus sexual conflict and environmental stress. Evolution 68, 2184–2196.

[76] Kasimatis KR, Nelson TC, Phillips PC. 2017 Genomic signatures of sexual conflict. Journal of Heredity 108, 780–790.

[77] Kasimatis KR, Abraham A, Ralph PL, Kern AD, Capra JA, Phillips PC. 2020 Sexually Antagonistic Selection on Genetic Variation is Rare in Humans. bioRxiv.

[78] Singh A, Punzalan D. 2018 The strength of sex-specific selection in the wild. Evolution 72, 2818–2824.

[79] Sanjak JS, Sidorenko J, Robinson MR, Thornton KR, Visscher PM. 2018 Evidence of directional and stabilizing selection in contemporary humans. Proceedings of the National Academy of Sciences 115, 151–156.

[80] Mank JE. 2017 The transcriptional architecture of phenotypic dimorphism. Nature Ecology & Evolution 1, 1–7.

[81] Meszéna G, Czibula I, Geritz SAH. 1997 Adaptive dynamics in a 2-patch environment: A toy model for allopatric and parapatric specation. Journal of Biological Systems 5, 265–284.

[82] Lythgoe KA. 1997 Consequences of gene flow in spatially structured populations. Genetics Research 69, 49–60.

[83] Spichtig M, Kawecki TJ. 2004 The maintenance (or not) of polygenic variation by soft selection in heterogeneous environments. The American Naturalist 164, 70–84.

[84] Doorn GSV, Dieckmann U. 2006 The long-term evolution of multilocus traits under frequencydependent disruptive selection. Evolution 60, 2226–2238.

[85] Svardal H, Rueffler C, Hermisson J. 2015 A general condition for adaptive genetic polymorphism in temporally and spatially heterogeneous environments. Theoretical Population Biology 99, 76– 97.

[86] Débarre F, Gandon S. 2011 Evolution in heterogeneous environments: between soft and hard selection. The American Naturalist 177, E84–E97.

[87] Connallon T, Débarre F, Li XY. 2018 Linking local adaptation with the evolution of sex differences. Proceedings of the Royal Society B.

[88] Sellis D, Callahan BJ, Petrov DA, Messer PW. 2011 Heterozygote advantage as a natural consequence of adaptation in diploids. Proceedings of the National Academy of Sciences 108, 20666– 20671.

[89] Lewis Z, Wedell N, Hunt J. 2011 Evidence for strong intralocus sexual conflict in the Indian meal moth, Plodia interpunctella. Evolution 65, 2085–2097.

[90] Tarka M, Åkesson M, Hasselquist D, Hansson B. 2014 Intralocus sexual conflict over wing length in a wild migratory bird. The American Naturalist 183, 62–73.

[91] Kaufmann P, Wolak ME, Husby A, Immonen E. 2021 Rapid evolution of sexual size dimorphism facilitated by Y-linked genetic variance. Nature Ecology & Evolution 5, 1394–1402.

[92] Zhu C, Ming MJ, Cole JM, Edge MD, Kirkpatrick M, Harpak A. 2023 Amplification is the primary mode of gene-by-sex interaction in complex human traits. Cell Genomics 3.

[93] Sylvestre F, Mérot C, Normandeau E, Bernatchez L. 2023 Searching for intralocus sexual conflicts in the three-spined stickleback (Gasterosteus aculeatus) genome. Evolution 77, 1667–1681.

[94] Barson NJ, Aykanat T, Hindar K, Baranski M, Bolstad GH, Fiske P, Jacq C, Jensen AJ, Johnston SE, Karlsson S et al.. 2015 Sex-dependent dominance at a single locus maintains variation in age at maturity in salmon. Nature 528, 405.

[95] Rusuwa BB, Chung H, Allen SL, Frentiu FD, Chenoweth SF. 2022 Natural variation at a single gene generates sexual antagonism across fitness components in Drosophila. Current Biology 32, 3161–3169.

[96] Rowe L, Chenoweth SF, Agrawal AF. 2018 The genomics of sexual conflict. The American Naturalist 192, 274–286.

