## Supplementary File for "The maintenance of genetic polymorphism underlying sexually antagonistic traits"

#### A Continuum-of-alleles model

##### A.1 Population genetic recursions

As a basis to our analysis of the continuum-of-alleles model, we consider one autosomal locus where a new allele A appears in a population fixed for allele a. To determine whether A invades or not, we first write recursion equations for the frequencies of these alleles in male and female gametes at generation $t+1$ as a function of their frequencies in generation $t$. Selection influences allele frequencies through the fecundities of adults carrying the three diploid genotypes AA, Aa, aa and expressing the respective phenotypes $z_{\text{AA}}$, $z_{\text{Aa}}$, $z_{\text{aa}}$ (i.e., where $z_u$ is phenotype encoded by genotype $u \in \{\text{AA}, \text{Aa}, \text{aa}\}$). As a result, the frequency, $p_{v,t+1}$, of A in the pool of gametes produced by sex $v \in \{\text{m}, \text{f}\}$ at generation

$t+1$ is given by

$$\begin{aligned} p_{\text{m},t+1} &= M(p_{\text{m},t}, p_{\text{f},t}) \\ p_{\text{f},t+1} &= F(p_{\text{m},t}, p_{\text{f},t}), \end{aligned} \tag{A-1}$$

where

$$\begin{aligned} M(p_{\text{m},t}, p_{\text{f},t}) &= p_{\text{m},t}p_{\text{f},t}\frac{w_{\text{m}}(z_{\text{AA}})}{\overline{w}_{\text{m},t}} + \frac{1}{2}\big(p_{\text{m},t}(1-p_{\text{f},t}) + (1-p_{\text{m},t})p_{\text{f},t}\big)\frac{w_{\text{m}}(z_{\text{Aa}})}{\overline{w}_{\text{m},t}} \\ F(p_{\text{m},t}, p_{\text{f},t}) &= p_{\text{m},t}p_{\text{f},t}\frac{w_{\text{f}}(z_{\text{AA}})}{\overline{w}_{\text{f},t}} + \frac{1}{2}\big(p_{\text{m},t}(1-p_{\text{f},t}) + (1-p_{\text{m},t})p_{\text{f},t}\big)\frac{w_{\text{f}}(z_{\text{Aa}})}{\overline{w}_{\text{f},t}}, \end{aligned} \tag{A-2}$$

and $w_v(z_u)$ is the number of sperm or eggs produced an individual of the genotype $u \in \{\text{AA}, \text{Aa}, \text{aa}\}$ expressing the phenotype $z_u$ in sex $v$ (e.g., eq. I.A for power functions), and $\overline{w}_{\text{m},t} = p_{\text{m},t}p_{\text{f},t}w_{\text{m}}(z_{\text{AA}}) + \big(p_{\text{m},t}(1-p_{\text{f},t}) + (1-p_{\text{m},t})p_{\text{f},t}\big)w_{\text{m}}(z_{\text{Aa}}) + (1-p_{\text{m},t})(1-p_{\text{f},t})w_{\text{m}}(z_{\text{aa}})$ and $\overline{w}_{\text{f},t} = p_{\text{m},t}p_{\text{f},t}w_{\text{f}}(z_{\text{AA}}) + \big(p_{\text{m},t}(1-p_{\text{f},t}) + (1-p_{\text{m},t})p_{\text{f},t}\big)w_{\text{f}}(z_{\text{Aa}}) + (1-p_{\text{m},t})(1-p_{\text{f},t})w_{\text{f}}(z_{\text{aa}})$ are the mean male and female fecundities at generation $t$, respectively.

Allele A can invade when rare (i.e., when $p_{\text{m},t}$ and $p_{\text{f},t}$ are small) when the leading eigenvalue, $\lambda(0)$, of the Jacobian matrix,

$$\text{J}(0) = \begin{pmatrix} \dfrac{\partial M(p_{\text{m},t}, p_{\text{f},t})}{\partial p_{\text{m},t}} & \dfrac{\partial M(p_{\text{m},t}, p_{\text{f},t})}{\partial p_{\text{f},t}} \\ \dfrac{\partial F(p_{\text{m},t}, p_{\text{f},t})}{\partial p_{\text{m},t}} & \dfrac{\partial F(p_{\text{m},t}, p_{\text{f},t})}{\partial p_{\text{f},t}} \end{pmatrix}_{p_{\text{m},t}=p_{\text{f},t}=0}, \tag{A-3}$$

is greater than unity ($\lambda(0) > 1$). This leading eigenvalue equals:

$$\lambda(0) = \frac{1}{2}\frac{w_{\text{m}}(z_{\text{Aa}})}{w_{\text{m}}(z_{\text{aa}})} + \frac{1}{2}\frac{w_{\text{f}}(z_{\text{Aa}})}{w_{\text{f}}(z_{\text{aa}})}, \tag{A-4}$$

for any fitness functions in males and females, both for strong and weak selection, as long as the resident population consists only of a single allele, a.

#### A.2 Invasion fitness and adaptive dynamics

Here, we analyse evolutionary dynamics under the continuum-of-alleles model in diploids, following, e.g., [1]. As a basis to such an analysis, we need the invasion fitness (i.e., geometric growth rate), $W(x_\bullet, x)$, of a rare mutant allele that encodes an additive phenotypic effect $x_\bullet$ in a population that is otherwise fixed for an allele encoding an effect $x$. From the above (Appendix A.1), invasion of such a mutant allele is determined by eq. (A-4) with mutant heterozygote phenotype $z_{\mathrm{Aa}} = x_\bullet + x$ and resident homozygote phenotype $z_{\mathrm{aa}} = 2x$. Accordingly, invasion fitness is given by,

$$W(x_\bullet, x) = \frac{1}{2}\frac{w_{\mathrm{m}}(x_\bullet + x)}{w_{\mathrm{m}}(2x)} + \frac{1}{2}\frac{w_{\mathrm{f}}(x_\bullet + x)}{w_{\mathrm{f}}(2x)}. \tag{A-5}$$

Given a small change in phenotype (so that $x_\bullet$ is close to $x$), the selection gradient determines the form of directional selection:

$$S(x) = \left.\frac{\partial W(x_\bullet, x)}{\partial x_\bullet}\right|_{x_\bullet = x}, \tag{A-6}$$

i.e., $S(x)$ indicates whether in a population fixed for $x$, selection favours an increase in allelic value when $S(x) > 0$ or a decrease when $S(x) < 0$. A singular allelic value $x^*$ is such that when expressed by the whole population, there is no directional selection, i.e., such that

$$S(x^*) = 0. \tag{A-7}$$

Whether evolutionary dynamics converge to such a singularity depends on the sign of

$$S'(x^*) = \left.\frac{dS(x)}{dx}\right|_{x = x^*}. \tag{A-8}$$

If $S'(x^*) < 0$, then $x^*$ is an attractor of evolutionary dynamics, i.e., it is convergence stable [2], so that owing to directional selection and mutational input, the population will gradually converge to $x^*$ and thus express the phenotype $z^* = 2x^*$. By contrast, if $S'(x^*) > 0$, then $x^*$ is a repellor of evolutionary dynamics so that the population will gradually converge away from $x^*$.

If a population converges to a singular allelic value $x^*$, evolutionary dynamics are then determined by quadratic selection, i.e., by the second order fitness effect, which is given by

$$H(x^*) = \left.\frac{\partial^2 W(x_\bullet, x)}{\partial x_\bullet^2}\right|_{x_\bullet = x = x^*}. \tag{A-9}$$

When $H(x^*) < 0$, selection is stabilising [3], selecting against any mutant that deviates from $x^*$, i.e., $x^*$ is uninvadable [4]. When $H(x^*) > 0$, selection is diversifying (or negatively frequency-dependent and disruptive), favouring the emergence of polymorphism. The distribution of allelic values in the population thus goes from being unimodal to bimodal in a process referred to as evolutionary branching [5, 6].

#### A.3 General fecundity functions

For arbitrary male and female fecundity $w_{\mathrm{m}}(x)$ and $w_{\mathrm{f}}(x)$, we find that the selection gradient (eq. A-6) is given by

$$S(x) = \frac{1}{2}\left[\frac{w'_{\mathrm{f}}(2x)}{w_{\mathrm{f}}(2x)} + \frac{w'_{\mathrm{m}}(2x)}{w_{\mathrm{m}}(2x)}\right]. \tag{A-10}$$

This shows that allelic value $x^*$ is singular if

$$\frac{w'_{\mathrm{f}}(2x^*)}{w_{\mathrm{f}}(2x^*)} = -\frac{w'_{\mathrm{m}}(2x^*)}{w_{\mathrm{m}}(2x^*)}, \tag{A-11}$$

so that the relative fitness gain in one sex is balanced by the relative fitness loss in the other. Convergence stability (eq. A-8) of $x^*$ depends on the sign of:

$$S'(x^*) = \left[\frac{w''_{\mathrm{f}}(2x^*)}{w_{\mathrm{f}}(2x^*)} + \frac{w''_{\mathrm{m}}(2x^*)}{w_{\mathrm{m}}(2x^*)}\right] - \left\{\left(\frac{w'_{\mathrm{f}}(2x^*)}{w_{\mathrm{f}}(2x^*)}\right)^2 + \left(\frac{w'_{\mathrm{m}}(2x^*)}{w_{\mathrm{m}}(2x^*)}\right)^2\right\}, \tag{A-12}$$

while the existence of stabilizing vs. diversifying selection depends on (eq. A-9):

$$H(x^*) = \frac{1}{2}\left[\frac{w''_{\mathrm{f}}(2x^*)}{w_{\mathrm{f}}(2x^*)} + \frac{w''_{\mathrm{m}}(2x^*)}{w_{\mathrm{m}}(2x^*)}\right]. \tag{A-13}$$

For there to be diversifying selection, we must have $S'(x^*) < 0$ and $H(x^*) > 0$, which requires

$$0 < \left[\frac{w''_{\mathrm{f}}(2x^*)}{w_{\mathrm{f}}(2x^*)} + \frac{w''_{\mathrm{m}}(2x^*)}{w_{\mathrm{m}}(2x^*)}\right] < \left\{\left(\frac{w'_{\mathrm{f}}(2x^*)}{w_{\mathrm{f}}(2x^*)}\right)^2 + \left(\frac{w'_{\mathrm{m}}(2x^*)}{w_{\mathrm{m}}(2x^*)}\right)^2\right\} \tag{A-14}$$

The right-hand term in curly braces represents the strength of directional selection squared in both sexes and so is negligible when selection is weak. Thus, whatever the fecundity functions $w_{\mathrm{m}}(z)$ and $w_{\mathrm{f}}(z)$, selection can never be diversifying and polymorphism can never emerge from gradual evolution when variation in male and female fecundity ($w_{\mathrm{m}}(z)$ and $w_{\mathrm{f}}(z)$) is small. Strong selection can, however, allow for diversifying selection, but only when the fitness surfaces are positively curved, on average across the sexes (first condition), but not so curved that the second condition fails.

Equations (A-11)-(A-14) are given in terms of $z = 2x$ in main text eq. (4a)-(4b).

#### A.4 Power functions

##### A.4.1 Equal fecundity costs in males and females

We now consider the specific power fitness functions eq. (I.A) in Box I, assuming initially that fecundity declines with the same intensity away from the male and female optima ($c = c_{\text{m}} = c_{\text{f}}$) as this lends itself better to mathematical scrutiny. Plugging eq. (I.A) into eq. (A-5) and setting $c = c_{\text{m}} = c_{\text{f}}$, we find that invasion fitness reduces to

$$W(x_\bullet, x) = \frac{1}{2} \cdot \frac{1 - c\left[1 + 2^{-b}\left(\frac{\theta - x - x_\bullet}{\theta}\right)^b\right]}{1 - c\left[1 + \left(\frac{1}{2} - \frac{x}{\theta}\right)^b\right]} + \frac{1}{2} \cdot \frac{1 - c\left[1 + 2^{-b}\left(\frac{\theta + x + x_\bullet}{\theta}\right)^b\right]}{1 - c\left[1 + \left(\frac{1}{2} + \frac{x}{\theta}\right)^b\right]}. \tag{A-15}$$

In turn, plugging eq. (A-15) into (A-6) we obtain the selection gradient

$$S(x) = \frac{1}{2} bc \left[ \frac{1}{c + (1-c)\left(\frac{1}{2} + \frac{x}{\theta}\right)^{-b}(2x - \theta)} + \frac{1}{c + (1-c)\left(\frac{1}{2} - \frac{x}{\theta}\right)^{-b}(2x + \theta)} \right], \tag{A-16}$$

from which it can be seen that $x^* = 0$ is always a singular point (i.e., $S(0) = 0$). Substituting eq. (A-16) into eq. (A-16), and evaluating the result at $x^* = 0$ yields

$$S'(0) = \frac{2cb\left[2^b(b-1)(1-c) - c\right]}{\left[2^b + c(1 - 2^b)\right]^2}. \tag{A-17}$$

This show that $S'(0) < 0$ and $x^* = 0$ is an attractor of selection, either when

$$b < 1 \tag{A-18}$$

(i.e., fecundity curves are diminishing), or when

$$\begin{cases} b > 1 \\ c > \dfrac{1}{1 + \frac{1}{2^b(b-1)}} \end{cases} \tag{A-19}$$

(i.e., curves are accelerating and selection is sufficiently strong). Otherwise, $S'(0) > 0$ and $x^* = 0$ is a repellor, with selection being directional towards $x = \theta$ if the population initially is fixed for an allelic value above 0 ($x > 0$) or $x = -\theta$ if initially below 0 ($x < 0$).

Plugging eq. (A-15) into (A-9) and evaluating the result at $x^* = 0$ gives us

$$H(0) = \frac{cb(b-1)}{2^b(1-c)+c}. \tag{A-20}$$

This shows that whenever $b < 1$, the singular trait value $x^* = 0$ is under stabilising selection (as $H(0) < 0$). Otherwise, $H(0) > 0$ so that if eq. (A-19) also holds, $x^* = 0$ is an evolutionary branching point where selection is diversifying. These results are summarised in Fig. 2B.

##### A.4.2 Asymmetric fecundity costs

The case of power functions with asymmetric fecundity costs between the sexes ($c_\mathrm{m} \neq c_\mathrm{f}$) is more complicated. To gain some analytical insights, we first assume that selection is weak (i.e., that $c_\mathrm{m}$ and $c_\mathrm{f}$ are small).

**Weak selection** Substituting main text eq. (I.A) into eq. (A-5), mutant fitness can be expressed

$$W(x_\bullet, x) = \frac{1}{4}\left[2 - 2^{-b}c_\mathrm{m}\left(\left(1-\frac{2x}{\theta}\right)^b - \left(\frac{\theta - x_\bullet - x}{\theta}\right)^{b_\mathrm{m}}\right) - 2^{-b}c_\mathrm{f}\left(\left(1+\frac{2x}{\theta}\right)^b - \left(\frac{\theta + x_\bullet + x}{\theta}\right)^b\right)\right] + \mathcal{O}(\epsilon^2), \tag{A-21}$$

where $c_\mathrm{m} \sim \mathcal{O}(\epsilon)$, $c_\mathrm{f} \sim \mathcal{O}(\epsilon)$ and $0 < \epsilon \ll 1$ is a small parameter. Plugging eq. (A-21) into eq. (A-6), we then find the singular allelic value

$$x^* = \theta \frac{1 - (c_\mathrm{m}/c_\mathrm{f})^{1/(1-b)}}{2\left(1 + (c_\mathrm{m}/c_\mathrm{f})^{1/(1-b)}\right)} + \mathcal{O}(\epsilon^2). \tag{A-22}$$

The equilibrium phenotype, which is given by $z^* = 2x^*$, is shown in Fig. 2A.

Using our results for the stability of a singular strategy $x^*$ under general fitness functions (Appendix section A.3) assuming weak selection (so that the terms in curly braces in eq. A-12 are negligible) we find the following patterns. Whenever the curvature for the male and female fitness function are both negative (i.e., $b < 1$), the singular point will be convergence stable ($S'(x^*) < 0$) and subject to stabilizing selection ($H(x^*) < 0$, see eq. A-13). Conversely, when the curvature for the male and female fitness functions are both positive (i.e., $b > 1$), the singular point will be a repellor and not convergence stable ($S'(x^*) > 0$, $H(x^*) > 0$). Thus, with weak selection, there is either stabilising selection ($b < 1$) or directional selection ($b > 1$), but never diversifying selection (see also eq. A-14 for a more general argument).

**Arbitrarily strong selection** We analysed the general case ($c_\mathrm{m} \neq c_\mathrm{f}$) using a numerical approach. Specifically, we first solved numerically eq. (A-7) for $x^*$ and analysed its stability by evaluating eqs. (A-8)-(A-9). We did this for a range of parameter values within $c_\mathrm{m} \in [0,1]$, $b \in [0,2]$ and $\alpha \in [0,1]$ where $c_\mathrm{f} = \alpha c_\mathrm{m}$.

Our results are summarized in the top row panels of Supplementary Fig. 1. These show that selection is stabilising for$x^*$ whenever fecundity curves are diminishing ($b < 1$, Supplementary Fig. 1 purple region), that selection is diversifying when selection in males and females is strong and of similar intensity (Supplementary Fig. 1 green region), and that otherwise selection is directional towards $x = -\theta$ or $x = \theta$ depending on whether the population is initially below or above $x^*$ (Supplementary Fig. 1 yellow region).

##### A.4.3 Simulations

We ran individual-based simulations of the continuum-of-alleles model using SLiM 4 [7], whose results are presented in Fig. 2. Following the life cycle described in section 2, we consider a diploid population of 5,000 adult males and 5,000 adult females with non-overlapping generations. Each individual is characterised by the two allelic values it carries. At the beginning of the next generation, say $t+1$, 5,000 male and 5,000 female offspring are created, with the mother and father of each offspring being sampled from the adult females and males at generation $t$. Sampling is performed with replacement according to a multinomial distribution where the weighting on each male and female individual is according to their fecundity

$$\begin{aligned} w_\mathrm{m}(z) &= (1-c_\mathrm{m}) + c_\mathrm{m}\left(\frac{\theta - z}{2\theta}\right)^{b_\mathrm{m}} \\ w_\mathrm{f}(z) &= (1-c_\mathrm{f}) + c_\mathrm{f}\left(\frac{\theta + z}{2\theta}\right)^{b_\mathrm{f}}, \end{aligned} \tag{A-23}$$

(eq. I.A), where $z$ is the sum of the two allelic values carried. Each parent transmits one of the copies it carries, each with equal probability (i.e., no segregation bias). Mutations occur at a per-locus rate $\mu = 0.1$ with a new mutation adding a small deviate sampled from a Normal distribution $N(0,10^{-6})$ onto the existing allelic value at the mutating gene. After sampling, adults are removed from the simulations, juveniles become adults and the new generation begins.

### B Di-allelic model with one or more loci

In this appendix, we model trait $z$ as being encoded by $L$ autosomal di-allelic loci, with a male- and female-beneficial allele at each locus (when $L > 1$ this is the polygenic model described in the main text). We analyse this model mathematically for the cases $L = 1$ and $L = 2$ and with simulations for $L > 2$. Throughout we assume fecundity follows the power functions (eq. I.A in Box I). Our results show that polymorphism is more easily maintained where genetic constraints are extremely strong, that is where the trait is underlain by very few loci and thus where there are few alleles available.

#### B.1 Dynamics at a single locus

##### B.1.1 Reciprocal invasion

To connect to classical results [8], we first consider a trait encoded by a single di-allelic locus ($L = 1$), i.e., at which the alleles $A_1$ and $a_1$ (denoted A and a in this subsection) segregate. We assume that these alleles span the phenotypic range, so that the three diploid genotypes AA, Aa, aa express the phenotypes $z_{AA} = \theta$, $z_{Aa} = 0$, $z_{aa} = -\theta$. The recursion equations describing the change in frequency of these two alleles are given by eq. (A-1). Balancing selection at a locus occurs whenever each allele (A and a) can be invaded by the other (reciprocal invasion). First, allele A can invade a population fixed for a (i.e., where $p_{m,t} = p_{f,t} = 0$) when the leading eigenvalue, $\lambda(0)$, of the Jacobian matrix J(0) given by eq. (A-3) is greater than one. Similarly, allele a can invade a population fixed for A when the leading eigenvalue, $\lambda(1)$, of the Jacobian matrix J(1) is greater than one, where J(1) is given by the matrix eq. (A-3) evaluated now at $p_{m,t} = p_{f,t} = 1$.

Plugging eq. (I.A) into eq. (A-1) and eq. (A-1) into J(0) and J(1), we obtain a single condition for $\lambda(0) > 1$ and $\lambda(1) > 1$ to be satisfied,

$$\frac{2^b - 1}{1 + (2^b - 1)c_f} < c_m / c_f < \frac{1}{(2^b - 1)(1 - c_f)} \tag{B-1}$$

which corresponds to a classical condition for a protected polymorphism at a single sexually antagonistic locus (eq. 7 in [8] and eq. 3 in [9] where $h_m = h_f = 1 - 2^{-b}$). Inequality (B-1) is satisfied most readily when $b < 1$, as this favours an intermediate value of $z$ that is encoded by the heterozygote Aa, i.e., there is dominance reversal for fitness (Supplementary Fig. 1 middle row panels). This result differs substantially from the continuum-of alleles model (Supplementary Fig. 1 compare top and middle row panels), where polymorphism is never maintained with $b < 1$, and reflects the fact that we now constrain the system to have only two alleles, which straddle $x^* = 0$ (with phenotypes $\theta$ for AA and $-\theta$ for aa).

Polymorphism may also be maintained when $b > 1$ if sexual antagonism is strong ($c_\mathrm{m}$ and $c_\mathrm{m}$ both large and similar), although the conditions for this to occur are more permissive than in the continuum-of-alleles case (Supplementary Fig. 1 compare top and middle row panels). This is because the single locus model considers the segregation of two already highly differentiated alleles, A and a (rather than the gradual evolution of allelic values through mutations of small effect in the continuum-of-alleles model), which more easily allows for reciprocal invasibility.

#### B.2 Dynamics at two loci

Second we consider when $z$ is determined by two di-allelic loci (locus 1 where alleles $a_1$, $A_1$ segregate, and locus 2 where alleles $a_2$, $A_2$ segregate).

##### B.2.1 Recursions

To characterise allele frequency change at two loci, we now track the dynamics of the haploid male and female gamete genotypes [10–13] rather than gamete allele frequencies. We denote the sex-specific frequencies of each gametic genotype in generation $t$ (before syngamy) as $m_{1,t}, m_{2,t}, m_{3,t}, m_{4,t}$ in male gametes, and $f_{1,t}, f_{2,t}, f_{3,t}, f_{4,t}$ in female gametes, for the genotypes $A_1A_2$, $A_1a_2$, $a_1A_2$, and $a_1a_2$ respectively (where $\sum_{i=1}^4 m_{i,t} = \sum_{i=1}^4 f_{i,t} = 1$). Meanwhile, adult genotypes at generation $t$ are denoted $g^\mathrm{m}_{ij,t}$ in males and $g^\mathrm{f}_{ij,t}$ in females, where $i \in \{1,2,3,4\}$ and $j \in \{1,2,3,4\}$ respectively refer to the different paternally and maternally inherited haploid genotypes. Following the lifecycle in section 2, adult genotype frequencies in each sex depend on the sex-specific gametic frequencies according to

$$\begin{aligned} g^\mathrm{m}_{ij,t} &= f_{i,t}m_{j,t} \\ g^\mathrm{f}_{ij,t} &= f_{i,t}m_{j,t} \end{aligned} \tag{B-2}$$

in males and females, and from eqs. (2)-(3), the adult phenotypes are

$$
\begin{aligned}
&z_{11} = 4\delta \\
&z_{12} = z_{13} = 3\delta \\
&z_{14} = z_{22} = z_{22} = z_{23} = z_{33} = 0 \\
&z_{24} = z_{34} = -3\delta \\
&z_{44} = -4\delta,
\end{aligned}
\tag{B-3}
$$

where $z_{ij} = z_{ji}$ is the phenotype encoded by adult diploid genotype $ij$. We can therefore write recursion equations for the male and female gametes as

$$
\begin{aligned}
m_{1,t+1} &= M_1(\mathbf{g}_t^{\mathrm{m}}) \\
m_{2,t+1} &= M_2(\mathbf{g}_t^{\mathrm{m}}) \\
m_{3,t+1} &= M_3(\mathbf{g}_t^{\mathrm{m}}) \\
m_{4,t+1} &= M_4(\mathbf{g}_t^{\mathrm{m}}),
\end{aligned}
\tag{B-4}
$$

and

$$
\begin{aligned}
f_{1,t+1} &= F_1(\mathbf{g}_t^{\mathrm{f}}) \\
f_{2,t+1} &= F_2(\mathbf{g}_t^{\mathrm{f}}) \\
f_{3,t+1} &= F_3(\mathbf{g}_t^{\mathrm{f}}) \\
f_{4,t+1} &= F_4(\mathbf{g}_t^{\mathrm{f}}),
\end{aligned}
\tag{B-5}
$$

respectively, where $\mathbf{g}_t^{\mathrm{m}} = (g_{11,t}^{\mathrm{m}}, \dots, g_{44,t}^{\mathrm{m}})$ and $\mathbf{g}_t^{\mathrm{f}} = (g_{11,t}^{\mathrm{f}}, \dots, g_{44,t}^{\mathrm{f}})$ and where the functions $M_1(\mathbf{g}_t^{\mathrm{m}}), M_2(\mathbf{g}_t^{\mathrm{m}}), M_3(\mathbf{g}_t^{\mathrm{m}}), M_4(\mathbf{g}_t^{\mathrm{m}})$ and $F_1(\mathbf{g}_t^{\mathrm{f}}), F_2(\mathbf{g}_t^{\mathrm{f}}), F_3(\mathbf{g}_t^{\mathrm{f}}), F_4(\mathbf{g}_t^{\mathrm{f}})$ are

$$
\begin{aligned}
M_1(\mathbf{g}_t^{\mathrm{m}}) =& \frac{w_{\mathrm{m}}(z_{11})}{\overline{w}_{\mathrm{m},t}} g_{11,t}^{\mathrm{m}} + \frac{1}{2}\Big[ \frac{w_{\mathrm{m}}(z_{12})}{\overline{w}_{\mathrm{m},t}}(g_{12,t}^{\mathrm{m}} + g_{21,t}^{\mathrm{m}}) + \frac{w_{\mathrm{m}}(z_{13})}{\overline{w}_{\mathrm{m},t}}(g_{13,t}^{\mathrm{m}} + g_{31,t}^{\mathrm{m}}) + (1-r)\frac{w_{\mathrm{m}}(z_{14})}{\overline{w}_{\mathrm{m},t}}(g_{14,t}^{\mathrm{m}} + g_{41,t}^{\mathrm{m}}) + \\
& r\frac{w_{\mathrm{m}}(z_{23})}{\overline{w}_{\mathrm{m},t}}(g_{23,t}^{\mathrm{m}} + g_{32,t}^{\mathrm{m}})\Big] \\
M_2(\mathbf{g}_t^{\mathrm{m}}) =& \frac{w_{\mathrm{m}}(z_{22})}{\overline{w}_{\mathrm{m},t}} g_{22,t}^{\mathrm{m}} + \frac{1}{2}\Big[ \frac{w_{\mathrm{m}}(z_{12})}{\overline{w}_{\mathrm{m},t}}(g_{12,t}^{\mathrm{m}} + g_{21,t}^{\mathrm{m}}) + \frac{w_{\mathrm{m}}(z_{24})}{\overline{w}_{\mathrm{m},t}}(g_{24,t}^{\mathrm{m}} + g_{42,t}^{\mathrm{m}}) + (1-r)\frac{w_{\mathrm{m}}(z_{23})}{\overline{w}_{\mathrm{m},t}}(g_{23,t}^{\mathrm{m}} + g_{32,t}^{\mathrm{m}}) + \\
& r\frac{w_{\mathrm{m}}(z_{14})}{\overline{w}_{\mathrm{m},t}}(g_{14,t}^{\mathrm{m}} + g_{41,t}^{\mathrm{m}})\Big] \\
M_3(\mathbf{g}_t^{\mathrm{m}}) =& \frac{w_{\mathrm{m}}(z_{33})}{\overline{w}_{\mathrm{m},t}} g_{33,t}^{\mathrm{m}} + \frac{1}{2}\Big[ \frac{w_{\mathrm{m}}(z_{13})}{\overline{w}_{\mathrm{m},t}}(g_{13,t}^{\mathrm{m}} + g_{31,t}^{\mathrm{m}}) + \frac{w_{\mathrm{m}}(z_{34})}{\overline{w}_{\mathrm{m},t}}(g_{34,t}^{\mathrm{m}} + g_{43,t}^{\mathrm{m}}) + (1-r)\frac{w_{\mathrm{m}}(z_{23})}{\overline{w}_{\mathrm{m},t}}(g_{23,t}^{\mathrm{m}} + g_{32,t}^{\mathrm{m}}) + \\
& r\frac{w_{\mathrm{m}}(z_{14})}{\overline{w}_{\mathrm{m},t}}(g_{14,t}^{\mathrm{m}} + g_{41,t}^{\mathrm{m}})\Big] \\
M_4(\mathbf{g}_t^{\mathrm{m}}) =& \frac{w_{\mathrm{m}}(z_{44})}{\overline{w}_{\mathrm{m},t}} g_{44,t}^{\mathrm{m}} + \frac{1}{2}\Big[ \frac{w_{\mathrm{m}}(z_{24})}{\overline{w}_{\mathrm{m},t}}(g_{24,t}^{\mathrm{m}} + g_{42,t}^{\mathrm{m}}) + \frac{w_{\mathrm{m}}(z_{34})}{\overline{w}_{\mathrm{m},t}}(g_{34,t}^{\mathrm{m}} + g_{43,t}^{\mathrm{m}}) + (1-r)\frac{w_{\mathrm{m}}(z_{14})}{\overline{w}_{\mathrm{m},t}}(g_{14,t}^{\mathrm{m}} + g_{41,t}^{\mathrm{m}}) + \\
& r\frac{w_{\mathrm{m}}(z_{23})}{\overline{w}_{\mathrm{m},t}}(g_{23,t}^{\mathrm{m}} + g_{32,t}^{\mathrm{m}})\Big],
\end{aligned}
\tag{B-6}
$$

and

$$
\begin{aligned}
F_1(\mathbf{g}_t^{\mathrm{f}}) =& \frac{w_{\mathrm{f}}(z_{11})}{\overline{w}_{\mathrm{f},t}} g_{11,t}^{\mathrm{f}} + \frac{1}{2}\Big[ \frac{w_{\mathrm{f}}(z_{12})}{\overline{w}_{\mathrm{f},t}}(g_{12,t}^{\mathrm{f}} + g_{21,t}^{\mathrm{f}}) + \frac{w_{\mathrm{f}}(z_{13})}{\overline{w}_{\mathrm{f},t}}(g_{13,t}^{\mathrm{f}} + g_{31,t}^{\mathrm{f}}) + (1-r)\frac{w_{\mathrm{f}}(z_{14})}{\overline{w}_{\mathrm{f},t}}(g_{14,t}^{\mathrm{f}} + g_{41,t}^{\mathrm{f}}) + \\
& r\frac{w_{\mathrm{f}}(z_{23})}{\overline{w}_{\mathrm{f},t}}(g_{23,t}^{\mathrm{f}} + g_{32,t}^{\mathrm{f}})\Big] \\
F_2(\mathbf{g}_t^{\mathrm{f}}) =& \frac{w_{\mathrm{f}}(z_{22})}{\overline{w}_{\mathrm{f},t}} g_{22,t}^{\mathrm{f}} + \frac{1}{2}\Big[ \frac{w_{\mathrm{f}}(z_{12})}{\overline{w}_{\mathrm{f},t}}(g_{12,t}^{\mathrm{f}} + g_{21,t}^{\mathrm{f}}) + \frac{w_{\mathrm{f}}(z_{24})}{\overline{w}_{\mathrm{f},t}}(g_{24,t}^{\mathrm{f}} + g_{42,t}^{\mathrm{f}}) + (1-r)\frac{w_{\mathrm{f}}(z_{23})}{\overline{w}_{\mathrm{f},t}}(g_{23,t}^{\mathrm{f}} + g_{32,t}^{\mathrm{f}}) + \\
& r\frac{w_{\mathrm{f}}(z_{14})}{\overline{w}_{\mathrm{f},t}}(g_{14,t}^{\mathrm{f}} + g_{41,t}^{\mathrm{f}})\Big] \\
F_3(\mathbf{g}_t^{\mathrm{f}}) =& \frac{w_{\mathrm{f}}(z_{33})}{\overline{w}_{\mathrm{f},t}} g_{33,t}^{\mathrm{f}} + \frac{1}{2}\Big[ \frac{w_{\mathrm{f}}(z_{13})}{\overline{w}_{\mathrm{f},t}}(g_{13,t}^{\mathrm{f}} + g_{31,t}^{\mathrm{f}}) + \frac{w_{\mathrm{f}}(z_{34})}{\overline{w}_{\mathrm{f},t}}(g_{34,t}^{\mathrm{f}} + g_{43,t}^{\mathrm{f}}) + (1-r)\frac{w_{\mathrm{f}}(z_{23})}{\overline{w}_{\mathrm{f},t}}(g_{23,t}^{\mathrm{f}} + g_{32,t}^{\mathrm{f}}) + \\
& r\frac{w_{\mathrm{f}}(z_{14})}{\overline{w}_{\mathrm{f},t}}(g_{14,t}^{\mathrm{f}} + g_{41,t}^{\mathrm{f}})\Big] \\
F_4(\mathbf{g}_t^{\mathrm{f}}) =& \frac{w_{\mathrm{f}}(z_{44})}{\overline{w}_{\mathrm{f},t}} g_{44,t}^{\mathrm{f}} + \frac{1}{2}\Big[ \frac{w_{\mathrm{f}}(z_{24})}{\overline{w}_{\mathrm{f},t}}(g_{24,t}^{\mathrm{f}} + g_{42,t}^{\mathrm{f}}) + \frac{w_{\mathrm{f}}(z_{34})}{\overline{w}_{\mathrm{f},t}}(g_{34,t}^{\mathrm{f}} + g_{43,t}^{\mathrm{f}}) + (1-r)\frac{w_{\mathrm{f}}(z_{14})}{\overline{w}_{\mathrm{f},t}}(g_{14,t}^{\mathrm{f}} + g_{41,t}^{\mathrm{f}}) + \\
& r\frac{w_{\mathrm{f}}(z_{23})}{\overline{w}_{\mathrm{f},t}}(g_{23,t}^{\mathrm{f}} + g_{32,t}^{\mathrm{f}})\Big],
\end{aligned}
\tag{B-7}
$$

where $r$ is the recombination rate between the two loci, $w_u(z_{ij})/\overline{w}_u$ is the relative fecundity of a diploid adult with genotype $ij$ or $ji$ in sex $u$, and $\overline{w}_{u,t} = \sum_{i=1}^4 \sum_{j=1}^4 w_u(z_{ij})(g_{ij,t}^u + g_{ji,t}^u)$ is the sex-specific mean fecundity in the population at generation $t$.

##### B.2.2 Analysis

We focus here on a pair of unlinked loci ($r = 1/2$; linkage is considered in Appendix C.2). With $L = 2$, a necessary condition for balancing selection to occur at either locus is that all four gamete genotypes can be invaded by at least one other genotype when common, that is, when the equilibrium frequencies $m_{1,t} = f_{1,t} = 1, m_{2,t} = f_{2,t} = 1, m_{3,t} = f_{3,t} = 1$ and $m_{4,t} = f_{4,t} = 1$ are all unstable. For compactness, let us collect haploid genotype frequencies into a vector $\mathbf{n}_t = (m_{1,t}, m_{2,t}, m_{3,t}, m_{4,t}, f_{1,t}, f_{2,t}, f_{3,t}, f_{4,t})$. Following this lexicographic order, can write the four fixation states as $\hat{\mathbf{n}}_1 = (1,0,0,0,1,0,0,0)$, $\hat{\mathbf{n}}_2 = (0,1,0,0,0,1,0,0)$, $\hat{\mathbf{n}}_3 = (0,0,1,0,,0,1,0)$, $\hat{\mathbf{n}}_4 = (0,0,0,1,0,0,0,1)$. A given fixation state $\hat{\mathbf{n}}_i$ is unstable when the leading eigenvalue, $\lambda(\hat{\mathbf{n}}_i)$, of the Jacobian matrix

$$\mathrm{J}(\hat{\mathbf{n}}_i) = \begin{pmatrix} \frac{\partial F_1(\mathbf{g}_t^{\mathrm{f}})}{\partial f_{1,t}} & \cdots & \frac{\partial F_1(\mathbf{g}_t^{\mathrm{f}})}{\partial m_{4,t}} \\ \vdots & \vdots & \vdots \\ \frac{\partial M_4(\mathbf{g}_t^{\mathrm{m}})}{\partial f_{1,t}} & \cdots & \frac{\partial M_4(\mathbf{g}_t^{\mathrm{m}})}{\partial m_{4,t}} \end{pmatrix}_{\mathbf{n}_t=\hat{\mathbf{n}}_i} \tag{B-8}$$

is greater than unity ($\lambda(\hat{\mathbf{n}}_i) > 1$). Therefore all genotypes can be invaded when $\lambda(\mathbf{n}_i) > 1$ for all $i \in \{1,2,3,4\}$, which is the condition for selection to be balancing at a minimum of one locus. Plugging eq. (I.A) into eqs. (B-6)-(B-7), eqs. (B-6)-(B-7) into eqs. (B-4)-(B-5), and then eqs. (B-4)-(B-5) into eq. (B-8) and calculating the relevant eigenvalues we can derive the invasion conditions for all four genotypes. The resulting expressions are typically unsightly and thus not easily interpretable and so are plotted for different numerical values in Supplementary Fig. 1 (bottom row, we interpret these results in the next section, Appendix B.2.3).

Invadability of all genotypes is a sufficient condition for balancing selection to happen at least at one of the two loci. But it is not sufficient to know whether polymorphism will be maintained at both loci simultaneously. To investigate whether balancing selection occurred at one or two loci, we iterated numerically the recursions eqs. (B-4)-(B-5) and recorded how many loci were still polymorphic after $10^5$ generations (which was sufficient time for equilibrium polymorphic frequencies to be reached). We did this for a wide range of parameter values within $c_{\mathrm{m}} \in [0,1]$, $b \in [0,2]$ and $\alpha \in [0,1]$ where $c_{\mathrm{f}} = \alpha c_{\mathrm{m}}$, and with initial starting genotype frequencies of $m_1 = f_1 = 0.2499, m_2 = f_2 = 0.2499, m_3 = f_3 = 0.2501, m_4 = f_4 = 0.2501$. If the minor allele frequency at a locus exceeded $10^{-3}$, then this locus was considered to be polymorphic and under balancing selection.

###### B.2.3 Results

Analysis of the two-locus model reveals that polymorphism may be maintained at both loci, at only one locus, or at neither, depending on the shape of fecundity curves ($b$) and the strength of fecundity costs in each sex ($c_\mathrm{m}, c_\mathrm{f}$).

In line with the results from our continuum-of-alleles approach, polymorphism is maintained across both loci (Supplementary Fig. 1 bottom row panels, black regions) under the conditions that fecundity curves are accelerating ($b > 1$) and decline both strongly and with similar intensity in males and females ($c_\mathrm{m} \sim c_\mathrm{f} \gg 0$).

Meanwhile, balancing selection at just one locus requires that fecundity curves are diminishing ($b < 1$) and decline at moderately different rates between the sexes ($c_\mathrm{m} \neq c_\mathrm{f}$, Supplementary Fig. 1 bottom row panels, gray regions). Here, the allele favouring the more strongly selected sex becomes fixed at one locus ($\mathrm{A}_1$ or $\mathrm{A}_2$ when $c_\mathrm{f} > c_\mathrm{m}$, and $\mathrm{a}_1$ or $\mathrm{a}_2$ when $c_\mathrm{m} > c_\mathrm{f}$) and both alleles persist at the other. In this case, that is, given fixation at one locus, we can use eqs. (B-6)-(B-7) to find the equilibrium allele frequency at the remaining polymorphic locus (i.e., by finding equilibrium frequencies of the two segregating haploid genotypes), as well as the level of heterozygosity this creates. The equilibrium frequency, $p_k^*$, of the female-beneficial allele $\mathrm{A}_k$ at locus $k$ when $c_\mathrm{m} > c_\mathrm{f}$ is

$$p_k^* = \frac{(2^b - 3^b)c_\mathrm{f} + (2^b - 1)c_\mathrm{m}}{(2^b - 2\cdot 3^b) + 4^b)c_\mathrm{f} + (2^b - 2)c_\mathrm{m}} + \mathcal{O}(\epsilon^2) \tag{B-9}$$

(i.e., the frequency of $\mathrm{A}_1$ in a population fixed for $\mathrm{a}_2$, or the frequency of $\mathrm{A}_2$ in a population fixed for $\mathrm{a}_1$, where $c_\mathrm{m} \sim \mathcal{O}(\epsilon), c_\mathrm{f} \sim \mathcal{O}(\epsilon)$, Appendix Fig. 1A), and when $c_\mathrm{m} < c_\mathrm{f}$, the equilibrium frequency, $q_k^* = 1 - p_k^*$, of the male-beneficial allele $\mathrm{a}_k$ at locus $k$ is

$$q_k^* = \frac{(2^b - 1)c_\mathrm{f} + (2^b - 3^b)c_\mathrm{m}}{(2^b - 2)c_\mathrm{f} + (2^b - 2\cdot 3^b + 4^b)c_\mathrm{m}} + \mathcal{O}(\epsilon^2) \tag{B-10}$$

(i.e., the frequency of $\mathrm{a}_1$ when the population is fixed for $\mathrm{A}_2$, and $\mathrm{a}_2$ when the population is fixed for $\mathrm{A}_1$). From Appendix Fig. 1A we see that, at the polymorphic locus $k \in \{1, 2\}$, the frequency of the female-beneficial allele $\mathrm{A}_k$ decreases (and accordingly, the frequency of the male-beneficial allele $\mathrm{a}_k$ a increases) sharply with the strength of fecundity costs in males relative to females ($c_\mathrm{m}/c_\mathrm{f}$). Consequently, there is relatively little parameter space where $p_k^*$ and $q_k^*$ have intermediate frequencies (i.e. close to 0.5), and so average heterozygosity across both loci is typically low even when selection is balancing at one locus (Appendix Fig. 1B).

Finally, for the remaining combinations of parameters $b, c_{\mathrm{m}}$ and $c_{\mathrm{f}}$, selection disfavours polymorphism at both loci (Supplementary Fig. 1 bottom row panels, white regions) driving the fixation of one of the four homozygote genotypes ($A_1A_1a_2a_2$, $a_1a_1A_2A_2$, $a_1a_1a_2a_2$, or $A_1A_1A_2A_2$).

In sum, when $L = 2$, the parameter space for sexually antagonistic selection to drive polymorphism at at least one locus is relatively permissive and similar to that when $L = 1$. However, the conditions for polymorphism at two loci are restrictive, requiring strong selection. Thus, sexual antagonism has only a limited capacity to increase heterozygosity across both loci.

##### B.3 Dynamics at many loci

To characterise evolutionary dynamics for a polygenic trait, we ran individual-based simulations of our di-allelic model with many loci. All simulations were conducted using SLiM 4 [7].

Following the lifeycle described in section 2, we consider a diploid population of fixed size comprising 5,000 adult males and 5,000 adult females with non-overlapping generations. At the beginning of the offspring generation, say $t+1$, 5,000 male and 5,000 female offspring are created, with the mother of each offspring being sampled from amongst the female adults of the previous generation $t$, and the fathers being sampled from amongst the male adults. Sampling performed is with replacement from a multinomial distribution where the weighting on each individual is given by their fecundity, which, following eq. (I.A), is

$$\begin{aligned} w_{\mathrm{m}}(z) &= (1-c_{\mathrm{m}}) + c_{\mathrm{m}}\left(\frac{\theta - z}{2\theta}\right)^{b_{\mathrm{m}}} \\ w_{\mathrm{f}}(z) &= (1-c_{\mathrm{f}}) + c_{\mathrm{f}}\left(\frac{\theta + z}{2\theta}\right)^{b_{\mathrm{f}}}, \end{aligned} \tag{B-11}$$

in males and females, respectively. After sampling, adults are removed from the simulations, juveniles become adults and the new generation begins. Recombination and mutation occurs for each juvenile as outlined in section 2.

Simulations were initiated with equal allele frequencies at all loci ($p_k = 0.5$ for all $k$, where $p_k$ is frequency of allele A at locus $k$) and with genotypes at Hardy-Weinberg proportions, and run for $10^6$ generations. In a given generation, heterozygosity at locus $k$ was calculated as

$$H_k = 1 - (1-p_k)^2 - p_k^2 \tag{B-12}$$

and average heterozygosity across loci was calculated as

$$H = \sum_{k=1}^{L} \frac{H_k}{L}. \tag{B-13}$$

To produce Fig. 3B, calculations were performed at at 100 generation intervals and then averaged over the last 900,000 generations of a simulation. The expected heterozgosity for a neutral locus was given by $H = 4N\mu/[1+8N\mu] = 1/7$ (eq. 2.24 in [14] with their $k = 2$).

### C Extensions

#### C.1 Loci with different effect sizes

Here, we investigate the case where loci have different phenotypic effects in our di-allelic model with power functions (eq. (I.A)), i.e., we allow variation in $\delta_k$ among the $L$ loci encoding $z$.

##### C.1.1 Two locus model

For the case of $L = 2$, we assume that locus 1 has greater effect than locus 2 and set $\delta_2 = \rho\delta_1$ with $0 < \rho < 1$ (and where allelic effects at loci 1 and 2 are given by eq. 3). Using the same notation as in Appendix B.2 (i.e., $z_{ij}$ is the phenotype encoded by the diploid genotype $ij$ where $i \in \{1,2,3,4\}$ and $j \in \{1,2,3,4\}$ are the paternally and paternally inherited haploid genotypes, respectively), we have the following phenotypes

$$\begin{aligned}
z_{11} &= 2\delta(1+\rho)\\
z_{12} &= 2\delta\\
z_{13} &= 2\delta\rho\\
z_{14} &= 0\\
z_{22} &= 2\delta(1-\rho)\\
z_{24} &= -2\delta\rho\\
z_{33} &= 2\delta(\rho-1)\\
z_{34} &= -2\delta\\
z_{44} &= -2\delta(1+\rho),
\end{aligned} \tag{C-1}$$

with female and male optima of $\theta = 2\delta(1+\rho)$ and $-\theta = -2\delta(1+\rho)$, respectively. After plugging eq. (C-1) into eqs. (B-4)-(B-7) we analyse this model following the same procedure as in Appendix B.2.

We find that the main consequence of varying the ratio of the locus effect sizes ($\rho$) is that it adjusts the parameter space for single locus polymorphisms when fecundity curves are diminishing ($b < 1$, Appendix Fig. 2, top row panels). This is because variation in the phenotypic contributions across loci modulates the phenotypes encoded by intermediate double homozygote genotypes (i.e. the phenotypes encoded by genotypes $A_1A_1a_2a_2$ and $a_1a_1A_2A_2$ diverge from each other and from $z = 0$ when $\rho < 1$). This changes the specific parameters for which double homozygotes will produce high fitness phenotypes, and thus the conditions that will favour a polymorphic remainder locus. By contrast, $\rho$ has relatively little effect on the conditions for maintaining two polymorphic loci, which still requires accelerating fecundity curves and strong selection ($b > 1$, $c_m$ and $c_f$ large and nearly equal, although note the parameter space for two-locus polymorphisms is slightly more permissive when $\rho$ is small, Appendix Fig. 2, bottom row panels).

##### C.1.2 Polygenic model

To run simulations of our polygenic model ($L \gg 1$) when alleles have different effect sizes, we used the same procedure as for equal effects (Appendix B.3), but with the effect sizes of individual loci now being drawn from an exponential distribution with mean 1, i.e., $\delta_k \sim \text{Exp}(1)$. The female optimum, which is given by the maximum phenotype, then is $\theta = \sum_1^L 2\delta_k$, and the male optimum, which is given by the minimum phenotype, is $-\theta = \sum_1^L 2\delta_k$.

We find that heterozygosity patterns are similar to the case with equal effect sizes in the sense that average heterozygosity is only greater than neutral when selection is diversifying (i.e., when $b > 1$ and $c_m$ and $c_f$ being similar and large; Appendix Fig. 3 left-most panels, compare top and bottom). In addition, we find that variation at individual loci that persists when selection is stabilising ($b < 1$) due to the lack of homozygous allelic combinations near $z^*$ exhibits the same transient behaviour as in the equal effects case (Appendix Fig. 4). This occurs because multiple allele combinations are capable of producing polygenic trait values similarly close to $z^*$ (even if locus effect sizes vary), so that drift and mutation to drive turnover in the identity of the polymorphic locus. Together with the results of section C.1.1, this indicates that variation in effect sizes across loci weakly influences the conditions under which elevated heterozygosity is expected, and only does so when the number of loci is especially small (e.g., $L = 2$) and variation especially large ($\rho$ small).

##### C.2 Effect of recombination between loci

Here we relax our previous assumption of free recombination and explore our two-locus and polygenic di-allelic model with a lower recombination rate between loci ($r \leq 0.5$).

###### C.2.1 Two locus model

We analysed the two-locus model with linkage using the recursions eq. (B-6)-(B-7), considering both equal and different phenotypic effect sizes (plugging eq. C-1 into eqs. B-4-B-7 with $\rho = 1$ and $\rho < 1$, respectively).

In general, reduced recombination has a limited effect on the maintenance of genetic variation (Appendix Fig. 5). Specifically, when loci had equal phenotypic effects ($\delta_1 = \delta_2 = \delta$), variation maintained by sexual antagonism was generally insensitive to recombination rate, with the exception that limited recombination moderately reduces the strength of the fecundity trade-off required for two-locus polymorphisms to arise under accelerating fecundity curves ($b > 1$, Appendix Fig. 5A). An additional effect of recombination arises when loci have different effect sizes, whereby tight linkage can generate two-locus polymorphisms when fecundity curves are diminishing ($b < 1$ and, e.g., $r \sim \mathcal{O}(0.001)$, Appendix Fig. 5B). Here, differences in allelic effect sizes mean that no homozygote genotype encodes a high fecundity intermediate phenotype, and the fittest genotypes are instead double heterozygotes. When recombination is weak, this leads to increased genetic variation because double heterozygotes produce almost exclusively $A_1a_2$ and $a_1A_2$ gametes that go on to form high fitness offspring, allowing selection to maintain both gametic genotypes and so two polymorphic loci. Such elevated genetic variation owing to tight linkage has been found in previous studies of sex-concordant stabilising selection [15, 16], although this effect breaks down when the number of trait loci increases ($L > 2$, [16]).

###### C.2.2 Polygenic Model

When $L$ is polygenic, limited recombination also has a weak effect on genetic variation. The conditions for elevated heterozygosity in the presence of genetic linkage are only slightly more permissive than those with free recombination (still requiring $b > 1$, $c_m$ and $c_f$ similar and large), even when loci have different effect sizes and recombination is extremely weak (Appendix Fig. 3). Notably, in contrast to the $L = 2$ case, elevated heterozygosity is not observed when $b < 1$ for any recombination rate considered, consistent with the notion that the effects of linkage on genetic variation under stabilising selection in two-locus models do not generalise to arbitrary numbers of loci [16]. This is likely because the large variety of allele combinations possible when $L$ is large means a homozygote genotype is typically available that encodes a fit enough phenotype to fix, and so linkage does not generate conditions for multi-locus polymorphisms. Weak recombination, however, drives higher levels of heterozygosity for the parameter values that lead to diversifying selection (Appendix Fig. 3). This is in line with theory on quantitative traits mediating intraspecific competition, where, under strong diversifying selection, population genetic variance, rather than the space maintaining polymorphism, is sensitive to recombination rate [17].

#### C.3 Differently shaped fecundity landscapes in males and females

Here, we repeat our analyses of the continuum-of-alleles and di-allelic models with power functions, now allowing for fecundity curves to have different shapes in males and females. Specifically, we now allow for different curvatures in the two sexes, $b_{\mathrm{m}} \neq b_{\mathrm{f}}$ in Eq. (I.A).

##### C.3.1 Continuum-of-alleles model

When fecundity curves have sex-specific shapes, evolutionary dynamics are complicated and so we proceeded with a numerical analysis. That is, we first plugged eq. (A-5) into eq. (A-6) and solved eq. (A-7) numerically with eq. (I.A) to find singular strategies $x^*$, whose stability was then determined. In particular, we focused on identifying the conditions under which sexual antagonism leads to a singularity exhibiting diversifying selection (i.e., an evolutionary branching point), as the existence of such a point is necessary for selection to favour polymorphism.

Overall, we find that the criteria for diversifying selection when $b_{\mathrm{m}} \neq b_{\mathrm{f}}$ are in line with our results for $b_{\mathrm{m}} = b_{\mathrm{f}}$, with such selection only arising under restricted conditions (Appendix Fig. 6 top row, green region). First, selection must be strong. Second, diversifying selection requires at least one fecundity curve be accelerating ($b_{\mathrm{m}} > 1$ and/or $b_{\mathrm{f}} > 1$), with conditions being most permissive when one curve is accelerating and the other is at least linear (i.e., $b \geq 1$ in the sex with the least accelerating curve, Appendix Fig. 6 top row). Diversifying selection is possible if one fecundity curve is diminishing, but requires the curve in the other sex to be especially accelerating (e.g., if $b_{\mathrm{f}} < 1$, $b_{\mathrm{m}}$ needs to well exceed 1, Appendix Fig. 6 top row left-hand panel). Even stronger sexually antagonistic selection is required when one of the curves is diminishing than when both are accelerating (Appendix Fig. 6).

##### C.3.2 Polygenic model

Following Appendix B.3, we ran simulations of our di-allelic model assuming $z$ has a polygenic basis ($L = 10$). As in the case of the sexes showing the same fecundity curves, we find that conditions for which we observe elevated heterozygosity closely resembles those for diversifying selection in the continuum-of-alleles model (Appendix Fig. 6, compare top and bottom rows), indicating that such selection is required for the maintenance of genetic polymorphism across loci. Furthermore, in line with our previouus results, we also find that heterozygosity is greatest where both curves are accelerating and selection is very strong ($b_\mathrm{m} > 1$, $b_\mathrm{f} > 1$, $c_\mathrm{m} = c_\mathrm{f} \sim 1$, Appendix Fig. 6, bottom righ-hand panel), as this generates the strongest diversifying selection.

#### C.4 Gaussian fecundity

Here we repeat our previous analysis of power fecundity functions but now assuming fecundity follows the Gaussian functions from continuous trait models of sex-specific selection (e.g., [11, 18–21]). Specifically,

$$\begin{aligned} w_\mathrm{m}(z) &= K_\mathrm{m} e^{-\sigma_\mathrm{m}(z+\theta)^2} \\ w_\mathrm{f}(z) &= K_\mathrm{f} e^{-\sigma_\mathrm{f}(z-\theta)^2}, \end{aligned} \tag{C-2}$$

where $\sigma_\mathrm{m} > 0$ and $\sigma_\mathrm{f} > 0$ scale the width of the fecundity functions around $\pm\theta$ and thus the intensity of the fitness decline as $z$ departs from each sex's optimum. These functions allow for a continuous range of phenotypes (so that we need not constrain $-2\theta \leq z \leq 2\theta$), but with the disadvantage the shape of the fecundity curves (e.g., $b_\mathrm{m}$ and $b_\mathrm{f}$ in eq. I.A) and the strength of the fecundity trade-off across the sexes (e.g., $c_\mathrm{m}$ and $c_\mathrm{f}$ in eq. I.A) cannot be tuned independently.

##### C.4.1 Continuum-of-alleles

Following eqs. A-5-A-9 we find that under the continuum-of-alleles model the selection gradient is given by

$$S(x) = \theta(\sigma_\mathrm{f} - \sigma_\mathrm{m}) - 2x(\sigma_\mathrm{f} + \sigma_\mathrm{m}) \tag{C-3}$$

with a single singularity at

$$x^* = \theta \cdot \frac{\sigma_\mathrm{f} - \sigma_\mathrm{m}}{2(\sigma_\mathrm{f} + \sigma_\mathrm{m})}. \tag{C-4}$$

This point is convergence stable for all values of $\sigma_\mathrm{m}, \sigma_\mathrm{f}$ because

$$S'(x^*) = -2(\sigma_\mathrm{f} + \sigma_\mathrm{m}) < 0 \tag{C-5}$$

is always negative, and thus selection is always either stabilising or diversifying depending on the sign of

$$H(x^*) = \frac{\sigma_\mathrm{f}^3 + 3\sigma_\mathrm{f}\sigma_\mathrm{m}^2 + \sigma_\mathrm{m}^3 + \sigma_\mathrm{f}^2\sigma_\mathrm{m}(16\theta^2\sigma_\mathrm{m} - 3)}{(\sigma_\mathrm{f} + \sigma_\mathrm{m})^2}, \tag{C-6}$$

which is positive whenever

$$4\theta > \sqrt{\frac{(\sigma_\mathrm{m} + \sigma_\mathrm{f})^3}{\sigma_\mathrm{m}^2\sigma_\mathrm{f}^2}}. \tag{C-7}$$

(see Appendix Fig. 7A for plot).

##### C.4.2 Di-allelic simulations

Following the same procedure as in Appendix A.4.3, we ran mutlilocus simulations of the di-allelic model where the weighting on each male and female during multinomial sampling is given by

$$\begin{aligned} w_\mathrm{m}(z) &= e^{-\sigma_\mathrm{m}(z+\theta)^2} \\ w_\mathrm{f}(z) &= e^{-\sigma_\mathrm{f}(z-\theta)^2}. \end{aligned} \tag{C-8}$$

We ran simulations for values of $\sigma_\mathrm{m} \in [0, X]$ and $\sigma_\mathrm{f} \in [0, X]$ with $\theta = 20$.

As with our results for power functions, we found the parameter combinations that produced elevated heterozygosity in the multilocus model well matched those inducing diversifying selection in the continuum-of-alleles model (Appendix Fig. 7A-B).

#### C.5 Polymorphism with large effect alleles

##### C.5.1 One locus

Our analyses of the continuum-of-allele model have previously considered the emergence of polymorhpism from a monomorphic population through mutations of small effect. In this section we investigate the potential for sexual antagonism to maintain a pre-existing genetic polymorphism, or one that arises from a large effect mutation. Specifically, we examine the conditions for a polymorphic equilibrium at a locus with two alleles, say $x_1$ and $x_2$, to be stable to the invasion of a third allele (i.e., is an uninvadable coalition). In general, this requires that we track the frequencies of the two resident alleles in male and female gametes separately (as in eq. A-1) and conduct a full stability analysis of the recursions. When selection is weak (in the sense that variation in $w_{\mathrm{m}}(z)$ and $w_{\mathrm{f}}(z)$ is small), however, the allele frequencies are approximately equal in the gamete pools of the two sexes. This considerably simplifies analysis, and we use this assumption throughout this section.

We focus our attention on functions that are symmetric across the sexes (about $z = 0$), which should be the most conducive to polymorphism. Specifically, we set

$$\begin{aligned} w_{\mathrm{m}}(z) &= \phi(-z) \\ w_{\mathrm{f}}(z) &= \phi(z) \end{aligned} \tag{C-9}$$

where $\phi : \mathbb{R} \to \mathbb{R}_{\geq 0}$ is the fecundity function in females. Before considering a polymorphism, we note that a singular trait valued $x^*$ is such that

$$\phi'(2x^*) = \phi'(-2x^*), \tag{C-10}$$

according to eq. (A-11) with eq. (C-9) under weak selection. Hence

$$x^* = 0, \tag{C-11}$$

is always a singular strategy, as expected given that the symmetry of the male and female fecundity curves about $z = 0$. Convergence stability and uninvadability of $x^*$, meanwhile, are determined by

$$\begin{aligned} S'(x^*) &= \phi''(2x^*) + \phi''(-2x^*) \\ H(x^*) &= \frac{1}{2}\left(\phi''(2x^*) + \phi''(-2x^*)\right), \end{aligned} \tag{C-12}$$

respectively.

Given the symmetry of the fecundity functions (eq. C-9), we expect that two alleles $x_1$ and $x_2$ must be equally distant from zero to be part of a stable coalition. So we can set

$$
\begin{aligned}
x_1 &= -x \\
x_2 &= x,
\end{aligned}
\tag{C-13}
$$

where $x > 0$. Reciprocal invasion of $x_1$ and $x_2$ requires that $W(x_1, x_2) > 1$ and $W(x_2, x_1) > 1$, which using from eqs. (A-4) and (C-13) is equivalent to,

$$
\phi(0) > \frac{\phi(-2x) + \phi(2x)}{2}. \tag{C-14}
$$

This can be seen as the requirement of dominance reversal as it says that the heterozygote (expressing 0) must have greater fecundity than the average of the two homozygotes (expressing $2x$ and $-2x$) in both sexes.

To determine whether a polymorphism consisting of $x_1$ and $x_2$ is an uninvadable coalition under the continuum-of-alleles model, we need to first characterize the invasion fitness $W(x_\bullet, (x_1, x_2))$ of a mutant $x_\bullet$ in a resident population that is polymorphic and at equilibrium (i.e., where the frequencies of $x_1$ and $x_2$ have converged to their equilibrium). Because selection is weak and the model shows perfect symmetry among males and females, the equilibrium frequency of each resident allele $x_1 = -x$ and $x_2 = x$ is simply $1/2$ in both males and females. It is straightforward to show that in this case, the invasion fitness of a third allele for the symmetric model (eq. C-9) is given by

$$
W(x_\bullet, (-x, x)) = 1 + \frac{1}{4}\Big(\phi(x_\bullet + x) + \phi(x_\bullet - x) + \phi(-x_\bullet - x) + \phi(-x_\bullet + x) - 2\phi(0) - \phi(2x) - \phi(-2x)\Big) + \mathcal{O}(s_\phi^2) \tag{C-15}
$$

where $s_\phi$ is the largest absolute difference in fecundity among two males or two females. Directional selection on each allelic value is given by

$$
\begin{aligned}
S_1(-x, x) &= \left.\frac{\partial W(x_\bullet, (-x, x))}{\partial x_\bullet}\right|_{x_\bullet = -x} \\
S_2(-x, x) &= \left.\frac{\partial W(x_\bullet, (-x, x))}{\partial x_\bullet}\right|_{x_\bullet = x},
\end{aligned}
\tag{C-16}
$$

respectively. A singular coalition $(-x^*, x^*)$ is defined such that

$$
S_1(-x^*, x^*) = S_2(-x^*, x^*) = 0. \tag{C-17}
$$

Substituting eq. (C-15) into eq. (C-16), we find that eq. (C-17) requires that

$$\phi'(2x^*) = \phi'(-2x^*), \tag{C-18}$$

i.e., that rises in fecundity in one sex are exactly offset by declines in the other near both resident homozygotes. Such a singular coalition will be approached gradually from nearby coalitions through evolution (i.e., be convergence stable), when the leading real part of the eigenvalues of the Jacobian matrix

$$\begin{pmatrix} \frac{\partial S_1(x_1,x_2)}{\partial x_1} & \frac{\partial S_1(x_1,x_2)}{\partial x_2} \\ \frac{\partial S_2(x_1,x_2)}{\partial x_1} & \frac{\partial S_2(x_1,x_2)}{\partial x_2} \end{pmatrix}_{\substack{x_1=-x^* \\ x_2=x^*}}, \tag{C-19}$$

is negative. We find that this is equivalent to

$$\begin{aligned} &\phi''(-2x^*) + \phi''(2x^*) < 0 \\ &2\phi''(0) + \phi''(-2x^*) + \phi''(2x^*) < 0. \end{aligned} \tag{C-20}$$

Note that it is possible for a polymorphic coalition to be convergence stable under weak selection, even though we do not expect evolutionary branching from the singular state at $x^* = 0$. This requires, however, that the fecundity curves be sufficiently negatively curved to satisfy eq. (C-20) and yet also have equal slopes on the fitness surface at both homozygous phenotypes so that eq. (C-18) holds (see Appendix Fig. 8 for an example).

Uninvadability of this coalition with respect to additional alleles requires that

$$\begin{aligned} &\left.\frac{\partial^2 W(x_\bullet,(-x,x))}{\partial x_\bullet^2}\right|_{x_\bullet=-x=-x^*} < 0 \\ &\left.\frac{\partial^2 W(x_\bullet,(-x,x))}{\partial x_\bullet^2}\right|_{x_\bullet=x=x^*} < 0. \end{aligned} \tag{C-21}$$

Plugging eq. (C-15) into eq. (C-21), we obtain that

$$2\phi''(0) + \phi''(-2x^*) + \phi''(2x^*) < 0 \tag{C-22}$$

must hold for the local uninvadability of a singular coalition $(-x^*, x^*)$. The fact that the same condition is necessary for convergence stability (eq. C-20) and for stabilizing selection (eq. C-22) indicates that there are no fecundity functions $\phi(z)$ such that another allele emerges from gradual evolution when the population is already dimorphic (i.e., there is no further branching under weak selection).

Comparing Eqs. (C-18), (C-20) and (C-22) with Eqs. (C-10) and (C-12) reveals that if a singular coalition $(-x^*, x^*)$ is convergence stable and thus uninvadable, then so are either $-x^*$ or $x^*$ in a monomorphic state (i.e., if Eqs. C-18, C-20 and C-22 hold, then so do Eqs. C-10 and C-12). This suggest that when a polymorphism that is locally stable exists, it is not very robust to perturbations as any monomorphic state where one of the two allele is lost (e.g., due to genetic drift) is also locally uninvadable.

Additionally, it is relatively straightforward for the strategy $x^* = 0$ (eq. C-11) to invade a stable coalition. Setting $x_\bullet = 0$ into eq. (C-15) and re-arranging, we obtain that this allele will invade a stable coalition when

$$\underbrace{\phi(0) - \frac{\phi(-2x^*) + \phi(2x^*)}{2}}_{>0 \text{ from eq. (C-14)}} - 2\left[\phi(0) - \frac{\phi(-x^*) + \phi(x^*)}{2}\right] > 0, \tag{C-23}$$

which essentially requires that $\phi(z)$ decelerates with $z$ between $-2x^*$ and $2x^*$ (assuming it is monotonic). This can be seen from the fact that

$$\phi(0) - \frac{\phi(-2x^*) + \phi(2x^*)}{2} - 2\left[\phi(0) - \frac{\phi(-x^*) + \phi(x^*)}{2}\right] = -\phi''(0){x^*}^2 + \mathcal{O}\left({x^*}^4\right). \tag{C-24}$$

Hence, $\phi''(0) < 0$ favours the invasion of an allele coding for zero in a dimorphic population. In addition, $\phi''(0) < 0$ entails that this allele is convergence stable and uninvadable when monomorphic (see eq. C-12).

##### C.5.2 Multiple loci

The above shows there exists fecundity functions $\phi(z)$ such that the population may switch from being monomorphic to polymorphic but this requires either a polymorphism to exist initially or large step mutations. Where a trait is polygenic and encoded by multiple loci of smaller effect, such large steps are unlikely as each locus contributes less variation – with such architecture, the appearance of highly divergent genotypes would entail, for example, secondary contact between two differentiated populations. Unless loci are in tight linkage, it is therefore more likely for the population to become monomorphic for a polygenic sexually antagonistic trait and remain so even when fecundity functions allow for the existence of stable coalitions.

We explored this with individual-based simulations. Specifically, we considered a version of our diallelic model where $z$ is encoded by $L = 10$ loci. We use fecundity functions that lead to stable dimorphism in the continuum-of-allele model (see Appendix Fig. 8). We initialise the population with two alleles $-x$ and $x$ present at each locus. We assume that there is initially full linkage disequilibrium, with all alleles of the same sign fully linked on the same homologous copy of a non-recombining chromosome (i.e., an individual carries the same allele at the first copy of every locus, and the same allele at the second copy of every locus). At the beginning of a simulation, chromosomes were constructed with all positive-effect alleles (carrying all $x$) or all negative-effect alleles (carrying all $-x$) with equal probability, so that simulations start with two segregating haplotypes at approximately equal frequency with total phenotypic contributions $10x$ and $-10x$ (as we assume $L = 10$). We initially assumed there was no recombination between loci and no new mutations ($r = 0$, $\mu = 0$). After 10,000 generations, we allowed recombination to occur ($r > 0$), so that alleles could become shuffled between chromosomes. We recorded average heterozygosity across loci through time (eq. B-13 where $p_k$ is the frequency of allele encoding $x$ at locus $k \in \{1, \ldots, 10\}$). The results of simulations with $r = 0.5$, $r = 0.05$, and $r = 0.005$ are shown in Appendix Fig. 9 (with fecundity functions as in Appendix Fig. 8 and $x = 0.059$ so that the two initial haplotypes, $10x = 0.59$ and $-10x = -0.59$, correspond to the stable allelic coalition at one locus for this scenario).

As expected, heterozygosity is initially high when linkage is complete (Appendix Fig. 9, $r = 0$), as both haplotypes are stably maintained at equal frequency by selection. However, once recombination occurs (Appendix Fig. 9, $r > 0$), heterozygosity rapidly decays as the shuffling of alleles reduces the divergence between haplotypes and the polymorphism becomes unstable. This continues until eventually a single haplotype encoding a monomorphic equilibrium fixes (eq. (C-10)). Thus, while it is possible under certain conditions for a sexually antagonistic polymorphism to be uninvadable by small-effect mutations and so maintained by selection at a single locus, this does not generalise to the maintenance of genetic variation in a polygenic trait.

##### C.6 Connections to other models of genetic conflict

Here, we connect the model of sexual antagonism that we study to a few other relevant models of genetic conflict that have been proposed as potential drivers of polymorphism. To make these connections and see more easily the similarities and differences among models, we use the invasion fitness $W(x_\bullet, x)$ of a rare genetic mutant coding for allelic value $x_\bullet$ in a population resident for $x$ (to be compared with eq. A-5). This allows us to focus on the fundamental drivers of polymorphism rather than on genetic constraints (such as number of loci, effect sizes, or non-additive genetic effect on phenotype).

**Local adaptation.** The closest connection our model has is with models of local adaptation where individuals inhabit one of two large patches. Here, the fitness of expressing phenotype $z$ is given by $w_1(z)$ in one patch and $w_2(z)$ in the other. Assuming additive gene action on the phenotype, that selection is soft (i.e. density regulation occurs before dispersal), and that generations do not overlap, invasion fitness of a rare allele coding for $x_\bullet$ in a diploid population monomorphic for $x$ is given by the leading eigenvalue $\lambda$ of the mean matrix, i.e. by

$$W(x_\bullet, x) = \lambda \begin{pmatrix} (1-m)\dfrac{w_1(x_\bullet + x)}{w_1(2x)} & m\dfrac{w_2(x_\bullet + x)}{w_2(2x)} \\ m\dfrac{w_1(x_\bullet + x)}{w_1(2x)} & (1-m)\dfrac{w_2(x_\bullet + x)}{w_2(2x)} \end{pmatrix} \tag{C-25}$$

where $m$ is the probability of dispersal [22, 23]. It is straightforward to show that when $m = 1/2$, $W(x_\bullet, x)$ reduces to eq. (A-5) with $w_1 = w_\mathrm{m}$ and $w_2 = w_\mathrm{f}$. This connection between the evolution of sexual antagonism and of local adaptation had also been highlighted using population genetics (i.e. change in allele frequency [24]). In addition, we also examine the link between local adaptation with limited dispersal ($m < 1/2$) and sex-specific selection at sex-linked loci in a later Appendix section (App. G.2).

**Sex-specific genotype-phenotype map.** Another relevant model we can connect to is one of sexual antagonism where it is assumed that selection is the same in males and females, but that the genotype-phenotype map differs among the sexes (although this map is additive [25]). To do this, we assume that when an allele encodes a phenotypic effect $x$ in females, it encodes an effect $z_\mathrm{m}(x)$ in males, i.e. $z_\mathrm{m}$ is the mapping from female to male allelic effects (letting $z_\mathrm{m}(x) = x$, this collapses to a model with sex-independent effects). Assuming a Wright-Fisher model of reproduction, the invasion fitness of a rare allele coding for $x_\bullet$ in a diploid population monomorphic for $x$ in this model can be written as,

$$W(x_\bullet, x) = \frac{1}{2}\frac{w(z_\mathrm{m}(x_\bullet) + z_\mathrm{m}(x))}{w(2z_\mathrm{m}(x))} + \frac{1}{2}\frac{w(x_\bullet + x)}{w(2x)}, \tag{C-26}$$

where $w(z)$ is the reproductive success of an individual of either sex with trait $z$. We can obtain further insights if we follow [25] and assume Gaussian selection $w(z) = \exp(-(s/2)z^2)$, which favours optimal trait value $z = 0$ with strength $s$, and that $z_\mathrm{m}(x) = \alpha_0 + \alpha_1 x$ is linear (this is what [25]'s approach of sampling allelic effects in males and females independently for two alleles at each locus boils down to if we focus on evolution at a single locus). Following the approach described in Appendix A.2, we find that selection is diversifying when

$$s > \frac{1}{\alpha_0^2} \frac{\left(1+\alpha_1^2\right)^3}{8\alpha_1^2} \tag{C-27}$$

holds, i.e. when selection is sufficiently strong ($s$ large), and male and female allelic values are sufficiently offset in a systematic manner (i.e. $\alpha_0$ large).

In performing a numerical analysis over "several thousands sets of parameters", [25] find no evidence of polygenic polymorphism, even when selection is strong ($s$ large). This suggests that selection across the sexes is always stabilising in their analyses. This is likely because, as indicated by eq. (C-27), diversifying selection requires the male and female genotype-phenotype maps be sufficiently offset such that the sexes produce highly differentiated phenotypes ($\alpha_0$ large). The conditions for this to happen in a polygenic model are not straightforward, entailing large systematic differences in sex-specific allelic values across loci: for example, where the effects of alleles on male phenotype at different loci all point in the same direction (e.g., all positive) and where this direction is different to that on female phenotype (e.g., all negative). Such systematic differences are especially unlikely under the assumptions made in [25], where alleles are constrained to have phenotypic effects of the same sign in both sexes.

Consequently, while both sex-specific fitness landscapes (eq. A-5) and sex-specific genotypic effects (eq. C-26) can produce diversifying selection at a single locus, polygenic variation is more likely in the former. Ultimately this difference reflects a distinction in the biology of sexual antagonism in these two cases: while male and female fitness landscapes may plausibly show strong divergence owing to sex-specific ecology, it is less obvious why male and female genotype-phenotype maps would differ so strongly and so systematically across many loci for a trait that is under sex-concordant stabilising selection.

**Disruptive but not frequency-dependent selection.** Another study we can link is [26], where a model of disruptive selection (but without negative-frequency dependence, i.e., not diversifying selection) is proposed. In this model, invasion fitness takes the general form

$$W(x_\bullet, x) = \frac{w_1(x_\bullet + x) + w_2(x_\bullet + x)}{w_1(2x) + w_2(2x)} \tag{C-28}$$

(eq. 1 in [26] where $w_1$ and $w_2$ are two Gaussian functions with different optima). Like eq. (C-25), eq. (C-28) is also compatible with a model of local adaption but where selection is hard, i.e. regulation occurs after dispersal (unlike eq. C-25 where recall selection is soft), and dispersal is random ($m = 1/2$). This entails that competition occurs equally among individuals of either patch type. The nature of competition here is therefore fundamentally different to the scenario that we explore, where competition occurs within each sex (as seen with the separate divisions by resident reproductive output for each sex in eq. A-5). Owing to competition within each sex, the case of sexual antagonism we explore allows for the emergence of polymorphism through evolutionary branching via diversifying selection. In contrast, a straightforward analysis following the approach described in Appendix A.2 with invasion fitness given by eq. (C-28) reveals that selection can never be diversifying here. We can therefore conclude that selection will never maintain variation within and between loci under eq. (C-28) in the absence of genetic constraints (i.e., that the trait is encoded by a single di-allelic locus). [26] arrives at the same conclusions using multi-loci population genetics recursions.

**Antagonistic pleiotropy and temporal heterogeneity.** Finally, we connect to models of antagonistic pleiotropy (e.g. [27]), which can be seen as a form of temporal heterogeneity within generations. A simple scenario envisaging this is where individuals experience two environments one after the other before reproduction. An individual expressing trait value $z$ survives the first environment with probability $w_1(z)$ and the second with probability $w_2(z)$. Assuming additive gene action on the phenotype, random mating, and that generations do not overlap, invasion fitness of a rare allele coding for $x_\bullet$ in a diploid population monomorphic for $x$ is given by

$$W(x_\bullet, x) = \frac{w_1(x_\bullet + x)\, w_2(x_\bullet + x)}{w_1(2x)\, w_2(2x)}. \tag{C-29}$$

Similarly to the scenario envisaged by eq. (C-28), individuals compete at random with respect to the environments they have experienced under eq. (C-29). This again entails that selection can never be diversifying here (shown straightforwardly following the approach described in Appendix A.2 with invasion fitness given by eq. C-29), and that polygenic variation cannot be maintained. This aligns with the findings in [27] that polymorphism in this model is difficult without dominance reversal at the phenotypic level (i.e. non-additive gene action on the phenotype), such that heterozygotes individuals can express the optimal trait value whenever they experience either environment in their lifetime. [27] argue that such dominance reversal is biologically unlikely.

### D Sex chromosomes

#### D.1 X-chromosome

In this section we investigate evolution of trait $z$ under the continuum-of-alleles model at a hemizygous locus on the non-recombining X-chromosome of a population with heteromorphic XY sex chromosomes (or Z-chromosome in ZW systems). As in many population genetic models of sexual antagonism on the X chromosome (e.g., [9, 28]), we initially assume that the Y-chromosome does not carry a functional copy of the gene and so does not contribute to variation in the trait $z$ (we relax this to consider the effect of a functional Y-gene in App. D.2).

To derive the invasion fitness relevant to our analysis, consider the case where two alleles a and A segregate at the X-linked locus of interest. Females may then express the phenotypes $z_{\mathrm{AA}}$, $z_{\mathrm{Aa}}$, $z_{\mathrm{aa}}$, where as in Appendix A.1, $z_u$ refers to the phenotype encoded by genotype $u \in \{\mathrm{AA}, \mathrm{Aa}, \mathrm{aa}\}$. By contrast, males are haploid for the X and so only express the phenotypes $z_{\mathrm{A}}$ or $z_{\mathrm{a}}$ (i.e., $z_u$ is the phenotype of genotype $u \in \{\mathrm{A}, \mathrm{a}\}$). Owing to male haploidy and sex-biased inheritance, the frequencies of A at generation $t$ in the male and female gametic pools given those frequencies at generation $t$ ($M(p_{\mathrm{m},t}, p_{\mathrm{f},t})$ and $F(p_{\mathrm{m},t}, p_{\mathrm{f},t})$ in eq. A-1) are

$$\begin{aligned}
M(p_{\mathrm{m},t}, p_{\mathrm{f},t}) &= p_{\mathrm{f},t}\frac{w_{\mathrm{m}}(z_{\mathrm{A}})}{\overline{w}_{\mathrm{m},t}} \\
F(p_{\mathrm{m},t}, p_{\mathrm{f},t}) &= p_{\mathrm{m},t}p_{\mathrm{f},t}\frac{w_{\mathrm{f}}(z_{\mathrm{AA}})}{\overline{w}_{\mathrm{f},t}} + \frac{1}{2}\big(p_{\mathrm{m},t}(1-p_{\mathrm{f},t}) + (1-p_{\mathrm{m},t})p_{\mathrm{f},t}\big)\frac{w_{\mathrm{f}}(z_{\mathrm{Aa}})}{\overline{w}_{\mathrm{f},t}},
\end{aligned} \tag{D-1}$$

where recall $w_v(z_u)$ is the number of gametes produced by an individual of sex $v \in \{\mathrm{f}, \mathrm{m}\}$ expressing the phenotype $z_u$ ($u \in \{\mathrm{AA}, \mathrm{Aa}, \mathrm{aa}, \mathrm{A}, \mathrm{a}\}$), and $\overline{w}_{\mathrm{m},t} = p_{\mathrm{f},t}w_{\mathrm{m}}(z_{\mathrm{A}}) + (1-p_{\mathrm{f},t})w_{\mathrm{m}}(z_{\mathrm{a}})$ and $\overline{w}_{\mathrm{f},t} = p_{\mathrm{m},t}p_{\mathrm{f},t}w_{\mathrm{f}}(z_{\mathrm{AA}}) + \big(p_{\mathrm{m},t}(1-p_{\mathrm{f},t}) + (1-p_{\mathrm{m},t})p_{\mathrm{f},t}\big)w_{\mathrm{f}}(z_{\mathrm{Aa}}) + (1-p_{\mathrm{m},t})(1-p_{\mathrm{f},t})w_{\mathrm{f}}(z_{\mathrm{aa}})$ are mean male and female fecundities at generation $t$, respectively [28]. In this case, the leading eigenvalue of the Jacobian matrix when $p_{\mathrm{m},t}$ and $p_{\mathrm{f},t}$ are both close to zero (eq. A-3) is

$$\lambda(0) = \frac{w_{\mathrm{f}}(z_{\mathrm{Aa}}) + \sqrt{w_{\mathrm{f}}(z_{\mathrm{Aa}})\left(w_{\mathrm{f}}(z_{\mathrm{Aa}}) + \frac{8w_{\mathrm{f}}(z_{\mathrm{aa}})w_{\mathrm{m}}(z_{\mathrm{A}})}{w_{\mathrm{m}}(z_{\mathrm{A}})}\right)}}{4w_{\mathrm{f}}(z_{\mathrm{aa}})}. \tag{D-2}$$

Writing $z_{\mathrm{A}} = x_\bullet$, $z_{\mathrm{a}} = x$, $z_{\mathrm{Aa}} = x_\bullet + x$ and $z_{\mathrm{aa}} = 2x$ in eq. (D-2), we find

$$W(x_\bullet, x) = \frac{w_{\mathrm{f}}(x_\bullet + x) + \sqrt{w_{\mathrm{f}}(x_\bullet + x)\left(w_{\mathrm{f}}(x_\bullet + x) + \frac{8w_{\mathrm{f}}(2x)w_{\mathrm{m}}(x_\bullet)}{w_{\mathrm{m}}(x_\bullet)}\right)}}{4w_{\mathrm{f}}(2x)} \tag{D-3}$$

for the invasion fitness for an X-linked mutant coding for effect $x_\bullet$ in a resident population monomorphic for $x$.

Plugging eq. (D-3) into eq. (A-6) we readily obtain the selection gradient acting at the X-linked locus,

$$S(x) = \frac{1}{3}\frac{w_{\mathrm{m}}'(x)}{w_{\mathrm{m}}(x)} + \frac{2}{3}\frac{w_{\mathrm{f}}'(x)}{w_{\mathrm{f}}(2x)}, \tag{D-4}$$

leading to a singular allelic value $x^*$ (i.e., such that $S(x^*) = 0$) whenever

$$\frac{w_{\mathrm{f}}'(2x^*)}{w_{\mathrm{f}}(2x^*)} = -\frac{1}{2}\frac{w_{\mathrm{m}}'(x^*)}{w_{\mathrm{m}}(x^*)}, \tag{D-5}$$

that is, when directional selection is opposing across the sexes and exactly half as strong in females than males.

Plugging eq. (A-6) into eq. (A-8) and eq. (D-3) into eq. (A-9), we find that convergence and evolutionary stability of the singular allelic value $x^*$ then depend on the sign of

$$S'(x^*) = \frac{4}{3}\left[\frac{w_{\mathrm{f}}''(2x^*)}{w_{\mathrm{f}}(2x^*)} + \frac{1}{4}\frac{w_{\mathrm{m}}''(x^*)}{w_{\mathrm{m}}(x^*)}\right] - \frac{4}{3}\left\{\left(\frac{w_{\mathrm{f}}'(2x^*)}{w_{\mathrm{f}}(2x^*)}\right)^2 + \frac{1}{4}\left(\frac{w_{\mathrm{m}}'(x^*)}{w_{\mathrm{m}}(x^*)}\right)^2\right\}, \tag{D-6}$$

and

$$H(x^*) = \frac{2}{3}\left[\frac{w_{\mathrm{f}}''(2x^*)}{w_{\mathrm{f}}(2x^*)} + \frac{1}{2}\frac{w_{\mathrm{m}}''(2x^*)}{w_{\mathrm{m}}(2x^*)}\right] - \frac{2}{3}\left\{\left(\frac{w_{\mathrm{f}}'(2x^*)}{w_{\mathrm{f}}(2x^*)}\right)^2 + \frac{1}{4}\left(\frac{w_{\mathrm{m}}'(2x^*)}{w_{\mathrm{m}}(2x^*)}\right)^2\right\}, \tag{D-7}$$

respectively (these conditions are interpreted in App. D.3).

#### D.2 Pseudo-autosomal regions

We next investigate evolution at a locus present in the pseudo-autosomal region of the X-chromosome ("PAR" [29]), assuming this locus recombines with an Y-linked non-recombining sex-determining-region at a rate $r \geq 0$ (with $r = 0.5$ this is equivalent to modelling an austosomal locus). Again, we consider our focal locus to be di-allelic with segregating alleles A and a. However, unlike in our analysis of the non-recombining X-chromosome, both males and females carry two functional copies of the focal gene and so may have genotype AA, Aa, or aa (leading to the phenotypes $z_u$ where $u \in \{\mathrm{AA}, \mathrm{Aa}, \mathrm{aa}\}$ in both sexes, as with an autosomally encoded trait).

For a locus in the PAR, an allele may be found in one of three chromosomal settings. First, it may be present in a female X-chromosome (having been either maternally or paternally inherited), second in a male X-chromosome (having been strictly maternally inherited), and third in a male Y-chromosome (having been strictly paternally inherited). We are therefore required to consider recursion equations of three frequencies of allele A at generation $t$: (1) its frequency in female gametes $p_{\mathrm{f},t}$, (2) its frequency in the male gametes carrying an X-chromosome $p^{\mathrm{X}}_{\mathrm{m},t}$, and (3) its frequency in the male gametes carrying a Y-chromosome $p^{\mathrm{Y}}_{\mathrm{m},t}$, i.e., $p^{u}_{\mathrm{m},t}$ is the frequency of A in sperm carrying a chromosome of type $u \in \{\mathrm{Y},\mathrm{X}\}$.

As with autosomes and non-recombining sex chromosomes, these frequencies are influenced by selection acting through adult male and female fecundities. Specifically, the frequency of allele A in each chromosome at generation at $t+1$ is

$$\begin{aligned} p^{\mathrm{X}}_{\mathrm{m},t+1} &= M^{\mathrm{X}}(p^{\mathrm{X}}_{\mathrm{m},t}, p^{\mathrm{Y}}_{\mathrm{m},t}, p_{\mathrm{f},t}) \\ p^{\mathrm{Y}}_{\mathrm{m},t+1} &= M^{\mathrm{Y}}(p^{\mathrm{X}}_{\mathrm{m},t}, p^{\mathrm{Y}}_{\mathrm{m},t}, p_{\mathrm{f},t}) \\ p_{\mathrm{f},t+1} &= F(p^{\mathrm{X}}_{\mathrm{m},t}, p^{\mathrm{Y}}_{\mathrm{m},t}, p_{\mathrm{f},t}), \end{aligned} \tag{D-8}$$

where

$$\begin{aligned} M^{\mathrm{X}}(p^{\mathrm{X}}_{\mathrm{m},t}, p^{\mathrm{Y}}_{\mathrm{m},t}, p_{\mathrm{f},t}) &= p_{\mathrm{f},t}\Big[\big(p^{\mathrm{X}}_{\mathrm{m},t}r + p^{\mathrm{Y}}_{\mathrm{m},t}(1-r)\big)\frac{w_{\mathrm{m}}(z_{\mathrm{AA}})}{\overline{w}_{\mathrm{m},t}} + \frac{1}{2}\big((1-p^{\mathrm{X}}_{\mathrm{m},t})r + (1-p^{\mathrm{Y}}_{\mathrm{m},t})(1-r)\big)\frac{w_{\mathrm{m}}(z_{\mathrm{Aa}})}{\overline{w}_{\mathrm{m},t}}\Big] \\ M^{\mathrm{Y}}(p^{\mathrm{X}}_{\mathrm{m},t}, p^{\mathrm{Y}}_{\mathrm{m},t}, p_{\mathrm{f},t}) &= \Big(p^{\mathrm{X}}_{\mathrm{m},t}r + p^{\mathrm{Y}}_{\mathrm{m},t}(1-r)\Big)\Big(p_{\mathrm{f},t}\frac{w_{\mathrm{m}}(z_{\mathrm{AA}})}{\overline{w}_{\mathrm{m},t}} + \frac{1}{2}(1-p_{\mathrm{f},t})\frac{w_{\mathrm{m}}(z_{\mathrm{Aa}})}{\overline{w}_{\mathrm{m},t}}\Big) \\ F(p^{\mathrm{X}}_{\mathrm{m},t}, p^{\mathrm{Y}}_{\mathrm{m},t}, p_{\mathrm{f},t}) &= \big(p^{\mathrm{X}}_{\mathrm{m},t}(1-r) + p^{\mathrm{Y}}_{\mathrm{m},t}r\big)p_{\mathrm{f},t}\frac{w_{\mathrm{f}}(z_{\mathrm{AA}})}{\overline{w}_{\mathrm{f},t}} + \frac{1}{2}\Big[\big(p^{\mathrm{X}}_{\mathrm{m},t}(1-r) + p^{\mathrm{Y}}_{\mathrm{m},t}r\big)(1-p_{\mathrm{f},t}) + \\ &\qquad \big((1-p^{\mathrm{X}}_{\mathrm{m},t})(1-r) + (1-p^{\mathrm{Y}}_{\mathrm{m},t})r\big)p_{\mathrm{f},t}\Big]\frac{w_{\mathrm{f}}(z_{\mathrm{Aa}})}{\overline{w}_{\mathrm{f},t}}, \end{aligned} \tag{D-9}$$

and mean male and female fecundities are

$$\begin{aligned} \overline{w}_{\mathrm{f},t} =& \frac{1}{2}(p^{\mathrm{X}}_{\mathrm{m},t} + p^{\mathrm{Y}}_{\mathrm{m},t})p_{\mathrm{f},t}w_{\mathrm{f}}(z_{\mathrm{AA}}) + \big(\frac{1}{2}(p^{\mathrm{X}}_{\mathrm{m},t} + p^{\mathrm{Y}}_{\mathrm{m},t})(1-p_{\mathrm{f},t}) + (1-p_{\mathrm{m},t})p_{\mathrm{f},t}\big)w_{\mathrm{f}}(z_{\mathrm{Aa}}) + \\ & (1-\frac{1}{2}(p^{\mathrm{X}}_{\mathrm{m},t} + p^{\mathrm{Y}}_{\mathrm{m},t})(1-p_{\mathrm{f},t})w_{\mathrm{f}}(z_{\mathrm{aa}}) \\ \overline{w}_{\mathrm{m},t} =& \frac{1}{2}(p^{\mathrm{X}}_{\mathrm{m},t} + p^{\mathrm{Y}}_{\mathrm{m},t})p_{\mathrm{f},t}w_{\mathrm{m}}(z_{\mathrm{AA}}) + \big(\frac{1}{2}(p^{\mathrm{X}}_{\mathrm{m},t} + p^{\mathrm{Y}}_{\mathrm{m},t})(1-p_{\mathrm{f},t}) + (1-p_{\mathrm{m},t})p_{\mathrm{f},t}\big)w_{\mathrm{m}}(z_{\mathrm{Aa}}) + \\ & (1-\frac{1}{2}(p^{\mathrm{X}}_{\mathrm{m},t} + p^{\mathrm{Y}}_{\mathrm{m},t})(1-p_{\mathrm{f},t})w_{\mathrm{m}}(z_{\mathrm{aa}}) \end{aligned} \tag{D-10}$$

This leads to the following Jacobian matrix when A is rare ($p^{\mathrm{Y}}_{\mathrm{m},t}$, $p^{\mathrm{X}}_{\mathrm{m},t}$, and $p_{\mathrm{f},t}$ are small):

$$\mathrm{J}(0) = \begin{pmatrix} \dfrac{\partial M^{\mathrm{X}}(p^{\mathrm{X}}_{\mathrm{m},t}, p^{\mathrm{Y}}_{\mathrm{m},t}, p_{\mathrm{f},t})}{\partial p^{\mathrm{X}}_{\mathrm{m},t}} & \dfrac{\partial M^{\mathrm{X}}(p^{\mathrm{X}}_{\mathrm{m},t}, p^{\mathrm{Y}}_{\mathrm{m},t}, p_{\mathrm{f},t})}{\partial p^{\mathrm{Y}}_{\mathrm{m},t}} & \dfrac{\partial M^{\mathrm{X}}(p^{\mathrm{X}}_{\mathrm{m},t}, p^{\mathrm{Y}}_{\mathrm{m},t}, p_{\mathrm{f},t})}{\partial p_{\mathrm{f},t}} \\ \dfrac{\partial M^{\mathrm{Y}}(p^{\mathrm{X}}_{\mathrm{m},t}, p^{\mathrm{Y}}_{\mathrm{m},t}, p_{\mathrm{f},t})}{\partial p^{\mathrm{X}}_{\mathrm{m},t}} & \dfrac{\partial M^{\mathrm{Y}}(p^{\mathrm{X}}_{\mathrm{m},t}, p^{\mathrm{Y}}_{\mathrm{m},t}, p_{\mathrm{f},t})}{\partial p^{\mathrm{Y}}_{\mathrm{m},t}} & \dfrac{\partial M^{\mathrm{Y}}(p^{\mathrm{X}}_{\mathrm{m},t}, p^{\mathrm{Y}}_{\mathrm{m},t}, p_{\mathrm{f},t})}{\partial p_{\mathrm{f},t}} \\ \dfrac{\partial F(p^{\mathrm{X}}_{\mathrm{m},t}, p^{\mathrm{Y}}_{\mathrm{m},t}, p_{\mathrm{f},t})}{\partial p^{\mathrm{X}}_{\mathrm{m},t}} & \dfrac{\partial F(p^{\mathrm{X}}_{\mathrm{m},t}, p^{\mathrm{Y}}_{\mathrm{m},t}, p_{\mathrm{f},t})}{\partial p^{\mathrm{Y}}_{\mathrm{m},t}} & \dfrac{\partial F(p^{\mathrm{X}}_{\mathrm{m},t}, p^{\mathrm{Y}}_{\mathrm{m},t}, p_{\mathrm{f},t})}{\partial p_{\mathrm{f},t}} \end{pmatrix}_{p^{\mathrm{Y}}_{\mathrm{m},t}=p^{\mathrm{X}}_{\mathrm{m},t}=p_{\mathrm{f},t}=0}, \quad \text{(D-11)}$$

with A invading when the leading eigenvalue of this matrix is greater than unity. This eigenvalue turns out to be complicated and we do not give it here, only noting that, consistent with eqs. A-4 and D-2, it depends on the phenotypes $z_{\mathrm{Aa}}$ and $z_{\mathrm{aa}}$ but not $z_{\mathrm{AA}}$ (given that A is rare).

Assuming the continuum-of-alleles, i.e., writing $z_{\mathrm{Aa}} = x^* + x$ and $z_{\mathrm{aa}} = 2x$, the leading eigenvalue of eq. (D-11) gives the invasion fitness of a rare PAR mutant encoding $x_\bullet$ for $r > 0$ (the case of $r = 0$ is discussed next). By plugging this into eq. (A-6) we find that the selection gradient $S(x)$ for a PAR-gene is in fact identical to that at an autosomal locus (eq. A-6), leading to the same singular allelic value $x^*$ and condition for convergence stability (eqs. A-11 and A-12). The condition for disruptive selection on the PAR, however, diverges from the autosomes, such that selection is stabilising or diversifying according to the sign of

$$H(x^*) = \frac{w''_{\mathrm{f}}(2x^*)}{w_{\mathrm{f}}(2x^*)} + \frac{w''_{\mathrm{m}}(2x^*)}{w_{\mathrm{m}}(2x^*)} + \frac{1}{2}\left(\frac{1}{r} - 2\right)\left[\left(\frac{w'_{\mathrm{f}}(2x^*)}{w_{\mathrm{f}}(2x^*)}\right)^2 + \left(\frac{w'_{\mathrm{m}}(2x^*)}{w_{\mathrm{m}}(2x^*)}\right)^2\right], \quad \text{(D-12)}$$

which is increasingly likely to be positive (leading to diversifying selection) as $r$ tends to 0 and linkage with the sex-determining region tightens (Appendix D.3 for interpretation).

For loci within the non-recombining region of the sex chromosome ($r = 0$), the Jacobian matrix (D-11) becomes block diagonal, with a $1 \times 1$ matrix that describes invasion fitness of new alleles that arise on the Y chromosome and a $2 \times 2$ matrix that describes invasion fitness of new alleles on the X. Consistent with our conclusions from eq. (D-12) as $r$ tends to 0, we find that fixation of a single allele encoding the same trait value on both the X and Y (i.e., a singular state) is always invadable, indicating that selection is diversifying, unless the fitness optimum is the same in males and females. Specifically, the leading eigenvalue of the $1 \times 1$ matrix for the Y is given by $w_{\mathrm{m}}(z_{\mathrm{Aa}})/w_{\mathrm{m}}(z_{\mathrm{aa}})$, implying that the only uninvadable allele A on the Y must maximize male fitness. If this A allele is also fixed on the X, then it can be invaded by small-effect mutations that increase fitness in females (i.e., the leading eigenvalue of the $2 \times 2$ matrix will always be greater than one for alleles that slightly increase $z$ or for alleles that slightly decrease $z$, unless females have the same fitness optimum). Thus, selection at homologous genes on the X and Y in the SDR ($r = 0$) is always diversifying with sexually antagonistic selection.

#### D.3 Comparisons across genomic regions

In order to more easily compare the conditions for diversifying selection in the continuum-of-alleles model across autosomes, the X-chromosome, and the PAR, it is helpful to decompose $H(x^*)$ and $S'(x^*)$ in the following ways. First we write $S'(x^*)$ as

$$S'(x^*) = H(x^*) + I(x^*) \tag{D-13}$$

where

$$I(x^*) = \left. \frac{\partial^2 W(x_\bullet, x)}{\partial x_\bullet \partial x} \right|_{x_\bullet = x = x^*} \tag{D-14}$$

[30, 31], and recall $W(x_\bullet, x)$ is the invasion fitness of a rare allelic mutant $x_\bullet$ in a resident allele population with effect $x$ (e.g., eq. A-5 for a mutant at an autosomal gene and eq. D-3 for an X-linked mutant). The term $I(x^*)$ captures frequency-dependent selection [5], i.e., the fitness effect of a joint increase in the mutant and resident allelic value when the resident allele encodes the singularity $x^*$. Owing to the fact that evolutionary branching occurs when $S'(x^*) < 0$ and $H(x^*) > 0$, this also requires that $I(x^*) < 0$, such that selection is negatively frequency-dependent.

Next we write $H(x^*)$ as

$$H(x^*) = 2h_{\text{q}}(x^*) + h_{\text{w}}(x^*), \tag{D-15}$$

where

$$h_{\text{q}}(x^*) = \sum_{i=1}^{M} \sum_{j=1}^{M} v_i^\circ(x) \left. \frac{\partial \omega_{ij}(x_\bullet, x)}{\partial x_\bullet} \right|_{x_\bullet = x = x^*} \left. \frac{\partial q_j(x_\bullet, x)}{\partial x_\bullet} \right|_{x_\bullet = x = x^*} \tag{D-16}$$

and

$$h_{\text{w}}(x^*) = \sum_{i=1}^{M} \sum_{j=1}^{M} v_i^\circ(x) \left. \frac{\partial^2 \omega_{ij}(x_\bullet, x)}{\partial x_\bullet^2} \right|_{x_\bullet = x = x^*} q_j^\circ(x), \tag{D-17}$$

([31] eqs. 3.4-3.5). In eq. (D-17), $\omega_{ij}(x_\bullet, x)$ is entry-$(i, j)$ of the invasion matrix J(0) (eq. A-3 for autosomes and X-linked loci and eq. D-11 for PAR genes with each phenotype given in terms of $x_\bullet$ and $x$, which is equivalent to the mean matrix of a multitype branching process), and $M$ is the number of genomic contexts a gene may find itself in, corresponding to the length of each dimension of the invasion matrix. For example, $M = 2$ for autosomes and X-chromosomes as a gene may be present in a male or in female chromosome, while $M = 3$ for PAR genes as a gene may be present in a female X chromosome, in a male X chromosome, or a male Y chromosome. The term $q_j(x_\bullet, x)$ is the $j^{\text{th}}$ entry of the right eigenvector $\boldsymbol{q}(x_\bullet, x)$ of the invasion matrix normalised such that is sums to 1 (i.e., $\sum_{j=1}^{M} q_j(x_\bullet, x) = 1$), which corresponds to the asymptotic frequency of mutants in each genomic context $j$. Finally, the terms $q_j^\circ(x)$ and $v_i^\circ(x)$ refer respectively to the $j^{\text{th}}$ and $i^{\text{th}}$ entries of the right $\boldsymbol{q}^\circ(x)$ and left $\boldsymbol{v}^\circ(x)$ eigenvectors of the invasion matrix evaluated at neutrality (i.e., J(0) with all entries evaluated at $x_\bullet = x$). In addition, these vectors have been normalised such that $\boldsymbol{q}^\circ(x) \cdot \boldsymbol{v}^\circ(x) = 1$. Here, $v_i^\circ(x)$ corresponds to the normalised reproductive value of a gene in genomic context $i$ in the absence of selection.

Intuitively, the quantity $h_{\text{w}}(x^*)$ captures the effect of non-linear fitness responses to a change in $x$ owing to the curvature in the fitness landscape in each sex around $x^*$. Specifically, $h_{\text{w}}(x^*)$ is positive, and so favours disruptive selection, in the presence of accelerating fitness returns (i.e., a positively curved fitness landscape). Meanwhile, $h_{\text{q}}(x^*)$ captures synergistic "context × fitness" effects of allelic expression, which here refers to the fitness effect of a change in allelic value on a mutant's fitness in a given genomic context multiplied by the probability that a mutant will be present in that context. If positive, this term indicates that the phenotypic effect encoded by a mutant allele increases the representation of this allele in the genomic context in which it is beneficial, so favouring disruptive selection ([31] for discussion).

From eqs. (D-13) and (D-15) we can write the conditions for evolutionary branching at $x^*$ (i.e., $S'(x^*) < 0$ and $H(x^*) > 0$) as

$$-2h_{\text{q}}(x^*) < h_{\text{w}}(x^*) < -I(x^*) - 2h_{\text{q}}(x^*). \tag{D-18}$$

Written in this form, the condition for diversifying selection (given previously in eq. A-14) at an autosomal locus is

$$\underbrace{0}_{-2h_{\text{q}}(x^*)} < \underbrace{\frac{1}{2}\left[\frac{w_{\text{f}}''(z^*)}{w_{\text{f}}(z^*)} + \frac{w_{\text{m}}''(z^*)}{w_{\text{m}}(z^*)}\right]}_{h_{\text{w}}(x^*)} < \underbrace{\left(\frac{w_{\text{f}}'(z^*)}{w_{\text{f}}(z^*)}\right)^2 + \left(\frac{w_{\text{m}}'(z^*)}{w_{\text{m}}(z^*)}\right)^2 - \frac{1}{2}\left[\frac{w_{\text{f}}''(z^*)}{w_{\text{f}}(z^*)} + \frac{w_{\text{m}}''(z^*)}{w_{\text{m}}(z^*)}\right]}_{-I(x^*)} + \underbrace{0}_{-2h_{\text{q}}(x^*)}. \tag{D-19}$$

This tells us that there are no synergistic effects arising from genomic context at an autosomal locus, due to these loci showing symmetric inheritance across males and females. Instead disruptive selection emerges purely from non-linear fitness response to trait change (i.e., from an accelerating fitness landscape), with strong selection driving negative-frequency dependence through competition within the sexes.

For an X-linked locus, diversifying selection occurs when

$$\underbrace{\frac{2}{3}\left[\left(\frac{w_{\mathrm{f}}'(z^*)}{w_{\mathrm{f}}(z^*)}\right)^2+\frac{1}{4}\left(\frac{w_{\mathrm{m}}'(z^*)}{w_{\mathrm{m}}(z^*)}\right)^2\right]}_{-2h_{\mathrm{q}}(x^*)}<\underbrace{\frac{2}{3}\left[\frac{w_{\mathrm{f}}''(z^*)}{w_{\mathrm{f}}(z^*)}+\frac{1}{2}\frac{w_{\mathrm{m}}''(z^*)}{w_{\mathrm{m}}(z^*)}\right]}_{h_{\mathrm{w}}(x^*)}<\underbrace{\frac{4}{3}\left[\left(\frac{w_{\mathrm{f}}'(z^*)}{w_{\mathrm{f}}(z^*)}\right)^2+\frac{1}{4}\left(\frac{w_{\mathrm{m}}'(z^*)}{w_{\mathrm{m}}(z^*)}\right)^2\right]-\frac{2}{3}\frac{w_{\mathrm{f}}''(z^*)}{w_{\mathrm{f}}(z^*)}}_{-I(x^*)-2h_{\mathrm{q}}(x^*)}. \tag{D-20}$$

From this we can see there are two routes for diversifying selection on the X-chromosome. First, such selection can occur under weak sexually antagonistic selection (i.e., such that terms $[w_{\mathrm{m}}'(x^*)/w_{\mathrm{m}}(x^*)]^2$ and $[w_{\mathrm{f}}'(2x^*)/w_{\mathrm{f}}(2x^*)]^2$ are negligible) when

$$0<-2\frac{w_{\mathrm{f}}''(2x^*)}{w_{\mathrm{f}}(2x^*)}<\frac{w_{\mathrm{m}}''(x^*)}{w_{\mathrm{m}}(x^*)}<-4\frac{w_{\mathrm{f}}''(2x^*)}{w_{\mathrm{f}}(2x^*)}, \tag{D-21}$$

that is, when the fitness landscape around $x^*$ is positively curved in males and negatively curved in females to a particular degree. This is due to female dominance patterns induced by these landscapes that are conducive to X-linked polymorphism [9, 28]. While this in principle indicates that balanced sexually antagonistic polymorphism may be maintained on the X-chromosome without strong selection, these curvature conditions are restrictive, however, and are rarely fulfilled with the power functions explored here Appendix (Fig. 10A) and never fullfilled with Gaussian functions (Fig. 10C). Second, as with autosomes, diversifying selection can arise from strong sexually antagonistic selection on the X-chromosome when the fitness landscape is accelerating at $x^*$ ($h_{\mathrm{w}}(x^*)>0$). However, this condition is typically more restrictive on the X-chromosome than autosomes (Appendix Fig. 10B-C) owing to the fact that $h_{\mathrm{q}}(x^*)$ is negative, i.e., sexually antagonistic X-linked loci show a negative context×fitness synergy (see main text section 3.3 for interpretation).

Finally, conditions for diversifying selection on the PAR are

$$\underbrace{\frac{1}{2}\left(2-\frac{1}{r}\right)\left[\left(\frac{w_{\mathrm{f}}'(z^*)}{w_{\mathrm{f}}(z^*)}\right)^2+\left(\frac{w_{\mathrm{m}}'(z^*)}{w_{\mathrm{m}}(z^*)}\right)^2\right]}_{-2h_{\mathrm{q}}(x^*)}<\underbrace{\frac{1}{2}\left[\frac{w_{\mathrm{f}}''(z^*)}{w_{\mathrm{f}}(z^*)}+\frac{w_{\mathrm{m}}''(z^*)}{w_{\mathrm{m}}(z^*)}\right]}_{h_{\mathrm{w}}(x^*)}< \underbrace{\left(\frac{w_{\mathrm{f}}'(z^*)}{w_{\mathrm{f}}(z^*)}\right)^2+\left(\frac{w_{\mathrm{m}}'(z^*)}{w_{\mathrm{m}}(z^*)}\right)^2-\frac{1}{2}\left[\frac{w_{\mathrm{f}}''(z^*)}{w_{\mathrm{f}}(z^*)}+\frac{w_{\mathrm{m}}''(z^*)}{w_{\mathrm{m}}(z^*)}\right]}_{-I(x^*)-2h_{\mathrm{q}}(x^*)}. \tag{D-22}$$

Eq. (D-22) reveals that with $r=0.5$ (i.e., loose linkage with the SDR), the criteria for polymorphism at a pseudoautosomal locus is equivalent to an autosomal locus (i.e., eq. D-22 reduces to eq. D-19), such that diversifying selection requires strong sexual antagonism and an accelerating fitness landscape. However, as $r$ decreases, PAR genes become significantly more amenable to sexually antagonistic polymorphism. This is because weak recombination between these loci and the SDR leads to a positive context×fitness synergy and so permissive conditions for diversifying selection (main text section 3.3 for interpretation). Specifically, selection will be diversifying whenever

$$r < \frac{1}{2} \frac{\left(\frac{w_{\mathrm{f}}'(2x^*)}{w_{\mathrm{f}}(2x^*)}\right)^2 + \left(\frac{w_{\mathrm{m}}'(2x^*)}{w_{\mathrm{m}}(2x^*)}\right)^2}{\left(\frac{w_{\mathrm{f}}'(2x^*)}{w_{\mathrm{f}}(2x^*)}\right)^2 + \left(\frac{w_{\mathrm{m}}'(2x^*)}{w_{\mathrm{m}}(2x^*)}\right)^2 - \left[\frac{w_{\mathrm{f}}''(2x^*)}{w_{\mathrm{f}}(2x^*)} + \frac{w_{\mathrm{m}}''(2x^*)}{w_{\mathrm{m}}(2x^*)}\right]}. \tag{D-23}$$

This tells us that with tight linkage, polymorphism can arise on both accelerating and diminishing landscapes and, if recombination is sufficiently weak, in the absence of strong sexual antagonism (requires $r \sim \mathcal{O}(s^2)$ where $s = w_{\mathrm{m}}'(2x^*)/w_{\mathrm{m}}(2x^*) = w_{\mathrm{f}}'(2x^*)/w_{\mathrm{f}}(2x^*)$ is the strength of directional selection in each sex).

### E Genomic tests for sexually antagonistic loci

In this appendix, we describe the procedures used to characterise genetic variation under sexual antagonism using methods from population genomics (and generate Fig. 4). We applied two approaches to simulated data: (i) we calculated between-sex $F_{\mathrm{ST}}$, a measure of allelic differentiation; and (ii) we calculated the sex-difference in the covariance between within-individual allele frequency and fitness. Calculations were performed on 1000 replicates after a burn-in of $10^5$ generations (in order to provide mutation-selection-drift equilibrium levels of variation). We considered the case where the trait is encoded by ten loci, $L = 10$, and where the total population size is $N = 10{,}000$ (consisting of 5,000 females and 5,000 males).

#### E.1 Allelic differentiation

We computed between-sex allelic differentiation at each locus $k \in \{1, \dots, 10\}$ when polymorphic as

$$F_{\mathrm{ST},k} = \frac{(\nu_{k,\mathrm{f}} - \nu_{k,\mathrm{m}})^2}{4\nu_k(1-\nu_k)}, \tag{E-1}$$

[32, 33] where

$$\nu_{k,u} = \frac{1}{N}\sum_{i=1}^{N} \nu_{k,u,i}, \tag{E-2}$$

where $v_{k,u,i} \in \{0, 0.5, 1\}$ is the within-individual frequency of allele A in the parent of sex $u \in \{f, m\}$ of recruited offspring $i \in \{1, 2, ..., N\}$, and $v_k = (v_{k,m} + v_{k,f})/2$. In other words, $v_{k,f}$ and $v_{k,m}$ are the frequencies of allele A among successful female and male parents respectively, weighted by the number of recruited offspring these parents produced. Eq. (E-1) thus gives a measure of differentiation owing to fecundity selection [34].

We calculated $F_{ST,k}$ in generation $10^5$ for each locus $k \in \{1, \ldots, 10\}$ that was polymorphic with a minor allele frequency greater that 0.05 and for each replicate simulation. We then pooled all $F_{ST,k}$ values across loci and across simulation replicates, and then sampled 1000 values at random for each treatment, which were used to plot Fig. 4A (this sampling ensured the same number of data points for each treatment). For control, this procedure was then repeated but with the sex of each individual randomly assigned (so that the variable $u$ in each $v_{k,u,i}$ is a Bernoulli distributed random variable with $u = f$ with probability 1/2 and $u = m$ otherwise).

##### E.2 Associations tests

We also computed the difference in the linear regression slope of reproductive success on the frequency of allele A, separately in males and females. In generation $10^5$, for each individual $i \in \{1, \ldots, 5{,}000\}$ of sex $u \in \{f, m\}$, we measured the number $\omega_{u,i}$ of offspring this individual produced and the frequency $p_{k,u,i} \in \{0, 0.5, 1\}$ of allele A it carries at each locus $k \in \{1, \ldots, 10\}$. For each sex $u$ and each locus $k$ that is polymorphic with minor allele frequency greater than 0.05, we then fitted a Generalised Linear Model with Poisson error distribution, with reproductive success $\omega_{u,i}$ as a response variable and allele frequency $p_{k,u,i}$ as a fixed effect, to estimate the regression slope $\beta_{k,u}$. Finally, we calculated the absolute between-sex difference in the regression slopes $\Delta\beta_k = |\beta_{k,m} - \beta_{k,f}|$.

As with between-sex $F_{ST}$, we pooled all $\Delta\beta_k$ values across loci and replicates, and then sampled 1000 values at random for each treatment, which were used to plot Fig. 4B (this sampling ensured the same number of data points for each treatment). For control, this procedure was then repeated but with the sex of each individual randomly assigned (so that the variable $u$ in each $v_{k,u,i}$ is a Bernoulli distributed random variable with $u = f$ with probability 1/2 and $u = m$ otherwise).

### F Selection Gradient Estimates

In this Appendix, we compare our results for the conditions required to maintain elevated autosomal heterozygosity across loci, and estimates of sex-specific selection gradients from natural populations. First, we used data from a recent meta-analysis of 634 sex-specific directional selection gradient estimaes from wild populations [35] (data available at: doi.org/10.5061/dryad.b0bp357) to find candidate traits that could be under diversifying selection (assuming an autosomal genetic basis). To do this, we calculated the proportion of trait estimates that could be at an evolutionary equilibrium, i.e., showed directional selection gradients [3] of opposing sign but similar magnitude, such that the weaker gradient was 90% the size of the stronger one (i.e., $-0.9 > \text{gradient}_\text{m}/\text{gradient}_\text{f} > -1/0.9$, where $\text{gradient}_u$ is the directional selection gradient estimated in sex $u$). We found 16 traits which fulfilled this criteria, and for each trait we referred to the original paper to find the quadratic selection gradients, if they were calculated, before plugging all four gradients (directional and quadratic gradients in males and females) into inequality (4b). We found that none of the traits for which directional and quadratic gradients were available satisfied eq. (4b), indicating that none of them show fitness variation consistent with diversifying selection.

Second, we applied the same procedure to a recent collection of directional and quadratic selection gradients estimates from a large human dataset [36] (data available as supporting information at: doi.org/10.1073/pnas.1707227114, Dataset S1). In this case, no traits appeared to be sexually antagonistic and at an evolutionary equilibrium.

### G Supplementary Discussion

In this section, we discuss in more detail our modelling assumptions, as well as the relationship between our conclusions and existing results on the maintenance of genetic variation.

#### G.1 Genetic assumptions

Throughout our study, we take an idealised view of the genetic basis of phenotypic variation, with the specific aim of generating the most conducive conditions for sexually antagonistic polymorphism. In particular, we made two key assumptions to ensure that sex-specific selection on the trait $z$ leads to the strongest possible sexual conflict. First, we assumed that $z$ has a purely genetic basis, i.e., that there are no environmental effects on trait expression. In reality, quantitative traits overwhelmingly show environmental dependence [37, 38], and this is especially true for sexual phenotypes, which are commonly condition-dependent [39–42]. By producing non-heritable variation, environmental effects should dampen the impact of sexually antagonistic selection on genetic variation, making it less likely for such selection to be able to maintain elevated polymorphism. Second, we assumed a perfect genetic correlation between males and females, i.e., that alleles had completely sex-concordant phenotypic effects (although see App. C.6 for discussion of antagonism arising from sex-specific allelic effects). This is likely true at the onset of sex-specific selection [18, 43, 44], but may be relaxed if alleles appear that are differentially expressed, or encode phenotypic effects with different directions, across the sexes – as they will be favoured by selection. Consequently, complete or partial sexual dimorphism can evolve, thus ameliorating sexual antagonism and so reducing the potential for balancing selection [21, 45, 46]. In sum, by excluding environmental variation and sex-dependent allelic effects, our model therefore represents a best-case scenario for the maintenance of sexually antagonistic polymorphism, indicating our main conclusion (i.e., that sexually antagonistic polymorphism is rarely favoured) is conservative.

Additionally, to simplify analysis, our polygenic model makes some further assumptions that are in line with classic quantitative genetics [14, 47]. In particular, we considered $z$ to be encoded by freely recombining loci with equal phenotypic effects. In Appendices C.1-C.2 we show our results are robust to deviations from these baseline assumptions. Specifically, variation in effect sizes ($\delta_k$) simply alters the phenotypes expressed by different homozygote genotypes. Under stabilising selection, this may affect which genotypes are close to the optimum $z^*$ and therefore whether any loci or which locus eventually remains polymorphic. But it is still the case that ultimately a maximum of one locus experiences balancing selection. Similarly, decreasing the recombination rate between loci ($r < 0.5$) has little effect on the conditions for polymorphism, with the exception of extremely tight linkage relative to selection. In this case, polymorphism across multiple loci is more easily maintained under stabilising selection, as polygenic haplotypes behave increasingly like a single locus and the behaviour of the system approaches the classic single-locus model.

Another important genetic assumption in our polygenic model was that loci have additive effects on the phenotype. Epistatic genetic effects on fitness therefore arise due to the mapping of phenotype to fitness, but these are an emergent property. Introducing arbitrary epistatic variation directly at the phenotypic level (i.e., non-additive phenotypic effects caused by specific interactions between genes [48, 49]) will alter the fitness of different genotypes and thus may impact the conditions for maintaining polymorphism [50–52]. Indeed, such epistatic effects explain some of the discrepancies between our results and those from previous two-locus diallelic models of sexual antagonism, where genotypic fitness follows a sex-specific multiplicative model [10, 12] (with the addition of multiplicative epistasis in [53]). The increased genetic complexity of these models generates more permissive conditions for polymorphism because double heterozygotes can experience an intrinsic fitness advantage under certain parameter values, which does not arise in our additive quantitative trait approach. (Such a pattern of heterozygote advantage could arise from a genotype-phenotype map that incurs sex-specific phenotypic dominance reversal across multiple loci, although biological mechanisms would produce this are likely to be rare [27].) While it is unclear how easily such multiplicative fitness effects can extend over many loci for the same trait, these results suggest that there may exist scenarios where polymorphism is more readily favoured than in our model.

##### G.2 Ecological assumptions

In this study, we considered a large well-mixed population in which males and females mate randomly. Departures from such panmixia are common even in dioecious species, which often show some degree of assortative, or non-random, mating, e.g., with respect to a phenotype or inbreeding owing to spatial subdivision. Previous single-locus population genetics models have found that, by modulating the frequency of heterozygotes, these mating patterns can alter the conditions for sexually antagonistic variation to be maintained (i.e., by influencing the capacity of dominance-reversal to favour polymorphism [54–57]). However, these effects do not influence the segregation of quantitative variation in polygenic traits, due to the diminished role that fitness dominance plays in the maintenance of sexually antagonistic variation across loci. Instead, the importance of reproductive ecology on polymorphism in these traits will be through its effect on fitness variation within each sex, and thus on the strength of sexual antagonism. For example, in a population with female-limited reproduction, assortative mating for genotype could reduce the strength of selection in males if it meant maladapted (i.e., low fecundity) males tended to show elevated mating success with well-adapted (i.e., high fecundity) females [54]. Conversely, the opposite would be true if assortment was on fecundity, so that low fecundity males additionally show diminished mating success with high fecundity females [56]. Meanwhile, in spatially subdivided populations, limited dispersal will lead to inbreeding and kin competition, i.e., competition between phenotypically similar relatives, which can weaken selection in both sexes according to the sex-specific dispersal rate [57]. In all these cases, effects on the strength of selection in males and females would influence the intensity of sexual antagonism and so the maintenance of polymorphism. We therefore suggest that investigating the potential for diversifying selection in sexually antagonistic traits under different models of spatial structure, mate choice and mating benefits (including cases where both sexes gain a benefit from mating with high fitness partners) may provide a fruitful avenue for future research.

Consistent with existing theory of sex-specific selection [e.g. 8, 18, 20], our model assumes that competition (fecundity or survival) occurs within sexes. This allows for frequency-dependence and so, consequently, diversifying selection. Our models of sexual antagonism are therefore analogous to those of heterogenous environments connected by dispersal where selection is soft (i.e., density regulation occurs within patches, prior to the pooling of individuals [58], e.g., [59–64], see App. C.6). In these models, polymorphism owing to diversifying selection requires dispersal fall below a critical threshold relative to selection, otherwise selection is stabilising for a single optimum and so disfavours allelic variation. In fact, the condition for diversifying selection in a soft selection model of local adaptation with two patches connected by a dispersal probability $m$ is equivalent to the condition for diversifying selection owing to sexual antagonism on the PAR. This is straightforward to show by following the steps given in App. A.2 with eq. (C-25) in place of eq. (A-5), and leads to identical expressions as eq. (D-22) and eq. (D-23), with $r$ replaced by $m$, and $w_{\mathrm{m}}(z)$ and $w_{\mathrm{f}}(z)$ by $w_1(z)$ and $w_2(z)$ (which are the fecundities of expressing $z$ in each patch). Consequently, when dispersal is random ($m = 0.5$), the condition for polymorphism is equivalent to the case of sexual antagonism at an autosomal locus. Meanwhile, when dispersal is limited, $m < 0.5$, polymorphism can more easily be maintained as reduced gene flow allows selection to drive associations that lead alleles to be over represented in the patch they are adaptive in, as in sexual antagonism at pseudoautosomal loci. From this perspective, it is then not so surprising that only very strong sexually antagonistic selection can favour autosomal variation, but that polymorphism may be maintained easily in a PAR.

The maintenance of genetic variation (albeit under potentially restrictive conditions) in such soft selection models contrasts with those of hard selection, that is, where competition occurs at the global scale, e.g., where selection acts after the pooling of individuals in a heterogenous environment, or collectively amongst males and females in the case of sexual antagonism. Here, polymorphism is significantly less likely, owing to lack of negative frequency-dependent selection [58]. In particular, diversifying selection never occurs under hard selection in the presence of full mixing [26, 65] (i.e., random dispersal, or sex-symmetric inheritance of an autosomal allele, App. C.6). Owing to the nature of sex-specific selection, it is difficult to see how such a scenario could be captured by a hard selection model, because competition over reproduction necessarily occurs within sexes. Nevertheless, by modelling a purely soft selection case, our analyses again provide the most generous conditions for sexually antagonistic polymorphism.

As in models of local adaptation, we find that when selection is not strong enough to be diversifying, sexual antagonism erodes polymorphism across loci through stabilising or directional selection on the trait. This finding is consistent with existing theory of concordant selection, i.e., in the absence of different optimal trait values between sexes or environments. Such models predict that stabilising selection erodes genetic variation (reviewed in [16, 25, 66], chap. 28 in [14], see also [67] for fluctuating selection) and that, in the absence of tight linkage between loci, selection typically maintains a maximum of one balanced polymorphism (e.g., [16, 25, 68–71] – genetic variation across loci then segregates only ephemerally, owing to recurrent mutation and drift [72]). Selection diminishes variation here because heterozygote genotypes are more easily broken down by Mendelian segregation, resulting in a wider distribution of phenotypes among offspring, some of which deviate from the optimum under stabilising selection and so are less fit. Our results therefore reinforce the general notion that, whatever its ecological source, stabilising selection erodes variation by favouring intermediate homozygote genotypes over intermediate heterozygote genotypes.

Single-locus population genetics theory has highlighted that, even in the absence of balancing selection, genetic variation can persist more easily at mutation-selection-drift balance when selection is sexually antagonistic versus sexually concordant [73, 74]. This is because directional selection at a di-allelic locus (i.e., favouring the fixation of one allele the other) is – all else equal – weaker when this selection arises from antagonism rather than concordance, as directional selection within each sex is acting in different directions. This effect can also be seen from the general selection gradient on a trait experiencing sex-specific selection (eq. A-10), which indicates that the strength of directional selection on a shared male-female phenotype is weakened when selection in each sex acts in different directions (i.e., when $w_\mathrm{m}'(z)/w_\mathrm{m}(z)$ and $w_\mathrm{f}'(z)/w_\mathrm{f}(z)$ are of opposing sign). Consequently, convergence of a population to a singular phenotype $z^*$ should be slower under sexually antagonistic than sexually concordant selection and levels of standing variation during this phase will be slightly higher at mutation-selection-drift balance (as directional selection is less efficient at fixing alleles to drive the population towards an equilibrium). However, once the population reaches $z^*$, the strength of stabilising selection depends only on the curvature of the male and female fitness landscapes (eq. A-13) and thus is independent of the direction of selection acting in each sex. Consequently, once a population experiences stabilising selection, there is no reason to expect the levels of standing variation will fundamentally differ depending on whether selection across the sexes is antagonistic or concordant.

#### Appendix Figures

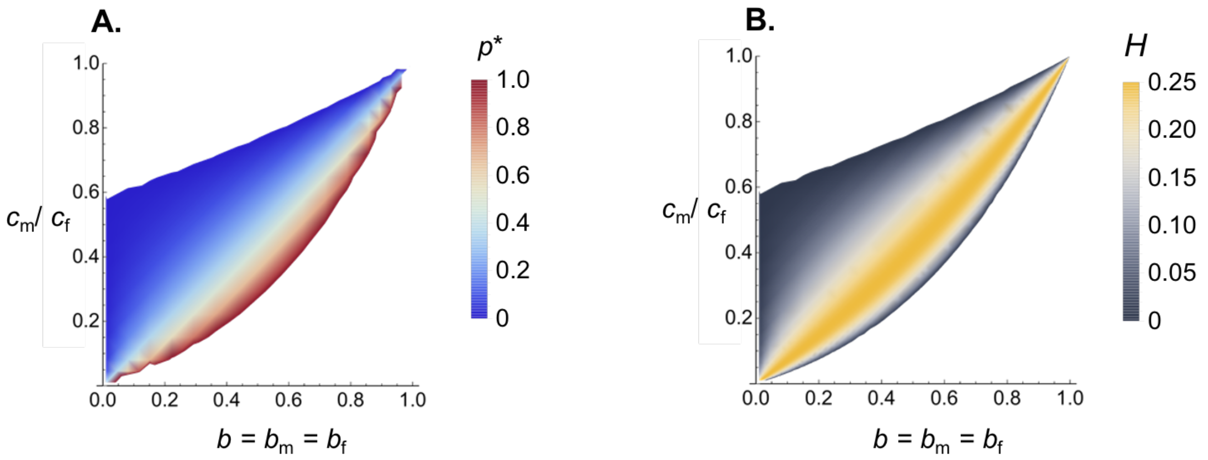


**Appendix Figure 1: Genetic variation when one locus is polymorphic in the two-locus model.** Panels show mathematical results from our di-allelic models with two loci ($L = 2$) when male and female fecundity follow power functions and one locus is fixed for one allele (eq. I.A, Box I, see Appendix B.2 for analysis, other parameters: $c_m = 0.1, \delta = 1$). Panel **A** shows the equilibrium frequency of allele $A_1$ ($p_1^*$, eq. B-9) at locus 1 when $z$ is encoded by two di-allelic loci, given that the second locus is fixed for $a_2$, as a function of the shape of fecundity curves ($b$) and strength of fecundity costs in males relative to females ($c_m/c_f$). Panels **B** shows average heterozygosity across both loci when $A_1$ is at a polymorphic equilibrium ($H = 1 - \sum_{k=1}^{2}(1-p_k)^2 - p_k^2$, where $p_1 = p_1^*$ and $p_2 = 0$).

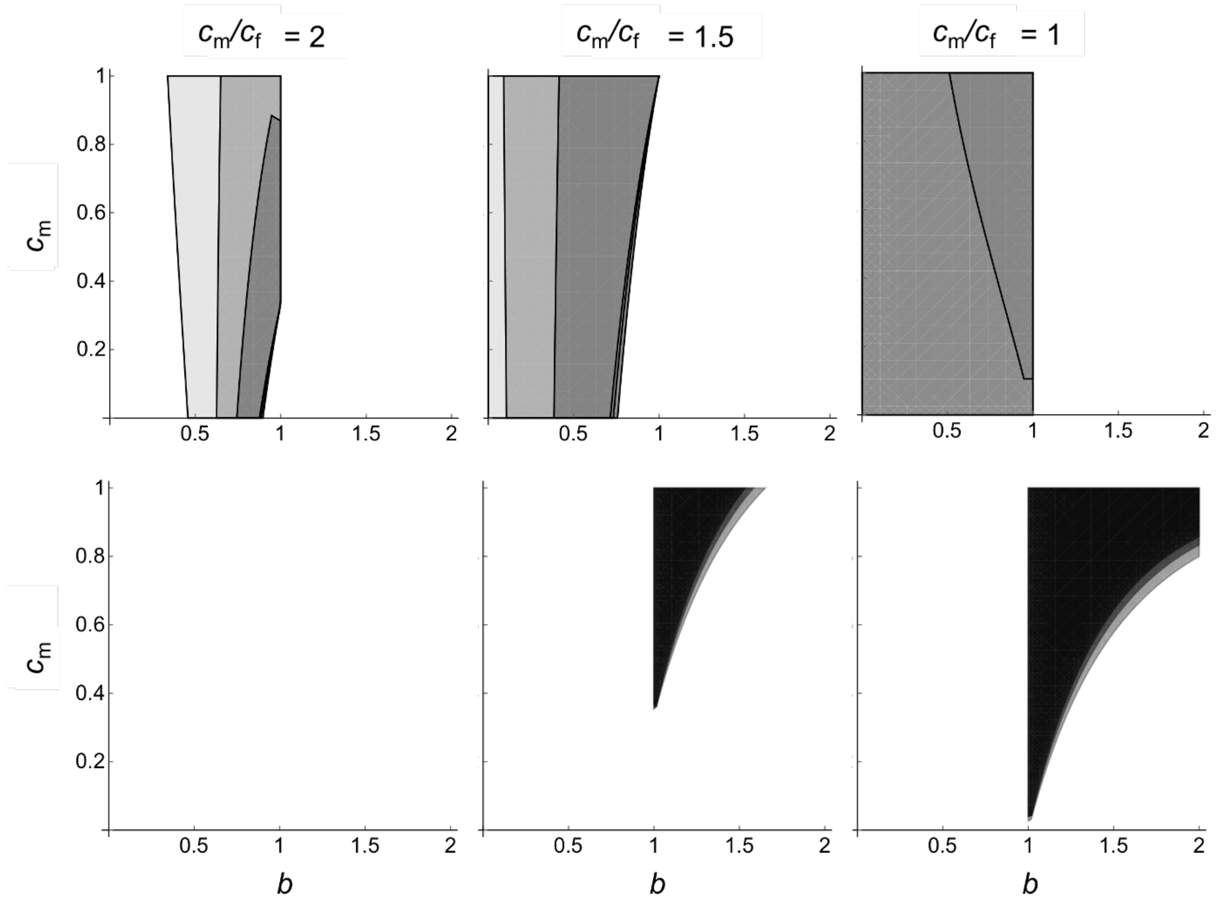


**Appendix Figure 2: Conditions for polymorphism in a two-locus trait** ($L = 2$) **allowing for unequal phenotypic effects.** Panels show parameter conditions for one-locus (top row panels) and two-locus (bottom row panels) polymorphisms from our di-allelic models with two loci ($L = 2$) when male and female fecundity follow power functions (eq. I.A, Box I, see Appendix B.2 for analysis). Panels show polymorphism space as a function of equal selection and similarly shaped fecundity landscapes across the sexes ($c = c_m = c_f$ and $b = b_m = b_f$). In all panels, darkest regions show the case of equal effects at both loci ($\delta_1 = \delta_2 = \delta$) and lighter regions show cases of unequal effects ($\delta_2 = \rho\delta_1$), where medium shade shows represents $\rho = 0.5$ and the lightest shade represents $\rho = 0.1$. Other parameters: $\delta = \delta_1 = 1, c_m = 0.1$.

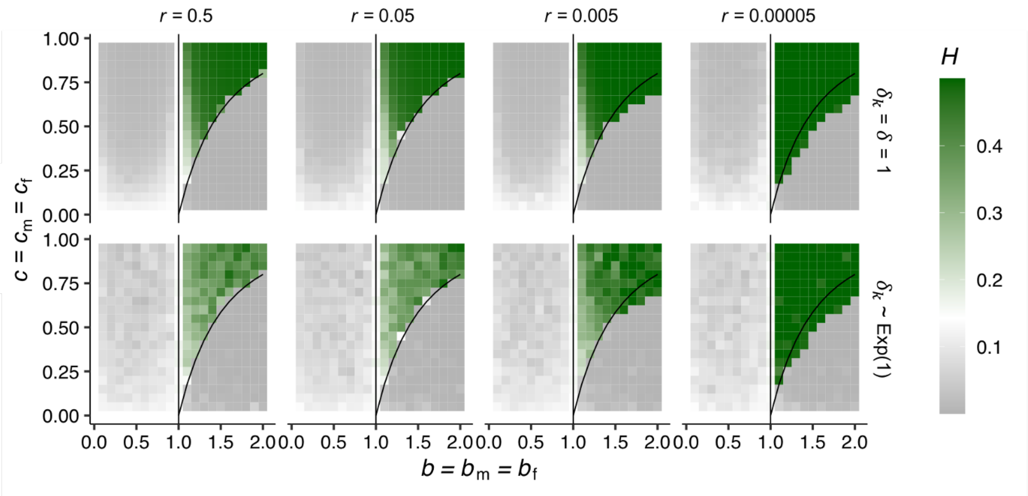


**Appendix Figure 3: Average heterozygosity in a polygenic trait for different effect sizes and different strengths of recombination.** Panels show heterozygosity in simulations of our polygenic di-allelic model when male and female fecundity follow power functions (eq. I.A, Box I, Appendices B.3, C.1 and C.2 for details, $L = 10, \mu = 10^{-5}$ for all panels). White, green, and grey squared show average heterozygosity values that are respectively equal, higher, and lower to that expected at a neutral locus at mutation-drift balance. Black lines show the conditions for diversifying selection calculated in the continuum-of-alleles model (eq. A-19). Panels in the top row show results when all alleles have equal effect size ($\delta_k = \delta = 1$ for all $k \in \{1, \ldots 10\}$), while the bottom row allows for unequal effects (where $\delta_k$ is drawn from an exponential distribution with mean $\delta = 1$). Columns show results for different strengths of genetic linkage between loci, $r$.

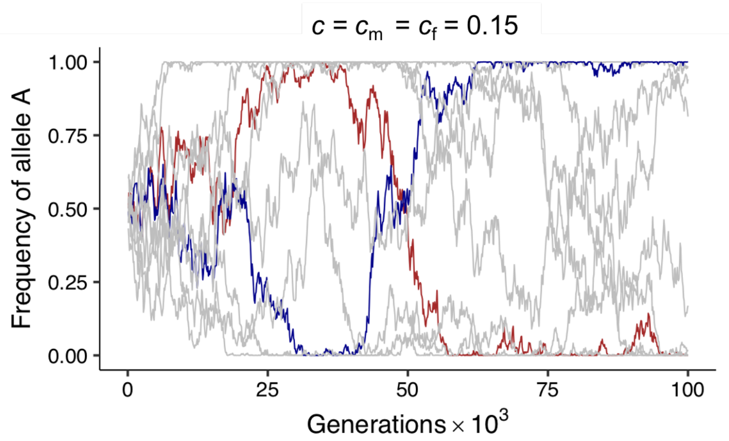


**Appendix Figure 4: Allele frequencies through time under stabilising selection with different effect sizes.** Curves show frequency of the female-beneficial allele $A_k$ at each locus $k$ through time in a simulation of our polygenic di-allelic model when male and female fecundity follow power functions (eq. I.A, Box I, Appendices B.3, C.1 for details, $L = 10, b = 0.5, \mu = 10^{-5}$). Coloured curves show an example of two loci that show transient genetic polymorphism. Effect size at each locus, $\delta_k$, is drawn from an exponential distribution with mean $\delta = 1$.

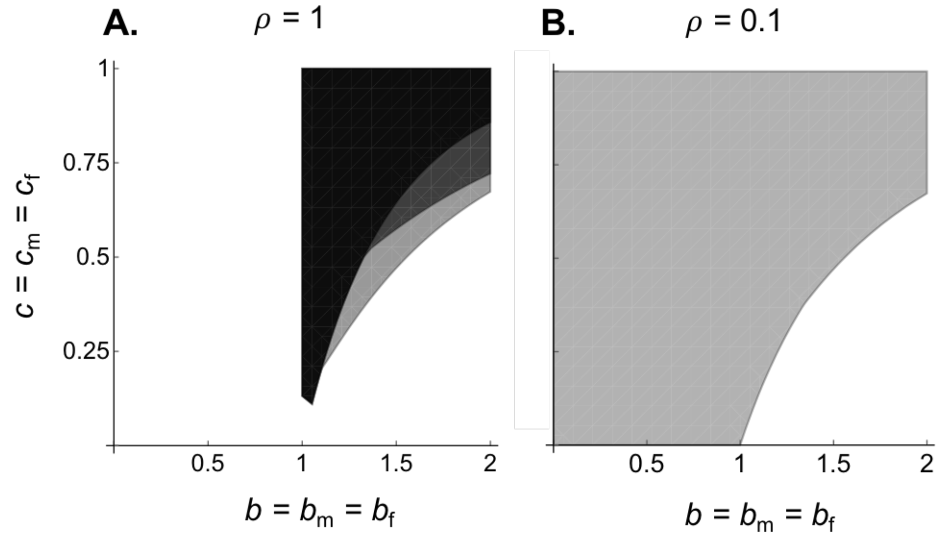


**Appendix Figure 5: Polymorphism conditions when $L = 2$ allowing for genetic linkage.** Panels show parameter conditions two-locus polymorphisms from our di-allelic models with two loci ($L = 2$) when male and female fecundity follow symmetrical power functions ($c = c_{\mathrm{m}} = c_{\mathrm{f}}$ and $b = b_{\mathrm{m}} = b_{\mathrm{f}}$ in eq. I.A, Box I, see Appendix B.2, C.1 and C.2 for details). Panel **A** shows conditions for equal effects across loci ($\delta_1 = \delta_2 = 1$) and Panel **B** shows an example of unequal effects ($\rho = 0.1$ where $\delta_2 = \rho\delta_1$ and $\delta_1 = 1$). Different shaded regions correspond to different rates of recombination between loci, black to $r = 0.5$, grey to $r = 0.05$, and light grey to $r = 0.005$.

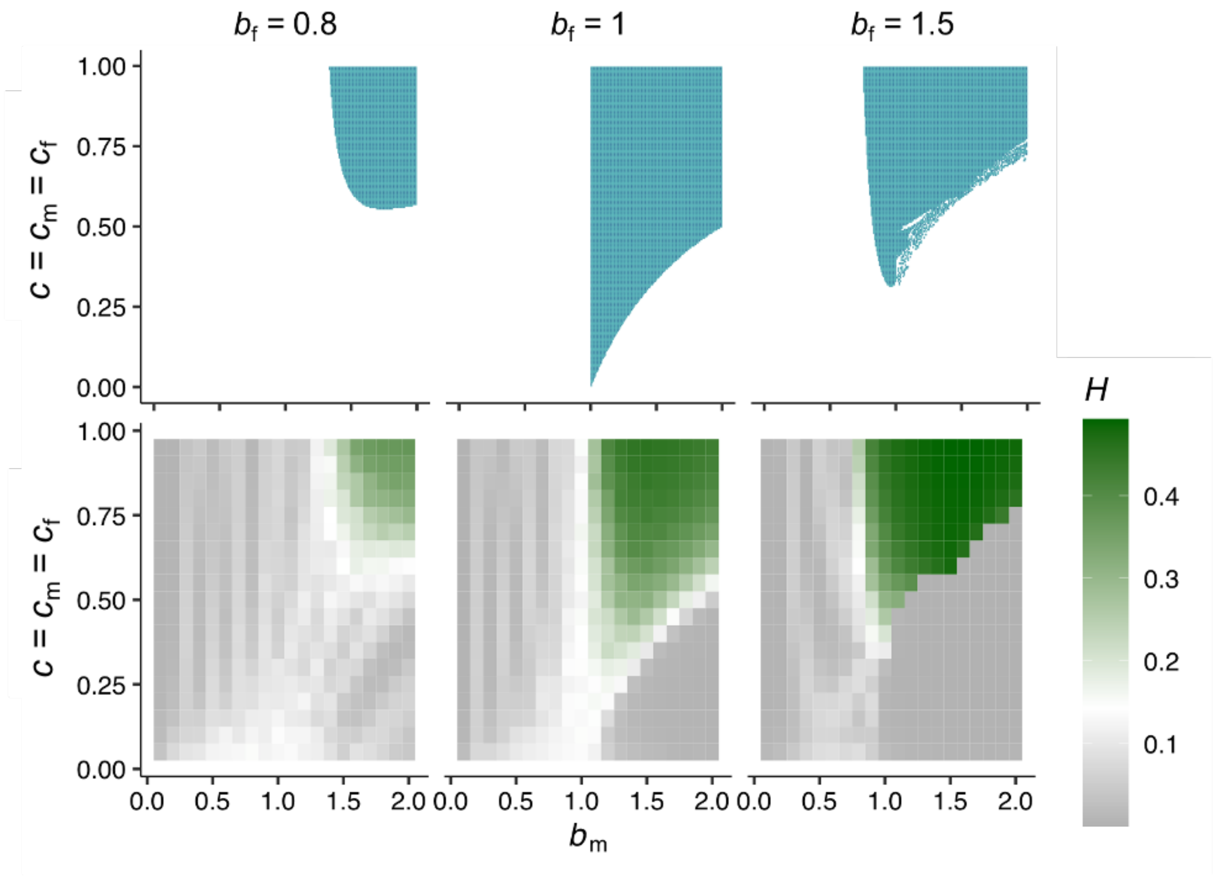


**Appendix Figure 6: Conditions for elevated genetic variation when the curvature of the fecundity curves is allowed to differ between the sexes.** Panels show mathematical and simulation results for the continuum-of-alleles and di-allelic polygenic model when male and female fecundity follow power functions that can show different curvature ($b_f \neq b_m$, eq. I.A, Box I, see Appendix C.3 for details). Top row panels show the conditions for diversifying selection (green regions) in the continuum-of-alleles model depending on the fecundity curve in each sex ($b_m$, $b_f$ and the strength of fecundity costs $c = c_m = c_f$). Bottom row panels show average heterozygosity from polygenic simulations of 10 di-allelic loci ($L = 10$) with $\delta = 1, \mu = 10^{-5}$.

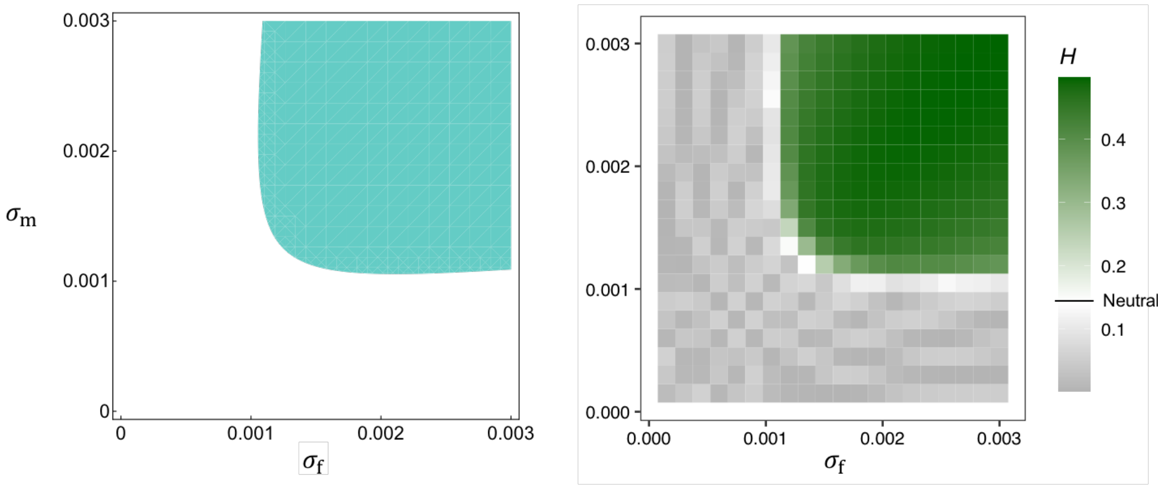


**Appendix Figure 7: Conditions for diversifying selection and heterozygosity with Gaussian fecundity curves.** Panels show mathematical and simulation results for the continuum-of-alleles and di-allelic polygenic model when male and female fecundity follow Gaussian functions (eq. C-2, Appendix C.4 for details). Left-hand panel shows conditions (green regions) for diversifying selection to arise in the continuum-of-alleles model with Gaussian fecundity curves (i.e., to satisfy eq. C-6). Right-hand panel shows heterozygosity ($H$) across loci from individual-based simulations of the di-allelic with Gaussian fecundity (Appendix C.4 for simulation details, $L = 10, \theta = 20, \delta = 1, \mu = 10^{-5}$).

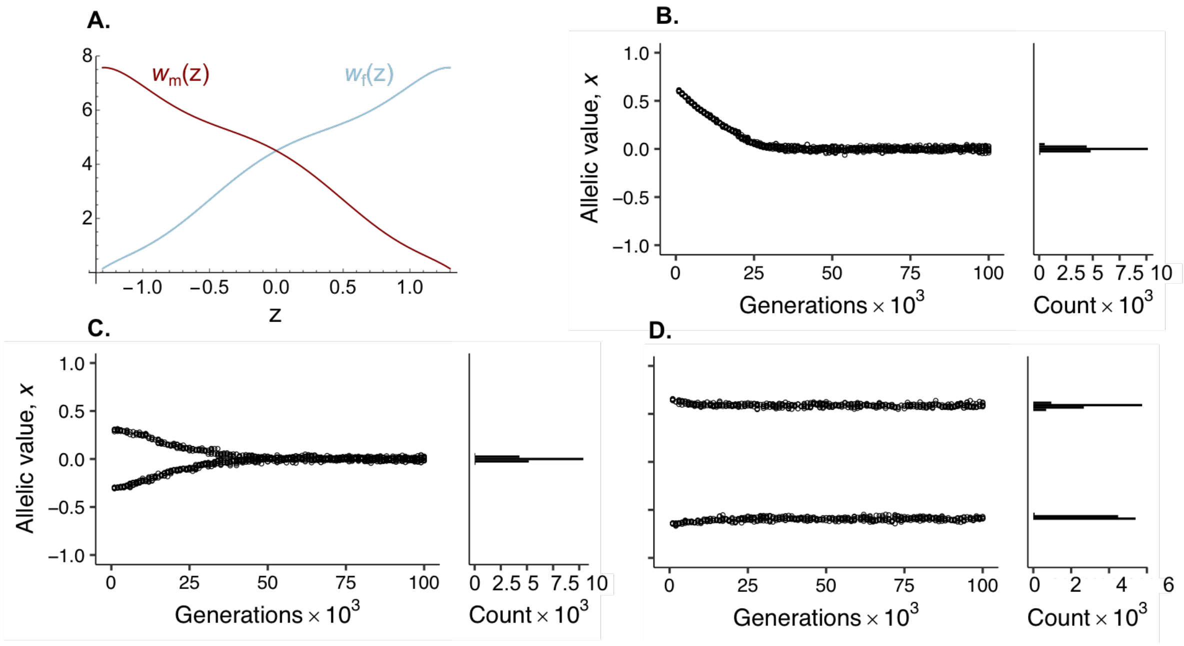


**Appendix Figure 8: Continuum-of-allele model with alternative fitness functions.** Panel **A** shows the male and female fecundity functions $w_\mathrm{m}(z) = \alpha_0 - \alpha_1 z + \alpha_2 z^2 - \alpha_3 z^3 + \alpha_4 z^4 - \alpha_5 z^5 + \alpha_6 z^6$ (red) and $w_\mathrm{f}(z) = \alpha_0 + \alpha_1 z + \alpha_2 z^2 + \alpha_3 z^3 + \alpha_4 z^4 + \alpha_5 z^5 + \alpha_6 z^6$ (blue) with $\alpha_0 = 4.5, \alpha_1 = 2.7, \alpha_2 = -2.1, \alpha_3 = 0.6, \alpha_4 = 2.2, \alpha_5 = -0.3, \alpha_6 = -0.7$, which satisfy the criterion (C-20) when selection is weak. For these functions, our mathematical analysis in Appendix C.5 indicates there is a stable monomorphic equilibrium at $x^* = 0$ and a stable dimorphic equilibrium at $(-x^*, x^*) = (-0.59, 0.59)$ (note that variation in $w_\mathrm{m}(z)$ and $w_\mathrm{f}(z)$ as given here is large; we checked through a numerical analysis that these functions entailed the existence of an uninvadable stable coalition at $(-x^*, x^*) = (-0.59, 0.59)$ under strong selection, which was also checked with simulations as shown in Panels B-D). Panels **B**-**D** show simulation results from the continuum-of-alleles model with small effect mutations (see Appendix A.4.3 for details) with fecundity curves shown in Panel **A**. In Panel **B**, simulations were initialised with a single allelic value at $x = 0.59$ (so that the population begins in a monomorphic state). Meanwhile, in Panels **C**-**D** populations are initialised with two alleles at equal frequencies: a negative allele $-x$ and a positive allele $x$ (with $x = 0.3$ for Panel **C** and $x = 0.59$ for Panel **D**). In line with our mathematical analysis, we found it was possible to maintain a stable two-allele polymorphism but only if the population was initialised in a dimorphic state with alleles that were sufficiently diverged from each other (i.e., with large enough $x$).

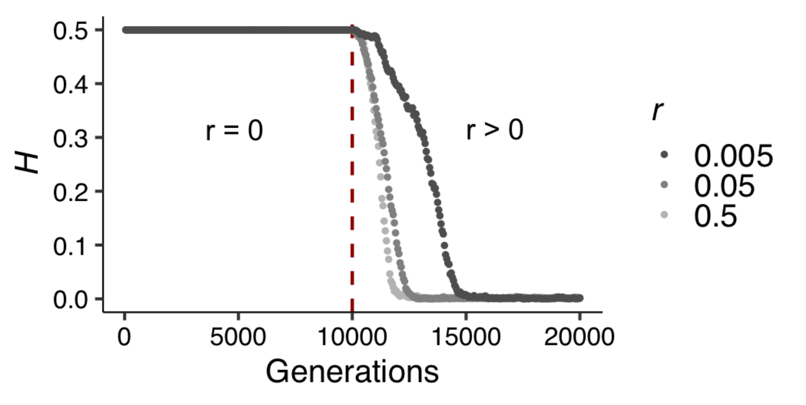


**Appendix Figure 9: Heterozygosity across loci in a polygenic trait** ($L = 10$) **with alternative fitness functions.** Plot shows average heterozygosity across loci through time in individual-based simulations of the di-allelic model with fecundity functions given in Appendix Fig. 8A (see legend and Appendix C.5.2 for simulation details). Simulations begin with a first phase where there is no recombination among loci ($r = 0$), followed by a second phase beginning at generation 10,000 (dashed red line) where recombination occurs. Different shades of gray refer to different strengths of recombination in the second phase.

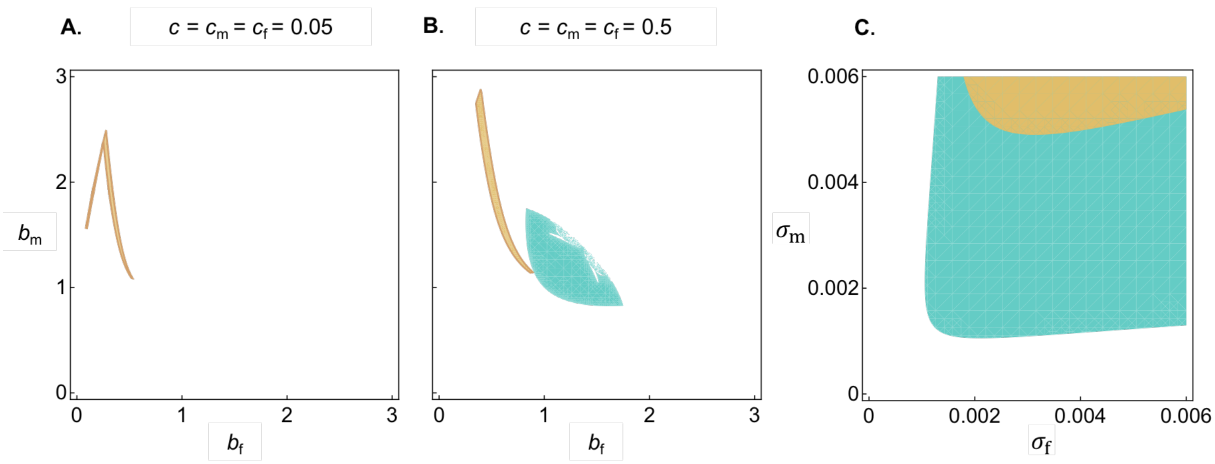


**Appendix Figure 10: Conditions for diversifying selection at X-linked and autosomal loci.** Coloured regions represent parameter values leading to diversifying selection at X-linked (gold regions, see inequality D-20) and autosomal (green regions, inequality eq. D-19) loci under the continuum-of-alleles model. Panels **A** and **B** show regions calculated from power fecundity functions (eq. I.A, Box I) under different strengths of sexual antagonism ($c = c_m = c_f$, with $c = 0.05$ in panel **A** and $c = 0.5$ in panel **B**), Panel **C** shows regions calculated from Gaussian fecundity functions (eq. C-2, Appendix C.4) where $\theta = 20$ (as assumed in polygenic individual-based simulations, Appendix C.4 for details). Note that the values for parameters $\sigma_{\rm m}$ and $\sigma_{\rm f}$ used in this panel correspond to extremely strong sexual conflict (e.g., $\sigma_{\rm m} = 0.002$ leads to a decrease in fecundity two expressing the female optimum of $1 - w_{\rm m}(\theta) \approx 0.96$), so that conditions for polymorphism at both X-linked and autosomal loci are restrictive.

#### References


[1] Geritz SA, Kisdi É. 2000 Adaptive dynamics in diploid, sexual populations and the evolution of reproductive isolation. *Proceedings of the Royal Society of London. Series B: Biological Sciences* **267**, 1671–1678.

[2] Dercole F, Rinaldi S. 2008 *Analysis of evolutionary processes*. Princeton University Press.

[3] Lande R, Arnold SJ. 1983 The measurement of selection on correlated characters. *Evolution* pp. 1210–1226.

[4] Parker GA, Smith JM. 1990 Optimality theory in evolutionary biology. *Nature* **348**, 27–33.

[5] Geritz SA, Kisdi E, Metz JA et al.. 1998 Evolutionarily singular strategies and the adaptive growth and branching of the evolutionary tree. *Evolutionary ecology* **12**, 35–57.

[6] Rueffler C, Van Dooren TJ, Leimar O, Abrams PA. 2006 Disruptive selection and then what?. *Trends in Ecology & Evolution* **21**, 238–245.

[7] Haller BC, Messer PW. 2023 SLiM 4: multispecies eco-evolutionary modeling. *The American Naturalist* **201**, E127–E139.

[8] Kidwell J, Clegg M, Stewart F, Prout T. 1977 Regions of stable equilibria for models of differential selection in the two sexes under random mating. *Genetics* **85**, 171–183.

[9] Fry JD. 2010 The genomic location of sexually antagonistic variation: some cautionary comments. *Evolution: International Journal of Organic Evolution* **64**, 1510–1516.

[10] Patten MM, Haig D, Ubeda F. 2010 Fitness variation due to sexual antagonism and linkage disequilibrium. *Evolution* **64**, 3638–3642.

[11] Connallon T, Clark AG. 2010 Sex linkage, sex-specific selection, and the role of recombination in the evolution of sexually dimorphic gene expression. *Evolution: International Journal of Organic Evolution* **64**, 3417–3442.

[12] Ubeda F, Haig D, Patten MM. 2011 Stable linkage disequilibrium owing to sexual antagonism. *Proceedings of the Royal Society B: Biological Sciences* **278**, 855–862.

[13] Otto SP. 2019 Evolutionary potential for genomic islands of sexual divergence on recombining sex chromosomes. *New Phytologist* **224**, 1241–1251.

[14] Walsh B, Lynch M. 2018 *Evolution and selection of quantitative traits*. Oxford University Press.

[15] Gavrilets S, Hastings A. 1993 Maintenance of genetic variability under strong stabilizing selection: a two-locus model.. *Genetics* **134**, 377–386.

[16] Bürger R, Gimelfarb A. 1999 Genetic variation maintained in multilocus models of additive quantitative traits under stabilizing selection. *Genetics* **152**, 807–820.

[17] Bürger R, Gimelfarb A. 2004 The effects of intraspecific competition and stabilizing selection on a polygenic trait. *Genetics* **167**, 1425–1443.

[18] Lande R. 1980 Sexual dimorphism, sexual selection, and adaptation in polygenic characters. *Evolution* **34**, 292–305.

[19] Connallon T, Clark AG. 2014a Balancing selection in species with separate sexes: insights from Fisher's geometric model. *Genetics* **197**, 991–1006.

[20] Connallon T, Clark AG. 2014b Evolutionary inevitability of sexual antagonism. *Proceedings of the Royal Society B: Biological Sciences* **281**, 20132123.

[21] Muralidhar P, Coop G. 2023 Polygenic outcomes of sexually antagonistic selection. *bioRxiv* pp. 2023–03.

[22] Levene H. 1953 Genetic Equilibrium When More Than One Ecological Niche is Available. *https://doi.org/10.1086/281792* **87**, 331–333.

[23] Christiansen FB. 1975 Hard and soft selection in a subdivided population. *The American Naturalist* **109**, 11–16.

[24] Connallon T, Débarre F, Li XY. 2018 Linking local adaptation with the evolution of sex differences. *Philosophical Transactions of the Royal Society B: Biological Sciences* **373**.

[25] Turelli M, Barton N. 2004 Polygenic variation maintained by balancing selection: pleiotropy, sex-dependent allelic effects and G× E interactions. *Genetics* **166**, 1053–1079.

[26] Felsenstein J. 1979 Excursions along the interface between disruptive and stabilizing selection. *Genetics* **93**, 773–795.

[27] Curtsinger JW, Service PM, Prout T. 1994 Antagonistic pleiotropy, reversal of dominance, and genetic polymorphism. *The American Naturalist* **144**, 210–228.

[28] Rice WR. 1984 Sex chromosomes and the evolution of sexual dimorphism. *Evolution* **38**, 735–742.

[29] Otto SP, Pannell JR, Peichel CL, Ashman TL, Charlesworth D, Chippindale AK, Delph LF, Guerrero RF, Scarpino SV, McAllister BF. 2011 About PAR: the distinct evolutionary dynamics of the pseudoautosomal region. *Trends in Genetics* **27**, 358–367.

[30] Eshel I. 1983 Evolutionary and continuous stability. *Journal of theoretical Biology* **103**, 99–111.

[31] Avila P, Mullon C. 2023 Evolutionary game theory and the adaptive dynamics approach: adaptation where individuals interact. *Philosophical Transactions of the Royal Society B* **378**, 20210502.

[32] Cheng C, Kirkpatrick M. 2016 Sex-specific selection and sex-biased gene expression in humans and flies. *PLoS Genetics* **12**.

[33] Ruzicka F, Dutoit L, Czuppon P, Jordan CY, Li XY, Olito C, Runemark A, Svensson EI, Yazdi HP, Connallon T. 2020 The search for sexually antagonistic genes: Practical insights from studies of local adaptation and statistical genomics. *Evolution letters* **4**, 398–415.

[34] Ruzicka F, Holman L, Connallon T. 2022 Polygenic signals of sex differences in selection in humans from the UK Biobank. *PLoS Biology* **20**, e3001768.

[35] Singh A, Punzalan D. 2018 The strength of sex-specific selection in the wild. *Evolution* **72**, 2818–2824.

[36] Sanjak JS, Sidorenko J, Robinson MR, Thornton KR, Visscher PM. 2018 Evidence of directional and stabilizing selection in contemporary humans. *Proceedings of the National Academy of Sciences* **115**, 151–156.

[37] Lynch M, Walsh B et al.. 1998 *Genetics and analysis of quantitative traits* vol. 1. Sinauer Sunderland, MA.

[38] Mackay TF, Stone EA, Ayroles JF. 2009 The genetics of quantitative traits: challenges and prospects. *Nature Reviews Genetics* **10**, 565–577.

[39] Rowe L, Houle D. 1996 The lek paradox and the capture of genetic variance by condition dependent traits. *Proceedings of the Royal Society of London. Series B: Biological Sciences* **263**, 1415–1421.

[40] Jennions MD, Moller AP, Petrie M. 2001 Sexually selected traits and adult survival: a meta-analysis. *The Quarterly Review of Biology* **76**, 3–36.

[41] Cornwallis CK, Uller T. 2010 Towards an evolutionary ecology of sexual traits. *Trends in Ecology & Evolution* **25**, 145–152.

[42] Dougherty LR. 2021 Meta-analysis reveals that animal sexual signalling behaviour is honest and resource based. *Nature Ecology & Evolution* **5**, 688–699.

[43] Bonduriansky R, Chenoweth SF. 2009 Intralocus sexual conflict. *Trends in ecology & evolution* **24**, 280–288.

[44] Poissant J, Wilson AJ, Coltman DW. 2010 Sex-specific genetic variance and the evolution of sexual dimorphism: a systematic review of cross-sex genetic correlations. *Evolution* **64**, 97–107.

[45] Mank JE. 2017 Population genetics of sexual conflict in the genomic era. *Nature Reviews Genetics* **18**, 721–730.

[46] Zhu C, Ming MJ, Cole JM, Edge MD, Kirkpatrick M, Harpak A. 2023 Amplification is the primary mode of gene-by-sex interaction in complex human traits. *Cell Genomics* **3**.

[47] Charlesworth B, Charlesworth D. 2010 *Elements of Evolutionary Genetics*. Roberts and Company Publishers.

[48] Phillips PC. 2008 Epistasis—the essential role of gene interactions in the structure and evolution of genetic systems. *Nature Reviews Genetics* **9**, 855–867.

[49] Huang W, Richards S, Carbone MA, Zhu D, Anholt RR, Ayroles JF, Duncan L, Jordan KW, Lawrence F, Magwire MM et al.. 2012 Epistasis dominates the genetic architecture of Drosophila quantitative traits. *Proceedings of the National Academy of Sciences* **109**, 15553–15559.

[50] Gimelfarb A. 1989 Genotypic variation for a quantitative character maintained under stabilizing selection without mutations: epistasis.. *Genetics* **123**, 217–227.

[51] Hermisson J, Hansen TF, Wagner GP. 2003 Epistasis in polygenic traits and the evolution of genetic architecture under stabilizing selection. *The American Naturalist* **161**, 708–734.

[52] Turelli M, Barton N. 2006 Will population bottlenecks and multilocus epistasis increase additive genetic variance?. *Evolution* **60**, 1763–1776.

[53] Arnqvist G, Vellnow N, Rowe L. 2014 The effect of epistasis on sexually antagonistic genetic variation. *Proceedings of the Royal Society B: Biological Sciences* **281**, 20140489.

[54] Arnqvist G. 2011 Assortative mating by fitness and sexually antagonistic genetic variation. *Evolution* **65**, 2111–2116.

[55] Tazzyman SJ, Abbott JK. 2015 Self-fertilization and inbreeding limit the scope for sexually antagonistic polymorphism. *Journal of Evolutionary Biology* **28**, 723–729.

[56] Kasimatis KR, Ralph PL, Phillips PC. 2019 Limits to genomic divergence under sexually antagonistic selection. *G3: Genes, Genomes, Genetics* **9**, 3813–3824.

[57] Flintham EO, Savolainen V, Mullon C. 2021 Dispersal alters the nature and scope of sexually antagonistic variation. *The American Naturalist* **197**, 543–559.

[58] Débarre F, Gandon S. 2011 Evolution in heterogeneous environments: between soft and hard selection. *The American Naturalist* **177**, E84–E97.

[59] Meszéna G, Czibula I, Geritz SAH. 1997 Adaptive dynamics in a 2-patch environment: A toy model for allopatric and parapatric specation. *Journal of Biological Systems* **5**, 265–284.

[60] Lythgoe KA. 1997 Consequences of gene flow in spatially structured populations. *Genetics Research* **69**, 49–60.

[61] Spichtig M, Kawecki TJ. 2004 The maintenance (or not) of polygenic variation by soft selection in heterogeneous environments. *The American Naturalist* **164**, 70–84.

[62] Doorn GSV, Dieckmann U. 2006 The long-term evolution of multilocus traits under frequency-dependent disruptive selection. *Evolution* **60**, 2226–2238.

[63] Svardal H, Rueffler C, Hermisson J. 2015 A general condition for adaptive genetic polymorphism in temporally and spatially heterogeneous environments. *Theoretical Population Biology* **99**, 76–97.

[64] Schmid M, Rueffler C, Lehmann L, Mullon C. 2024 Resource Variation Within and Between Patches: Where Exploitation Competition, Local Adaptation, and Kin Selection Meet. *The American Naturalist* **203**, E19–E34.

[65] Turelli M, Barton NH. 1994 Genetic and statistical analyses of strong selection on polygenic traits: what, me normal?. *Genetics* **138**, 913–941.

[66] Bulmer M. 1971 The effect of selection on genetic variability. *The American Naturalist* **105**, 201–211.

[67] Bürger R, Gimelfarb A. 2002 Fluctuating environments and the role of mutation in maintaining quantitative genetic variation. *Genetics Research* **80**, 31–46.

[68] Wright S. 1935 Evolution in populations in approximate equilibrium. *Journal of Genetics* **30**, 257–266.

[69] Robertson A. 1956 The effect of selection against extreme deviants based on deviation or on homozygosis: With Two Text-figures. *Journal of Genetics* **54**, 236–248.

[70] Barton NH. 1986 The maintenance of polygenic variation through a balance between mutation and stabilizing selection. *Genetics Research* **47**, 209–216.

[71] Gavrilets S, Hastings A. 1994 Dynamics of genetic variability in two-locus models of stabilizing selection.. *Genetics* **138**, 519–532.

[72] Lande R. 1976 Natural selection and random genetic drift in phenotypic evolution. *Evolution* pp. 314–334.

[73] Connallon T, Clark AG. 2012 A general population genetic framework for antagonistic selection that accounts for demography and recurrent mutation. *Genetics* **190**, 1477–1489.

[74] Mullon C, Pomiankowski A, Reuter M. 2012 The effects of selection and genetic drift on the genomic distribution of sexually antagonistic alleles. *Evolution: International Journal of Organic Evolution* **66**, 3743–3753.
